# The cause of on-target point mutations generated by CRISPR-Cas9 treatment in the yeast *Xanthophyllomyces dendrorhous*

**DOI:** 10.1101/2021.08.31.458371

**Authors:** Jixuan Hong, Ziyue Meng, Zixi Zhang, Hang Su, Yuxuan Fan, Ruilin Huang, Ruirui Ding, Ning Zhang, Fuli Li, Shi’an Wang

**Affiliations:** Shandong Provincial Key Laboratory of Synthetic Biology, Key Laboratory of Biofuels, Qingdao Institute of Bioenergy and Bioprocess Technology, Chinese Academy of Sciences, Qingdao 266101, China; Shandong Energy Institute, Qingdao 266101, China; University of Chinese Academy of Sciences, Beijing 100039, China

## Abstract

Recognizing outcomes of DNA repair induced by CRISPR-Cas9 cutting is vital for precise genome editing. Reported DNA repair outcomes after Cas9 cutting include deletions/insertions and low frequency of genomic rearrangements and nucleotide substitutions. Thus far, substitution mutations caused by CRISPR-Cas9 has not attracted much attention. Here, we identified on-target point mutations induced by CRISPR-Cas9 treatment in the yeast *Xanthophyllomyces dendrorhous* by Sanger and Illumina sequencing. Different from previous studies, our findings suggested that the on-target mutations are not random and they cannot render the gRNA effective. Moreover, these point mutations showed strong sequence dependence that is not consistent with the observations in Hela cells, in which CRISPR-mediated substitutions were considered lacking sequence dependence and conversion preferences. Furthermore, this study demonstrated that the NHEJ components *Ku70, Ku80, Mre11*, or *RAD50*, and the overlapping roles of non-essential DNA polymerases were necessary for the emergence of point mutations, increasing the knowledge on CRISPR-Cas9 mediated DNA repair.

## INTRODUCTION

CRISPR-Cas9 genome editing relies on sequence-specific double-strand DNA cleavage by Cas9 followed by DNA repair (1). Double-strand DNA breaks (DSBs) from Cas9 cutting can be repaired by non-homologous end joining (NHEJ), microhomology-mediated end joining (MMEJ), and homology-directed repair (HDR) (2). NHEJ and MMEJ often result in nucleotide deletion or insertion (3-6). Although accurate genetic modifications can be introduced by HDR using exogenous donor DNA, the recombination efficiency is relatively lower than that of NHEJ in mammalian cells (7). A variety of strategies have been proposed to improve precise gene editing, such as modification of host repair pathways (8-10), use of Cas9 and gRNA mutants (11), providing single-stranded DNA donors (12), and computation prediction of mutations (6, 13, 14). Most of these methods were developed to address previously identified repair outcomes and mechanisms.

Spontaneous repair of CRISPR-Cas9 mediated DNA cleavage usually leads to deletion and insertion (3-6). A low frequency of complex genomic rearrangements coming from the repair of CRISPR-Cas9-mediated DNA cleavage were also observed in mouse and human cell lines (15). In addition, single nucleotide substitutions around DNA cleavage sites after CRISPR-Cas9 treatment were reported in a few studies (4, 16, 17). Such substitution mutations caused by CRISPR-Cas9 are typically attributed to ineffective gRNA and spontaneous mutations, and this phenomenon has not attracted serious attention (4, 18). Recently, Hwang et al. developed a CRISPR-Sub method to statistically detect apparent substitutions in high-throughput sequencing data and proposed that DNA ligase IV, an essential component of the NHEJ repair pathway, is closely related with the nucleotide substitutions (18). However, the characteristics and the cause of nucleotide substitutions remain to be investigated.

When using CRISPR-Cas9 to edit genes in an astaxanthin-producing yeast *Xanthophyllomyces dendrorhous* (19), we identified on-target nucleotide substitutions. This phenomenon motivated us to comprehensively characterize DNA repair outcomes in CRISPR-Cas9 genome editing using this yeast. In this study, we confirmed that the repair of CRISPR-Cas9-mediated DNA cleavage do lead to on-target point mutations. Furthermore, our data suggested that the on-target point mutations induced by the repair of CRISPR-Cas9 cleavage requires non-homologous end joining complex and non-essential DNA polymerases.

## MATERIAL AND METHODS

### Yeast strains and manipulations

*X. dendrorhous* strains used in this study are described in Supplementary Table S2. For standard growth, the yeast strains were grown with rotary shaking at 250 rpm and 22 °C in liquid YPD medium (1% yeast extract, 2% peptone, 2% glucose). Transformation of *X. dendrorhous* yeasts was performed using an electroporation procedure as previously described (20). Briefly, a single yeast colony growing on the YPD plate was inoculated into 40 ml of YPD broth in a 250 mL flask and incubated for 48 h at 22°C and 250 rpm. The pre-cultured cells were inoculated into 50 ml of YPD broth in a 500 mL flask to an optical density (OD_600_) of 0.02 and incubate at 22°C and 250 rpm overnight until an OD_600_ of approximate 1.2. Then, cultures were centrifuged and the cell pellet was suspended in 25 mL of potassium phosphate buffer containing 25 mM freshly-made DTT and incubated for 15 min at room temperature. Next, the cells were washed twice using 25 mL of ice-cold STM buffer (270 mM sucrose, 10 mM Tris–HCl pH 7.5, 1 mM MgCl_2_) and resuspended in STM buffer to approximate 3 × 10^9^ cells/mL. DNA fragments were acquired by *Sma*I cutting, mixed with 60 μl of the competent cells on ice, and transferred into 0.2 cm cuvettes and then electroporated at 1000 Ω, 800 V, and 25 μF using BTX ECM830 Electroporator (Genetronics, San Diego, CA). Transformants were selected on YPD plate containing 200 μg/ml of G418 or 100 μg/ml of hygromycin, or 200 μg/ml of Zeocin.

### Plasmid construction

Plasmids and oligonucleotide primers used in this study are described in Supplementary Data S1. The Gibson assembly method was used to construct plasmids. DNA insertion fragments and backbones were amplified using the KAPA HiFi HotStart PCR kit (Kapa Biosystems, Boston, USA), and they were assembled at 50 °C for 1 h.

The Cas9 gene from the plasmid pCAS1yl (21) was codon-optimized and synthesized. An eGFP gene was employed to confirm the function of an SV40 nuclear localization sequence (NLS) that was codon-optimized according to the codon usage of *X. dendrorhous*. The SV40 NLS sequences were fused to both the C-terminal and the N-terminal of the codon-optimized Cas9 gene. The eGFP or the Cas9 gene was placed under the control of an ADH4 promoter. Expression of the sgRNA cassette was driven by a fusion promoter SCR1-tRNA^Ala^ (Data S2). The SCR1 and tRNA sequences were obtained by BLASTN search using the query sequences from *Saccharomyces cerevisiae* and *Yarrowia lipolytic*a, respectively. Structural analysis of the predicted tRNA-Ala sequence was retrieved by the tRNAscan-SE 2.0 software using ViennaRNA Package 2.0 algorithms (22). The gRNA targeting sequences were designed using the online tool CRISPy-web (23). Scaffold_69 (GenBank No. LN483157.1) and Scaffold_79 (GenBank No. LN483167.1) of the *X. dendrorhous* genome assembly were scanned. The DNA regions of geranylgeranyl pyrophosphate synthase gene *CrtE* (LN483167.1: 1539235 to 1541616) and astaxanthin synthase gene *CrtS* (LN483157.1: 1230024 to 1233189) were selected as the CRISPR-Cas9 targets. For ease of generating detectable phenotypes, the protospacers were designed in or near the active centers of CrtE, the structure of which was predicted by structure homology modeling using SWISS-MODEL server (24) and the human ortholog 2Q80 (PDB ID) as a template (25). The predicted structures were visualized using the program PyMOL version 1.7.0.0. Target sequences were fused to the upstream of the structural gRNA. The Cas9 and sgRNA sequences were assembled in the same plasmid by Gibson method. For homologous recombination of *Cas9* and sgRNA in *X. dendrorhous* genome, rDNA fragments were flanked to the CRISPR-Cas9 expression cassettes containing a geneticin resistant gene as the selective marker.

### Western blotting

The protein extraction protocol was modified according to a previous publication (26). Yeast strains were cultured in the YPD broth at 22°C and 250 rpm until the OD_600_ reached 2 to 4. The cells were harvested from 50 ml culture by centrifugation at 5,000 × g for 5 min at 4°C. The pellet was resuspended in a protein lysis buffer containing 100 mM NaHCO_3_, 0.5% Triton 100, cocktail protease inhibitors (Roche, Mannheim, Germany), and pH 8.5. Cells were disrupted for six times in a RiboLyzer for 20 s at 4.0 m/s and chilled on ice for 1 min between vortexing steps. The cell debris was removed by centrifugation at 14,000 rpm for 30 min at 4°C, and the supernatant was transferred to 1.5 ml tubes. The protein concentration was measured by Coomassie Brilliant Blue assay using bovine serum albumin (Sigma) as standard. Western detection was performed using the Cas9 (7A9-3A3) mouse mAb antibody (Cell Signaling Technology, Inc. MA, USA) and goat anti-rabbit antibody conjugated to horse radish peroxidase (Agrisera, Sweden). Images of the blots were exposed on films (Kodak, X-OMAT BT, USA).

### Gene editing

The CRISPR-Cas9 plasmids containing both Cas9 and sgRNA sequences were cut by *Sma*I and the expression cassettes were inserted into the rRNA gene loci by homologous recombination. Homologous recombination is the predominant mechanism in repair of DNA lesion in the yeast *X. dendrorhous*. In the gene editing procedure, no donor homologous DNA was supplied and thus the edited clones had been repaired by inherent cellular activities. The pigment-changed edited clones growing on YPD plates were picked up and used in characterizing the repair outcomes induced by CRISPR-Cas9 cleavage (Supplementary Data S3). Like in *S. cerevisiae* (27), the percent of edited clones was below 0.1% in *X. dendrorhous* seeing that no donor DNA had been supplied.

### Characterizing repair patterns by PCR

To characterize the repair outcome of edited clones, PCR was initially used. The DNA region including the CRISPR-Cas9 target sites was amplified by a variety of primer pairs (Supplementary Figure S2, S3). The amplicons were sequenced using conventional Sanger sequencing technology. For the samples that DNA regions spanning the target sites were not successfully amplified from them, both an upstream and a downstream DNA region of the break sites were amplified to evaluate if chromosome rearrangement happened in them. Additionally, genome sequencing and chromosome karyotype detection were used in characterizing DNA repair patterns induced by CIRISPR-Cas9 cleavage.

### Illumina genome sequencing and bioinformatic analyses

The genome sequences of representative edited clones were sequenced by Illumina technique. Standard genome sequencing and standard bioinformatic analyses were provided by Oebiotech (Shanghai, China). The filtered reads were mapped to the reference genome *X. dendrorhous* rsharma.33.13 (EMBL: ERR575093-ERR575095) (28) using the BWA 0.7.16a software. Raw read sequencing data has been deposited in the NCBI database under the accession number PRJNA665546. The structural variations (SVs) of large genomic deletions, translocations, insertions, and inversions were predicted by BreakDancer (29). The aligned sequence reads were visualized by Integrative Genomics Viewer 2.8.0 (IGV) (30).

Deep sequencing was performed on the Illumina NovaSeq sequencing platform (Illumina Inc., San Diego, CA, USA) by Oebiotech (Shanghai, China). The PCR products (250∼280 bp) were purified using a MinElute PCR Purification Kit (QIAGEN, USA) and the libraries were constructed with VAHTS Universal DNA Library Prep Kit for Illumina V3 (Vazyme, Nanjing, China). Briefly, the purified products were end repaired and then adapters were ligated onto the 3’ end of the products. After PCR amplification and purification, the final libraries were sequenced and 150 bp paired-end reads were generated. The raw sequence reads were assembled by the software PEAR v0.9.6 (31). Clean reads were compared with the reference sequence by bowtie2 v2.3.4.1 (32). Sequencing the DNA regions containing the *CrtE* site 3 was repeated independent twice with three and five technical replicates, respectively. Each of the other regions was sequenced for five technical replicates. The sequencing depth reached 4∼6 × 10^6^ times.

### Chromosome karyotype detection

The chromosome karyotypes were detected by pulsed-field gel electrophoresis (PFGE) (33). Yeast protoplasts were made from the yeast cells in early exponential phase (OD_600_=1.0 to 1.2) that were exposed in protoplast-forming enzyme lywallzyme in 0.7 M KC1 supplemented with 0.2% β-mercaptoethanol and kept at 30°C under gentle shaking for 1.5 h. Sample plugs were prepared according to a standard procedure (33). Separations were carried out with a CHEF Mapper XA electrophoresis system (Bio-Rad). Gels were cast from chromosomal-grade agarose (0.8%, Bio-Rad) in 0.5 ×TBE (TRIS-borate, 45 mM; EDTA, 50 mM, pH 8.0). The running buffer was 0.5 × TBE, continuously circulated at 10°C. Electrophoretic conditions for CHEF: 125 V for 48 h with a switch time of 450 s, and 125 V for 24 h with a switch time of 250 s. Gels were stained for 30 min in 0.5 mg ml^-1^ ethidium bromide.

### Gene deletion

Gene deletion was performed by homologous recombination. Diagnostic PCR and Sanger sequencing were employed to identify positive gene-disruption clones. The sequences of homologous arms and primers used for diagnostic PCR were showed in Supplementary Data S2. For double-gene deletion, two rounds of homologous recombination were carried out and transformants were grown on YPD plates supplemented with 100 μg/ml of hygromycin or 200 μg/ml of Zeocin, respectively.

### Statistical analysis

To investigate on-target point mutations in clones without color change, the base frequency data acquired by deep sequencing on the Illumina X ten platform were employed. Base percentage of target DNA regions between edited clones and controls was analyzed. Standard deviation of the quotient of base frequency was calculated by the following equation:

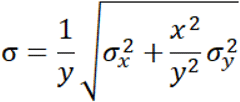

*σ*, standard deviation of the quotient of base frequency between edited clones and controls; *σ*_x_, the standard deviation of base frequency of edited clones. *σ*_y_, the standard deviation of base frequency of controls. x, average base frequency of edited clones. y, average base frequency of controls.

## RESULTS

### CRISPR-Cas9 genome editing induces diverse DNA repair patterns

We used the geranylgeranyl pyrophosphate synthase gene (*CrtE*) and the astaxanthin synthase gene (*CrtS*) as targets of CRISPR-Cas9 because inactivation of *CrtE* and *CrtS* results in color changes of red *X. dendrorhous* clones to white and yellow, respectively. The *Cas9* and gRNA genes were inserted at the ribosomal RNA gene loci and confirmed to show constitutive expression (Supplementary Figure S1A-C). Nineteen target sites in *CrtE* and *CrtS* of the wild type strain CBS 6938 were edited to generate mutants with changed pigment (Supplementary Figure S1D,E, and Data S2).

We identified 248 white mutants and 35 yellow mutants from 623,727 colonies after targeting strain CBS 6938. DNA deletion at target sites was detected by PCR amplification and Sanger sequencing. Most deletions were smaller than 1 kb in size and the largest deletion was 6.1 kb (Figure 1A), similar to previous reports (3-6, 15). In addition to DNA deletion, unexpected DNA repairs were also observed. Notably, we detected nucleotide substitutions near the PAM sequences of site 3 and site 7 in the *CrtE* gene (Figure 1B,C), representing a new kind of repair outcome of CRISPR-Cas9 genome editing. At *CrtE* site 3, more than half (54/84) of the sequenced mutants contain a T•A to A•T point mutations at position –2 and/or a C•G to A•T mutation at position –5 (Figure 1B). At *CrtE* site 7, a G•C to A•T point mutation at position –1 was observed (Figure 1C). These unexpected DNA repairs may profoundly influence the evaluation of precise editing by CRISPR-Cas9. PCR amplifications failed to obtain DNA amplicons from all the mutants edited at *CrtE* site 2, most of the mutants edited at *CrtE* site 7, and all *CrtS* sites (Supplementary Figure S2, S3), indicating that complex chromosome rearrangements occurred in the edited clones. Additionally, short reverse complement and deletion accompanying the insertion of short repeats were found at *CrtE* site 3 (Figure 1D-F).

**Figure 1.**
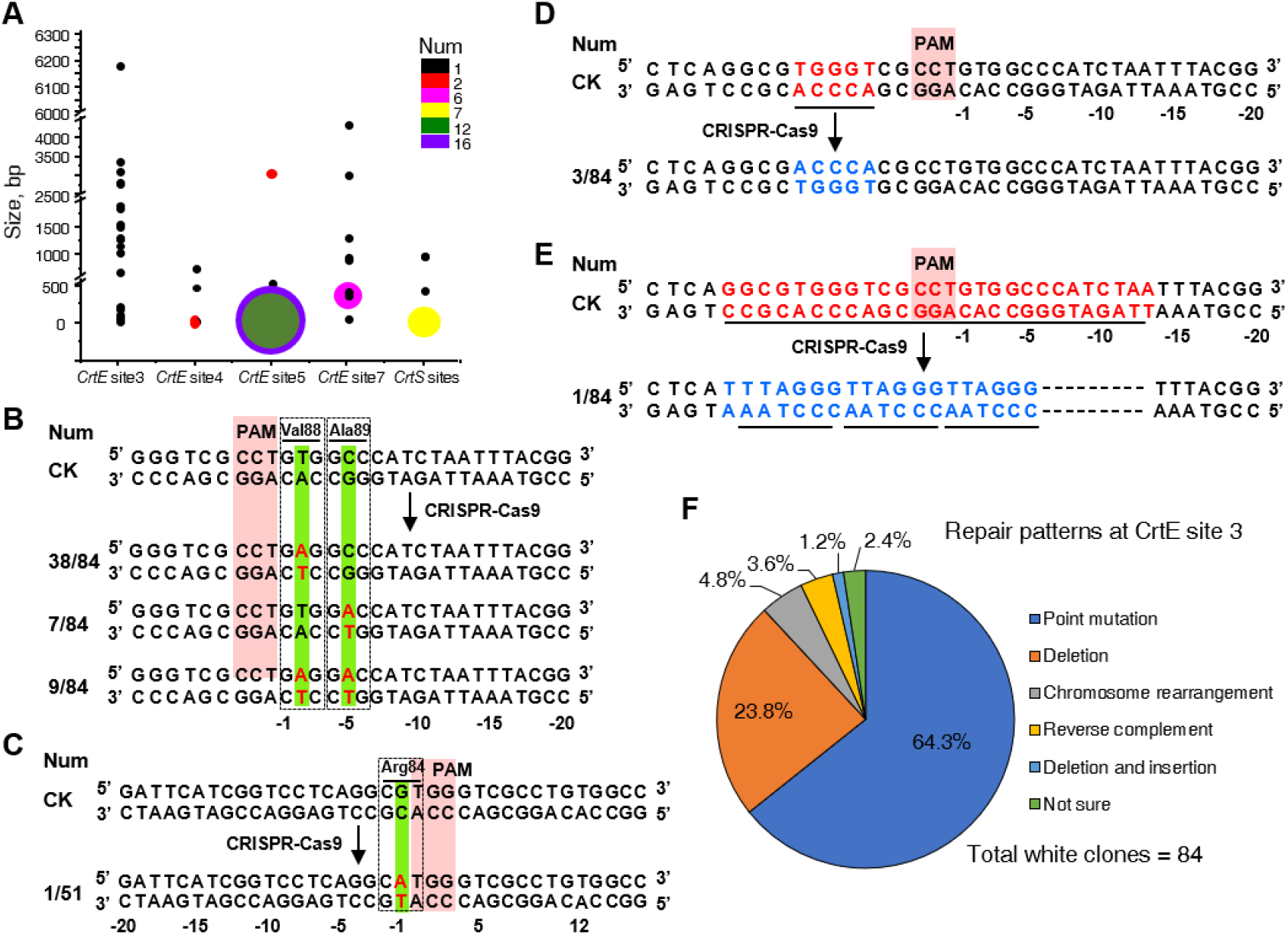
DNA repair patterns identified in CRISPR-Cas9 editing *CrtE* and *CrtS* in *X. dendrorhous*. (**A**) The size and number of the deleted DNA regions observed in white edited clones. (**B** and **C**) Nucleotide substitutions adjacent to the PAM sequences of the target *CrtE* site 3 (B) and site 7 (C) identified in white edited clones. The total numbers of white edited clones and those possessing nucleotide substitutions are shown. The light green highlights indicate the point-mutations after CRISPR-Cas9 gene editing. (**D** and **E**) DNA repair patterns: short reverse complement (D) and deletion accompanying insertion of short repeats (E) identified in edited *CrtE* site 3. (**F**) Diverse DNA repair patterns were observed in white edited clones in editing *CrtE* site 3. No donor DNA was supplied for these editing experiments. PAM, protospacer-associated motif. Num, numbers. CK, control.

### On-target point mutations resulting from CRISPR-Cas9 genome editing

We further assessed whether the identified nucleotide substitutions resulted from random mutation or DNA repair induced by on-target cleavage of Cas9. The point mutations T•A to A•T at position –2 and C•G to A•T at position –5 of *CrtE* site 3 encode amino acid variations V88E and A89D, respectively, and both changes resulted in white clones (Supplementary Figure S4A). Mutation analysis of *CrtE* site 7 showed that the nucleotide substitution G•C to A•T corresponding to the amino acid variation R84H resulted in white clones (Supplementary Figure S4B and Figure 1C). Saturated mutation analysis at Val88 and Ala89 sites demonstrated that variations V88L, V88M, and A89V also produced white clones, but these mutations were not observed in sequenced CRISPR-Cas9 edited clones (Supplementary Figure S4A, and Figure 1B,C). Structural modeling of CrtE showed that Val88 and Ala89 are located near the active sites (Supplementary Figure S4C,D). These results indicated that the observed on-target point mutations resulted from endogenous repair induced by CRISPR-Cas9 cleavage rather than random mutation.

Nucleotide substitutions that do not produce different color colonies cannot be identified by plate screening. To further investigate the extent of mutation at the CRISPR-Cas9 editing sites, we performed deep sequencing on Illumina platforms. DNA regions containing CRISPR-Cas9 target sites with a size of 250 bp to 280 bp were amplified from ∼1×10^4^ colonies for each edited site and sequenced. No on-target mutations were observed at *CrtS* sites 1 and 8 (Supplementary Figure S5), consistent with the results of DNA amplification and Sanger sequencing. At *CrtE* site 3, in addition to the mutations identified by sequencing white mutants, a low-frequency mutation T•A to C•G at position –2 was detected (Figure 2A). The sequencing results showed the occurrence of rare on-target mutations, with G•C to C•G at position –1 and C•G to T•A at position –7 of *CrtE* site 3, and G•C to C•G inside the PAM sequence of *CrtE* site 7 (Figure 2). The results indicate that both high-frequency and rare on-target mutations can occur during DNA repair induced by CRISPR-Cas9 cleavage.

**Figure 2.**
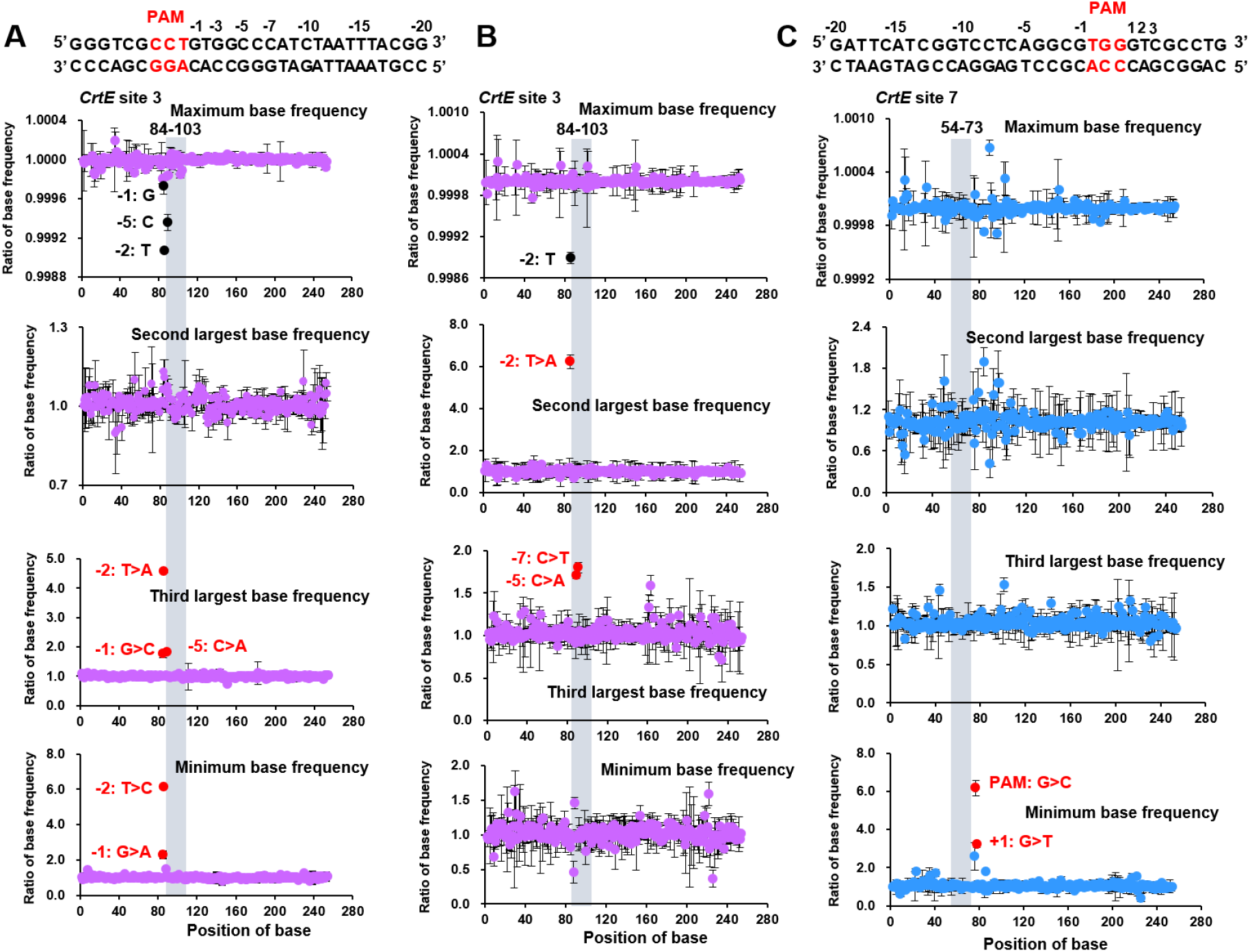
Identification of on-target nucleotide substitutions from DNA repair induced by CRISPR-Cas9 cleavage using deep sequencing. Ratio of base frequency of edited clones to the control for the four possibilities. (**A**) Ratio of base frequency at *CrtE* site 3 calculated based on the sequencing data for the DNA region containing the site; three technical repeats. (**B**) Results of a separate biological replicate experiment; data from sequencing the DNA region containing *CrtE* site 3; five technical repeats. (**C**) Ratio of base frequency at *CrtE* site 7 calculated using the sequencing data; five technical repeats. The light gray regions denote the 20-bp target sequence. PAM, protospacer-associated motif.

### Chromosomal rearrangements induced by CRISPR-Cas9

Chromosomal rearrangements in edited clones were evaluated by chromosomal karyotype analysis using pulsed-field gel electrophoresis (PFGE) and Illumina sequencing. Electrophoretic karyotype analysis revealed the loss of a large chromosome and the appearance of a mid-size chromosome in some of the CRISPR-Cas9 edited clones compared with the controls, indicating chromosomal rearrangements that also explained the failure of PCR amplification to produce amplicons (Supplementary Figure S2, S6). The altered chromosomes in the edited mutants at *CrtE* sites 2, 3, 5, and 7 showed equal sizes in the PFGE gels, suggesting a similar rearrangement mechanism had occurred (Supplementary Figure S6).

Structural variation analysis using Illumina sequencing data revealed interchromosomal translocations in the edited clones in which DNA regions spanning *CrtE* site 2 or *CrtS* sites were not successfully amplified by PCR (Figure 3A). The translocations happened between the promoter of the alcohol dehydrogenase gene *ADH4* (LN483144.1) and the Cas9 cutting positions in all tested strains (Supplementary Table S1). In different edited clones, there were identical break positions from Cas9 cutting at the same target sites (Supplementary Table S1). In detail, the breaks occurred at position – 1 of *CrtE* site 2 (Figure 3B) and adjacent to the PAM regions of *CrtS* sites (Figure 3C and Supplementary Figure S7A,B). In the clones edited at the *CrtE* target site 3, the break positions were not confirmed due to DNA deletions (Figure 3D). Cas9 expression was controlled by the endogenous *ADH4* promoter, and this may result in DNA breaking due to homologous recombination. Thus, we suggest that the observed interchromosomal translocations were generated through two breaks, one from homologous recombination at the *ADH4* locus and the other from on-target cutting by Cas9.

**Figure 3.**
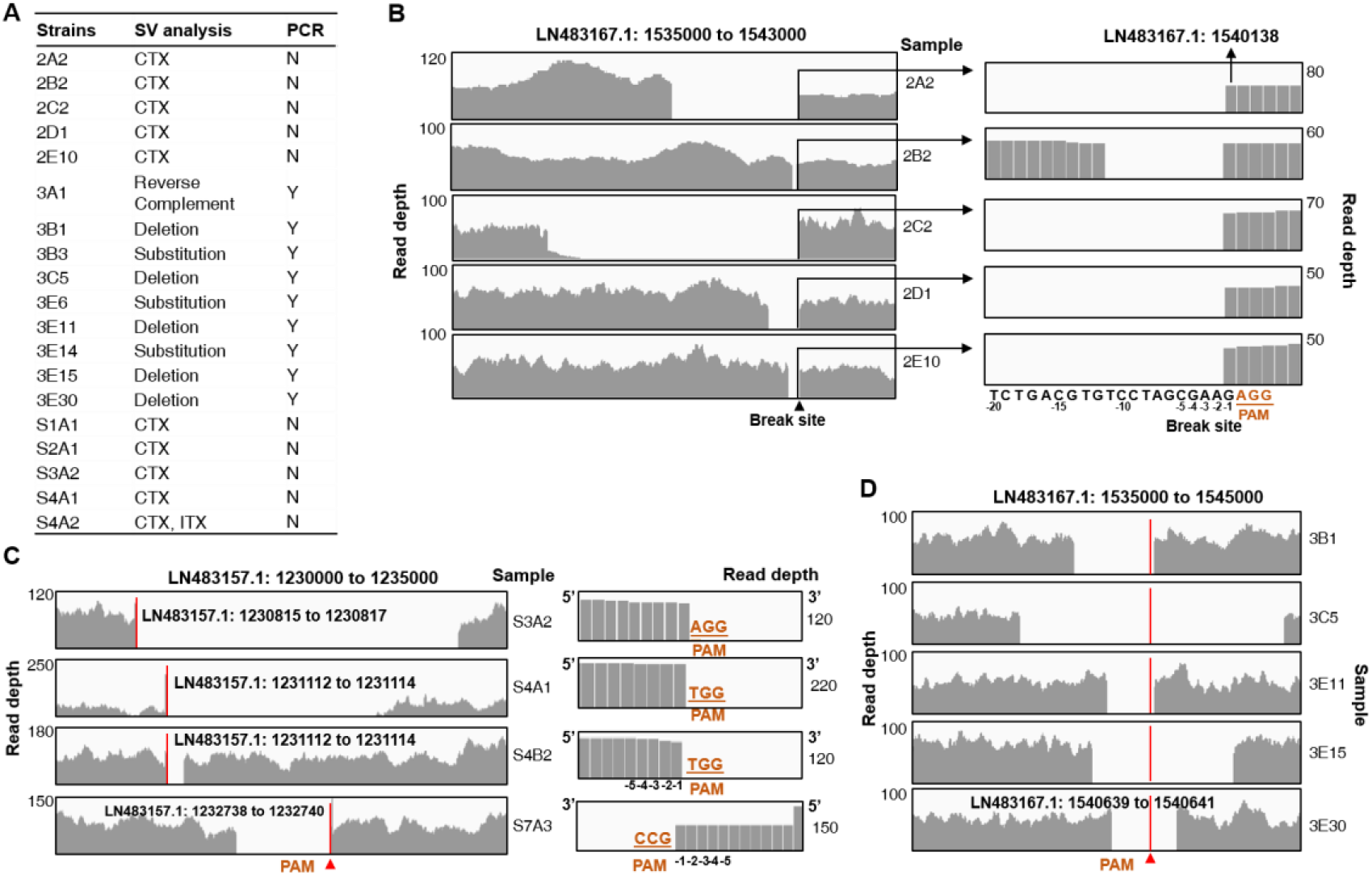
Analysis of chromosome rearrangements induced by CRISPR-Cas9 using genome sequencing. (**A**) Strains whose genomes were sequenced on Illumina platform. Structural variations (SVs) of large genomic deletions, translocations, insertions, and inversions are predicted by BreakDancer software package. CTX, interchromosomal translocation; ITX, intrachromosomal translocation; Y, positive PCR amplification; N, negative PCR amplification. (**B** and **C**) Break sites near PAM sequences induced by editing *CrtE* site 2 (B) and *CrtS* sites (C). PAM, protospacer-associated motif. (**D**) Diagram showing DNA deletion near *CrtE* site 3 in DNA repair induced by CRISPR-Cas9 editing.

### Pathways controlling the DNA repair in CRISPR-Cas9 genome editing

Point mutations and deletions were identified as the main repair patterns from editing *CrtE* site 3 (Figure 1). Repair of CRISPR-Cas9 cleavage requires endogenous cellular pathways (Supplementary Figure S8A). Analysis of deletion sequences showed that high frequency of microhomologies represented at the ends of the deletions (Supplementary Figure S8 and Data S4), indicating the involvement of the MMEJ pathway (34,35). However, it is unclear how the on-target point mutations were generated. To address this issue, we performed gene editing at the *CrtE* target site 3 in a series of strains disrupted for specific genes in repair pathways and acquired 193 white mutants from 400,187 colonies.

Disruption of the mismatch repair (MMR) proteins encoded by genes *MSH2* and *MLH1* (36) did not significantly alter the frequency of edited white clones (all observed mutants) or point mutations at *CrtE* site 3 (Figure 4A,B). We next investigated the NHEJ repair pathway. Deletion of the genes encoding Ku70 or Ku80 led to a sharp drop-off in the frequency of white edited clones and point mutation repair clones (Figure 4A,B). The Mre11-Rad50-Xrs2 (MRX) complex is one of the primary complexes responsible for DSBs repair in eukaryotes (4,37). Deletion of *Mre11* or *RAD50* resulted in a loss of white edited clones, indicating the dependence of DNA repair induced by CRISPR-Cas9 cleavage on MRX (Figure 4A,B). The Sae2 protein in budding yeast and the human ortholog CtIP stimulate Mre11 endonuclease activity (34). However, deletion of *Sae2* in *X. dendrorhous* did not reduce the frequency of white edited clones (Figure 4A,B), indicating that the endonuclease activity of Mre11 did not contribute to the DNA repair induced by Cas9 cleavage.

**Figure 4.**
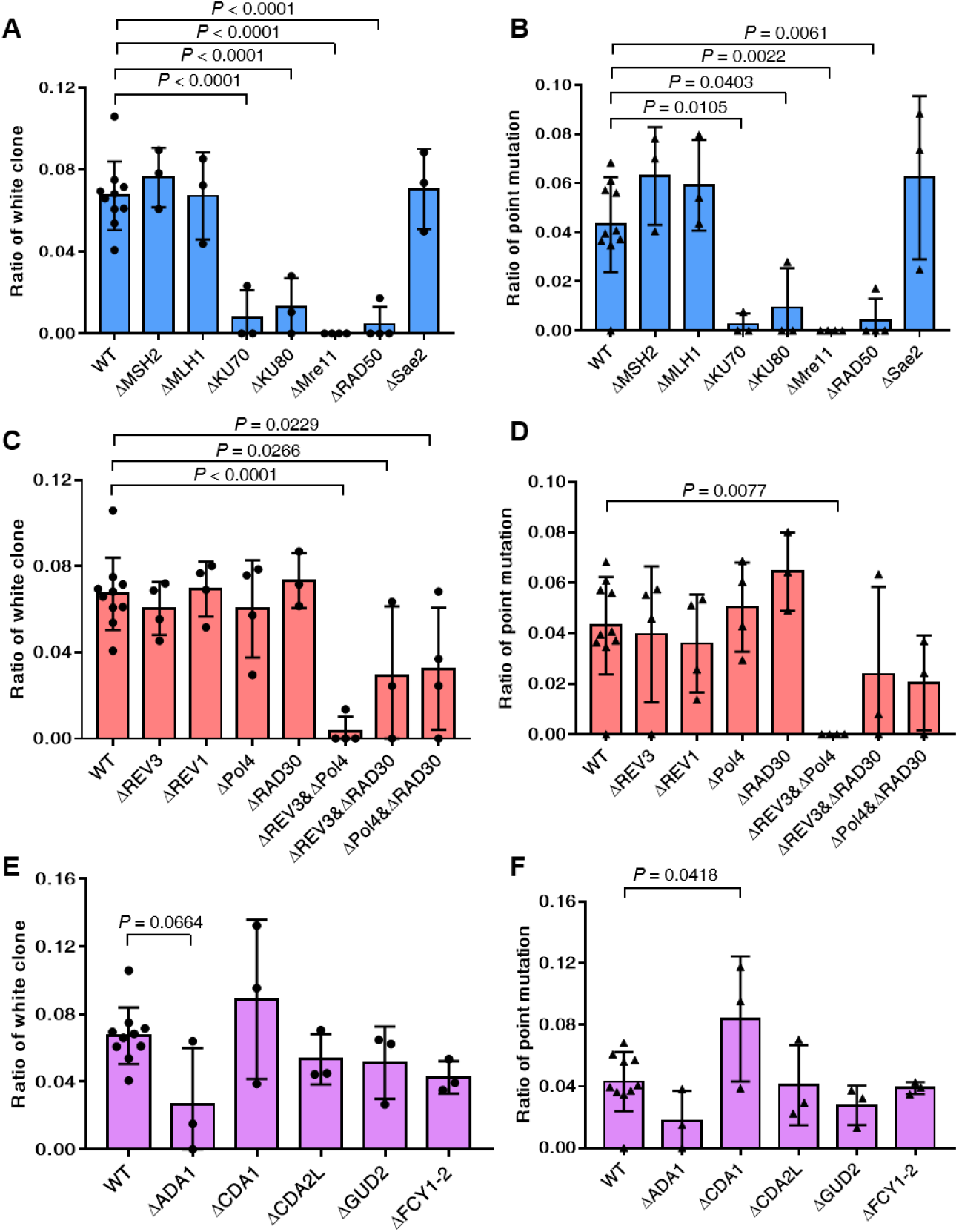
Identification of the cellular repair pathways participating in DNA repair induced by CRISPR-Cas9 cleavage. (**A** and **B**) Elucidation of the influence of genes in MMR and NHEJ pathways on DNA repair induced by Cas9 cleavage. (**C** and **D**) Elucidation of the influence of non-essential DNA polymerases on DNA repair induced by Cas9 cleavage. (**E** and **F**) Evaluation of the effect of deaminases on DNA repair induced by Cas9 cutting. The ratio of white edited clones (A, C, E) and nucleotide substitution repair clones (B, D, F) to all transformants growing on antibiotic-resistant plates, respectively. A-F, Data are the mean ± SD of at least three biological replicates. *P* values were determined by one-way analysis of variance (ANOVA) with Dunnett’s multiple comparisons test.

We also assessed the influence of non-essential DNA polymerases on DNA repair induced by CRISPR-Cas9 cleavage, given their low-fidelity properties. Disruptions of polymerases Pol4, Rev1, Rev3, and RAD30, orthologs of *Saccharomyces cerevisiae* (38), by single-gene deletion in *X. dendrorhous* did not significantly alter gene editing at the *CrtE* target site 3 (Figure 4C,D). Considering the redundancy of non-essential DNA polymerases, double deletion analysis was next performed. CRISPR-Cas9 editing tests showed that double deletion of *REV3*/*RAD30* (*p*=0.0266) and *POL4*/*RAD30* (*p*=0.0229) both led to significant reduction in the survival of white clones (Figure 4C,D). Noticeably, double deletion of *REV3* and *POL4* genes resulted in a 10-fold reduction in the survival of white clones and loss of the point mutation repair clones (Figure 4C,D). These results demonstrated overlapping roles effects of non-essential DNA polymerases, particularly *REV3* and *POL4*, to affect DNA repair induced by CRISPR-Cas9 cleavage.

We further analyzed the influence of deaminases on the DNA repair induced by CRISPR-Cas9 cleavage because deaminases can induce both transition and transversion events (39). Single deletion of the genes encoding the deaminases ADA1, CDA1, CDA2L, GUD2, and FCY1-2 did not evidently reduce the survival of white clones and point mutations (Figure 4E,F). In the budding yeast *S. cerevisiae*, Cas9 protein can also exhibit nickase activities, although this activity is not predominant (40). We tested previously characterized nicking Cas9 variants, D10A and H840A (41), but no point mutations were observed, indicating that the point mutations were not due to DNA nicking. Based on these findings and previous studies on DNA repair (42,43), we proposed that on-target point mutations were generated as a result of the low-fidelity of the non-essential DNA polymerases that participate in NHEJ repair induced by CRISPR-Cas9 gene editing (Figure 5)

**Figure 5.**
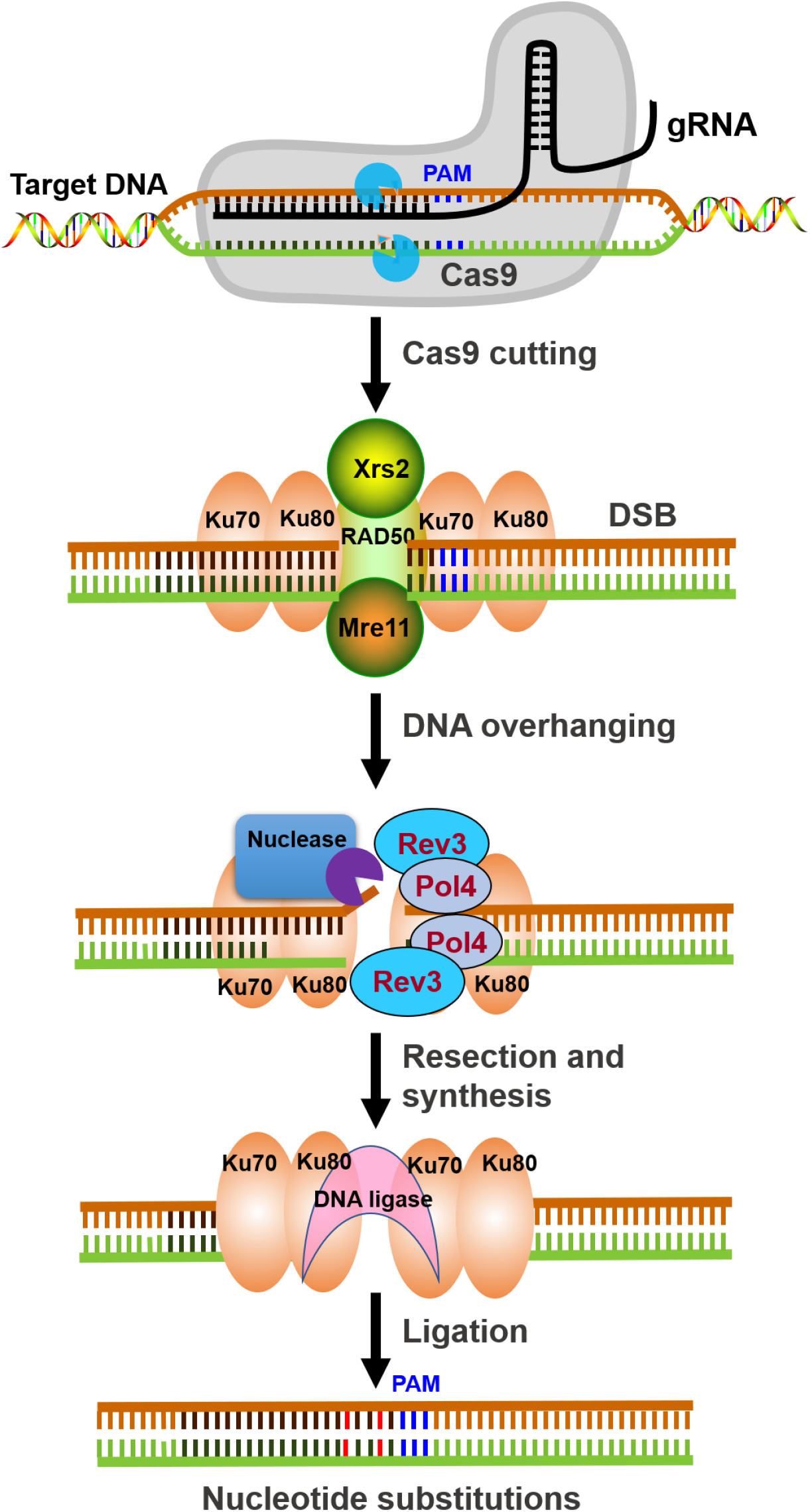
A model of nucleotide substitution repair induced by CRISPR-Cas9 editing. Low-fidelity and non-essential DNA polymerases, such as Rev3 and Pol4, may contribute to the observed DNA repair pattern of nucleotide substitutions. PAM, protospacer-associated motif. DSB, double strand break.

## DISCUSSION

Repair of CRISPR-Cas9-mediated DSBs is one of the key events in the application of CRISPR-Cas9 technique. Such DNA repair can be finished by hosts themselves or by hosts with exogenous donor DNA. When no donor DNA is supplied, hosts repairing DSBs cause DNA deletion/insertion (3-6) or low frequency of genomic rearrangements (15) and nucleotide substitutions (4, 16-18). Nucleotide substitutions or point mutations are not easily detected in genome editing for several reasons. First, the frequency of point mutations induced by Cas9 cutting is far lower than deletion/insertion. Second, most point mutations do not typically lead to obvious phenotypes. Third, present detection methods cannot fully identify all DNA repair outcomes after Cas9 cleavage (44). In this study, we identified high-frequency on-target mutations in specific gene sites induced by repairing of CRISPR-Cas9 cleavage in *X. dendrorhous*. Different from a previous study in *Saccharomyces cerevisiae* (4), our findings suggested that the on-target mutations in *X. dendrorhous* cannot render the gRNA effective and they are not random mutations. The point mutations T•A to A•T at position –2 and C•G to A•T at position –5 of *CrtE* site 3 are reproducible (Figure 1B). Moreover, these point mutations showed strong sequence dependence that is not consistent with the observations in Hela cells, in which CRISPR-mediated substitutions were considered lacking sequence dependence and conversion preferences (18). The differences may be attributed to different organisms and/or methods used in these studies. Additionally, similar to previous studies (4, 18), this study also found that substitutions frequently occur near CRISPR-Cas9 cleavage sites and the substitution frequency in most gRNA sites was extremely low while in some specific sites was even high than 40%.

The factors leading to point mutations in repairing Cas9 cleavage in *X. dendrorhous* were systematically analyzed in this study. Our data supported that all repair events were not induced by nicking Cas9. Although deaminases fused with dead or nicking Cas9 have been developed as base editing techniques to introduce point mutations (39,45,46), the point mutations observed in this study were not resulted from the activity of host’s deaminases. Dead Cas9 targeting can induce mutations in *S. cerevisiae* (47,48). In this study, the repair events in individuals of populations might mediated by either DSBs or non-cutting Cas9. By gene-deletion analysis, we demonstrated that on-target point mutations were generated as a result of the low-fidelity of the non-essential DNA polymerases that participate in NHEJ repair induced by CRISPR-Cas9 genome editing. A previous study pinpointed the correlation between DNA ligase IV and nucleotide substitutions (18). Our study further demonstrated that each of the genes *Ku70, Ku80, Mre11*, or *RAD50*, and the overlapping roles of *REV3* and *POL4* were necessary for the emergence of point mutations. When these genes were disrupted, other repair patterns also dramatically decreased along with point mutations. Therefore, the specific mechanism of point-mutation formation in self-repairing the Cas9-mediated DSBs remains to be elucidated. The enzymes *Mre11* and *POL4* were involved not only in the formation of point mutations as shown in this study, but also in the generation of insertion/deletions induced by CRISPR-Cas9 treatment in *S. cerevisiae* (4). Nevertheless, the dependence of point mutations on *Ku70, Ku80*, and DNA ligase 4 is not in accordance with the viewpoint that NHEJ of Cas9 cleavages are not more sensitive to the absence of Ku or DNA ligase 4 in *S. cerevisiae* (4). Different repair patterns may lead to the difference. Our study focused on point mutants, while the previous publication highlighted the repair pattern of insertion/deletions (4).

For safe clinical use, genome editing needs to be accurate. The efficacy of CRISPR–Cas9-induced genome editing has been demonstrated in various cells, but control of an exact editing outcome remains a challenge (49, 50). Recognizing the potential DNA repair outcomes and understanding the DNA repair mechanisms after on-target DNA cleavage facilitate precise gene editing. Our study suggested that the overlapping roles of the non-essential DNA polymerases and the components (*Ku70, Ku80, Mre11*, or *RAD50*) of NHEJ are among the key factors leading to on-target point mutations in CRISPR-Cas9 genome editing. Given that many point mutations occurring in human cancer cells arise from the error-generating activities of low-fidelity DNA polymerases (51,52) and NHEJ is an evolutionarily conserved pathway to repair DSBs (53), more attention should be paid to on-target point mutations induced by CRISPR-Cas9.

## DATA AVAILABILITY

Sequence data generated in this study have been deposited in the NCBI Sequence Read Archive (SRA) database under the accession numbers PRJNA665073 and PRJNA665546.

## SUPPLEMENTARY DATA

Supplementary Data are available at NAR online.

## ACKNOWLEDGEMENT

We thank Y. L. Shen (State Key Laboratory of Microbial Technology, Shandong University, China) and C. H. Bi (Tianjin Institute of Industrial Biotechnology, Chinese Academy of Sciences) for valuable discussions.

## REFERENCES

1. Mali, P., Yang, L.H., Esvelt, K.M., et al. (2013). RNA-guided human genome engineering via Cas9. Science 339, 823–826.

2. Cubbon, A., Ivancic-Bace, I., Bolt, E.L. (2018). CRISPR-Cas immunity, DNA repair and genome stability. Biosci. Rep. 38, BSR20180457.

3. van Overbeek, M., Capurso, D., Carter, M. M., et al. (2016). DNA repair profiling reveals nonrandom outcomes at Cas9-mediated breaks. Mol. Cell 63, 633–646.

4. Lemos, B.R., Kaplan, A. C., Bae, J. E., et al. (2018). CRISPR/Cas9 cleavages in budding yeast reveal templated insertions and strand-specific insertion/deletion profiles. Proc. Natl. Acad. Sci. U.S.A. 115, E2040–E2047.

5. Pannunzio, N.R., Watanabe, G., Lieber, M.R. (2018). Nonhomologous DNA end-joining for repair of DNA double-strand breaks. J. Biol. Chem. 293, 10512–10523.

6. Allen, F., Crepaldi, L., Alsinet, C., et al. (2019). Predicting the mutations generated by repair of Cas9-induced double-strand breaks. Nat. Biotechnol. 37, 64–72.

7. Filippo, J.S., Sung, P., Klein, H. (2008). Mechanism of eukaryotic homologous recombination. Annu. Rev. Biochem. 77, 229–257.

8. Chu, V.T., Weber, T., Wefers, B., et al. (2015). Increasing the efficiency of homology-directed repair for CRISPR-Cas9-induced precise gene editing in mammalian cells. Nat. Biotechnol. 33, 543–548.

9. Maruyama, T., Dougan, S.K., Truttmann, M.C., et al. (2015). Increasing the efficiency of precise genome editing with CRISPR-Cas9 by inhibition of nonhomologous end joining. Nat. Biotechnol. 33, 538–542.

10. Canny, M.D., Moatti, N., Wan, L.C.K., et al. (2018). Inhibition of 53BP1 favors homology-dependent DNA repair and increases CRISPR-Cas9 genome-editing efficiency. Nat. Biotechnol. 36, 95–102.

11. Kato-Inui, T., Takahashi, G., Hsu, S., et al. (2018). Clustered regularly interspaced short palindromic repeats (CRISPR)/CRISPR-associated protein 9 with improved proof-reading enhances homology-directed repair. Nucleic Acids Res. 46, 4677–4688.

12. Richardson, C.D., Ray, G.J., DeWitt, M.A., et al. (2016). Enhancing homology-directed genome editing by catalytically active and inactive CRISPR-Cas9 using asymmetric donor DNA. Nat. Biotechnol. 34, 339–344.

13. Shen, M.W., Arbab, M., Hsu, J.Y., et al. (2018). Predictable and precise template-free CRISPR editing of pathogenic variants. Nature 563, 646–651.

14. Molla, K.A., and Yang, Y. (2020). Predicting CRISPR/Cas9-induced mutations for precise genome editing. Trends Biotechnol. 38, 136–141.

15. Kosicki, M., Tomberg, K., Bradley, A. (2018). Repair of double-strand breaks induced by CRISPR-Cas9 leads to large deletions and complex rearrangements. Nat. Biotechnol. 36, 765–771.

16. Bassett, A.R., Tibbit, C., Ponting, C.P., Liu, J.L. (2013). Highly efficient targeted mutagenesis of Drosophila with the CRISPR/Cas9 system. Cell Rep. 4, 220–228.

17. Wang, T., Wei, J.J., Sabatini, D.M., et al. (2014). Genetic screens in human cells using the CRISPR-Cas9 system. Science. 343, 80–84.

18. Hwang G.H., Yu J., Yang S., Son W.J., Lim K., Kim H.S., Kim J.S., Bae S. (2020). CRISPR-sub: analysis of DNA substitution mutations caused by CRISPR–Cas9 in human cells. Comput. Struct. Biotechnol. J. 18,1686–1694.

19. Mannazzu, I., Landolfo, S., Lopes da Silva, T., et al. (2015). Red yeasts and carotenoid production: outlining a future for non-conventional yeasts of biotechnological interest. World J. Microbiol. Biotechnol. 31, 1665–1673.

20. Wery, J., Verdoes, J.C., van Ooyen, A.J.J. (1998). Efficient transformation of the astaxanthin-producing yeast Phaffia rhodozyma. Biotechnol. Tech. 12, 399–405.

21. Gao, S., Tong, Y., Wen, Z., et al. (2016). Multiplex gene editing of the Yarrowia lipolytica genome using the CRISPR-Cas9 system. J. Ind. Microbiol. Biot. 43, 1085–1093.

22. Lorenz, R., Bernhart, S.H., Honer Zu Siederdissen, C., et al. (2011). ViennaRNA Package 2.0. Algorithm. Mol. Biol. 6, 26.

23. Blin, K., Pedersen, L.E., Weber, T., et al. (2016). CRISPy-web: An online resource to design sgRNAs for CRISPR applications. Synth. Syst. Biotechnol. 1, 118–121.

24. Waterhouse, A., Bertoni, M., Bienert, S., et al. (2018). SWISS-MODEL: homology modelling of protein structures and complexes. Nucleic. Acids Res. 46, W296–W303.

25. Kavanagh, K.L., Dunford, J.E., Bunkoczi, G., et al. (2006). The crystal structure of human geranylgeranyl pyrophosphate synthase reveals a novel hexameric arrangement and inhibitory product binding. J. Biol. Chem. 281, 22004–22012.

26. Martinez-Moya, P., Watt, S.A., Niehaus, K., et al. (2011). Proteomic analysis of the carotenogenic yeast Xanthophyllomyces dendrorhous. BMC Microbiol. 11, 131.

27. DiCarlo, J.E., Norville, J.E., Mali, P., et al. (2013). Genome engineering in Saccharomyces cerevisiae using CRISPR-Cas systems. Nucleic Acids Res. 41, 4336–4343.

28. Sharma, R., Gassel, S., Steiger, S., et al. (2015). The genome of the basal agaricomycete Xanthophyllomyces dendrorhous provides insights into the organization of its acetyl-CoA derived pathways and the evolution of Agaricomycotina. BMC Genomics 16, 233.

29. Chen, K., Wallis, J.W., McLellan, M.D., et al. (2009). BreakDancer: an algorithm for high-resolution mapping of genomic structural variation. Nat. Methods 6, 677–681.

30. Robinson, J.T., Thorvaldsdottir, H., Winckler, W., et al. (2011). Integrative genomics viewer. Nat. Biotechnol. 29, 24–26.

31. Zhang, J.J., Kobert, K., Flouri, T., et al. (2014). A. Stamatakis, PEAR: a fast and accurate Illumina Paired-End reAd mergeR. Bioinformatics 30, 614–620.

32. Langmead, B., and Salzberg, S.L. (2012). Fast gapped-read alignment with Bowtie 2. Nat. Methods 9, 357–359.

33. Nagy, A., Garamszegi, N., Vagvolgyi, C., et al. (1994). Electrophoretic karyotypes of Phaffia rhodozyma strains. FEMS Microbiol. Lett. 123, 315–318.

34. Meyer, D., Fu, B.X.H., Heyerar W.D. (2015). DNA polymerases delta and lambda cooperate in repairing double-strand breaks by microhomology-mediated end-joining in Saccharomyces cerevisiae. P. Natl. Acad. Sci. U.S.A. 112, E6907–E6916.

35. Owens, D.D.G., Caulder, A., Frontera, V., et al. (2019). Microhomologies are prevalent at Cas9-induced larger deletions. Nucleic Acids Res. 47, 7402–7417.

36. Gupta, D., and Heinen, C.D. (2019). The mismatch repair-dependent DNA damage response: Mechanisms and implications. DNA Repair 78, 60–69.

37. Paull, T.T. (2018). 20 years of Mre11 biology: no end in sight. Mol. Cell 71, 419–427.

38. Meyer, D., Fu, B.X.H., Chavez, M., et al. (2019). Cooperation between non-essential DNA polymerases contributes to genome stability in Saccharomyces cerevisiae. DNA Repair 76, 40–49.

39. Kim, H.S., Jeong, Y.K., Hur, J.K., et al. (2019). Adenine base editors catalyze cytosine conversions in human cells. Nat. Biotechnol. 37, 1145–1148.

40. Fu, B.X.H. Smith, J.D., Fuchs, R.T., et al. (2019). Target-dependent nickase activities of the CRISPR-Cas nucleases Cpf1 and Cas9. Nat. Microbiol. 4, 888–897.

41. Jinek, M., Chylinski, K., Fonfara, I., et al. (2012). A programmable dual-DNA-guided DNA endonuclease in adaptive bacterial immunity. Science 337, 816–821.

42. Li, P., Li, J., Li, M., et al. (2012). Multiple end joining mechanisms repair a chromosomal DNA break in fission yeast. DNA Repair 11, 120–130.

43. Chang, H.H.Y., Pannunzio, N.R., Adachi, N., et al. (2017). Non-homologous DNA end joining and alternative pathways to double-strand break repair. Nat. Rev. Mol. Cell Bio. 18, 495–506.

44. Roidos, P., Sungalee, S., Benfatto, S., et al. (2020). A scalable CRISPR/Cas9-based fluorescent reporter assay to study DNA double-strand break repair choice. Nat. Commun. 11, 4077.

45. Gaudelli, N.M., Komor, A.C., Rees, H.A., et al. (2017). Programmable base editing of A.T to G.C in genomic DNA without DNA cleavage. Nature 551, 464–471.

46. Komor, A.C., Kim, Y.B., Packer, M.S., et al. (2016). Programmable editing of a target base in genomic DNA without double-stranded DNA cleavage. Nature 533, 420–424.

47. Doudna, J.A. (2020). The promise and challenge of therapeutic genome editing. Nature 578, 229–236.

48. Laughery, M.F., Mayes, H.C., Pedroza, I.K. and Wyrick, J.J. (2018). R-loop formation by dCas9 is mutagenic in Saccharomyces cerevisiae. Nucleic Acids Res. 47, 2389–2401.

49. Doi, G., Okada, S., Yasukawa, T., et al. (2021). Catalytically inactive Cas9 impairs DNA replication fork progression to induce focal genomic instability. Nucleic Acids Res. 49, 954–968.

50. Shivram, H., Cress, B.F., Knott, G.J., et al. (2021). Controlling and enhancing CRISPR systems. Nat. Chem. Biol. 17, 10–19.

51. Lange, S.S., Takata, K., Wood, R.D. (2011). DNA polymerases and cancer. Nat. Rev. Cancer 11, 96–110.

52. Reijns, M.A.M., Kemp, H., Ding, J., et al. (2015). Lagging-strand replication shapes the mutational landscape of the genome. Nature 518, 502–506.

53. Emerson, C.H., and Bertuch, A.A. (2016). Consider the workhorse: Nonhomologous end-joining in budding yeast. Biochem. Cell Biol. 94, 396–406.

