## Supplemental Figures and Tables for "The cause of on-target point mutations generated by CRISPR-Cas9 treatment in the yeast *Xanthophyllomyces dendrorhous*"

**This PDF file includes:**

Figures S1 to S8

Tables S1 and S2

Captions for Data S1 to S4

**Other Supplementary Materials for this manuscript include the following:**

Data S1 [Primers and plasmids used in this study]

Data S2 [Characteristics of the DNA sequences used in this study]

Data S3 [CRISPR-Cas9 edited and pigment-changed clones]

Data S4 [Deletion sequences generated in repair of Cas9 cleavages]

Source Data [Figure 1, Figure 2A, Figure 2B, Figure 2C, Figure 4; Figure S2, Figure S3, Figure S5A, Figure S5B]

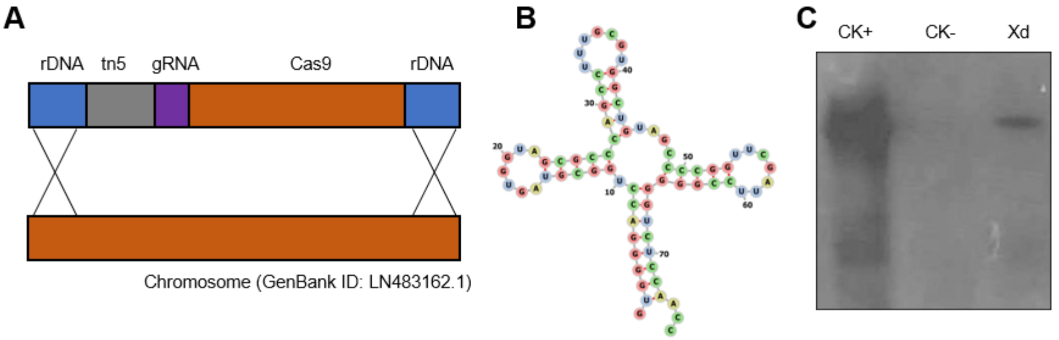


D

| Target | Sequence (5'-3') | PAM |
|---|---|---|
| E1 | CTCAATATACTTCCAGATCA | GGG |
| E2 | TCTGACGTGTCCTAGCGAAG | AGG |
| E3 | CCGTAAATTAGATGGGCCAC | AGG |
| E4 | TCCCATTAGAATGGACGATG | TGG |
| E5 | CATTAGAATGGACGATGTGG | AGG |
| E6 | GGACGATGTGGAGGATTCAT | CGG |
| E7 | GATTCATCGGTCCTCAGGCG | TGG |
| E8 | TCATATAGTCATCCCTGATC | TGG |
| E9 | TATGAACCTTCAGTCTTCTG | AGG |

E

| Target | Sequence (5'-3') | PAM |
|---|---|---|
| S1 | CGAGTGCTCAGAATTGCTAC | CGG |
| S2 | ATCTCAAAAGCTGTCCGATG | AGG |
| S3 | AAGCTCATAAGCGACATCGA | AGG |
| S4 | TCGAGCTACTCTGGACGTCA | TGG |
| S5 | TTCTCGAAGTTTATCCTGAA | CGG |
| S6 | GAACCAACCGCTTCGAGATA | AGG |
| S7 | ACGTCCTCAAGCCACCTCTC | CGG |
| S8 | AACAGTATTGAAGCACCCTA | TGG |
| S9 | AAACTTACAAGCAAGCTCT | GGG |
| S10 | GAAACTTACAAGCAAGCTC | TGG |

**Figure S1. CRISPR-Cas9 system for the yeast *X. dendrorhous* and the target sites.** (**A**) Diagram of CRISPR-Cas9 DNA cassettes recombining into the chromosomal rRNA gene loci in engineered *X. dendrorhous* strains. rDNA, ribosomal rDNA; tn5, gene encoding geneticin resistance; gRNA, guide RNA. Cas9 and gRNA are constitutively expressed. (**B**) Structural analysis of a predicted tRNA-Ala sequence retrieved by tRNAscan-SE 2.0 software using ViennaRNA Package 2.0 algorithms. The tRNA-Ala is used to control the expression of gRNA. (**C**) Assay of the Cas9 expression by western blotting. CK+, commercial Cas9 proteins; CK-, protein extracts from strains without Cas9 genes; Xd, protein extracts from *X. dendrorhous* strains expressing a codon-optimized *Cas9* gene. (**D** and **E**) Selected target sites in the *CrtE* gene (D) and the *CrtS* gene (E). PAM, protospacer-associated motif.

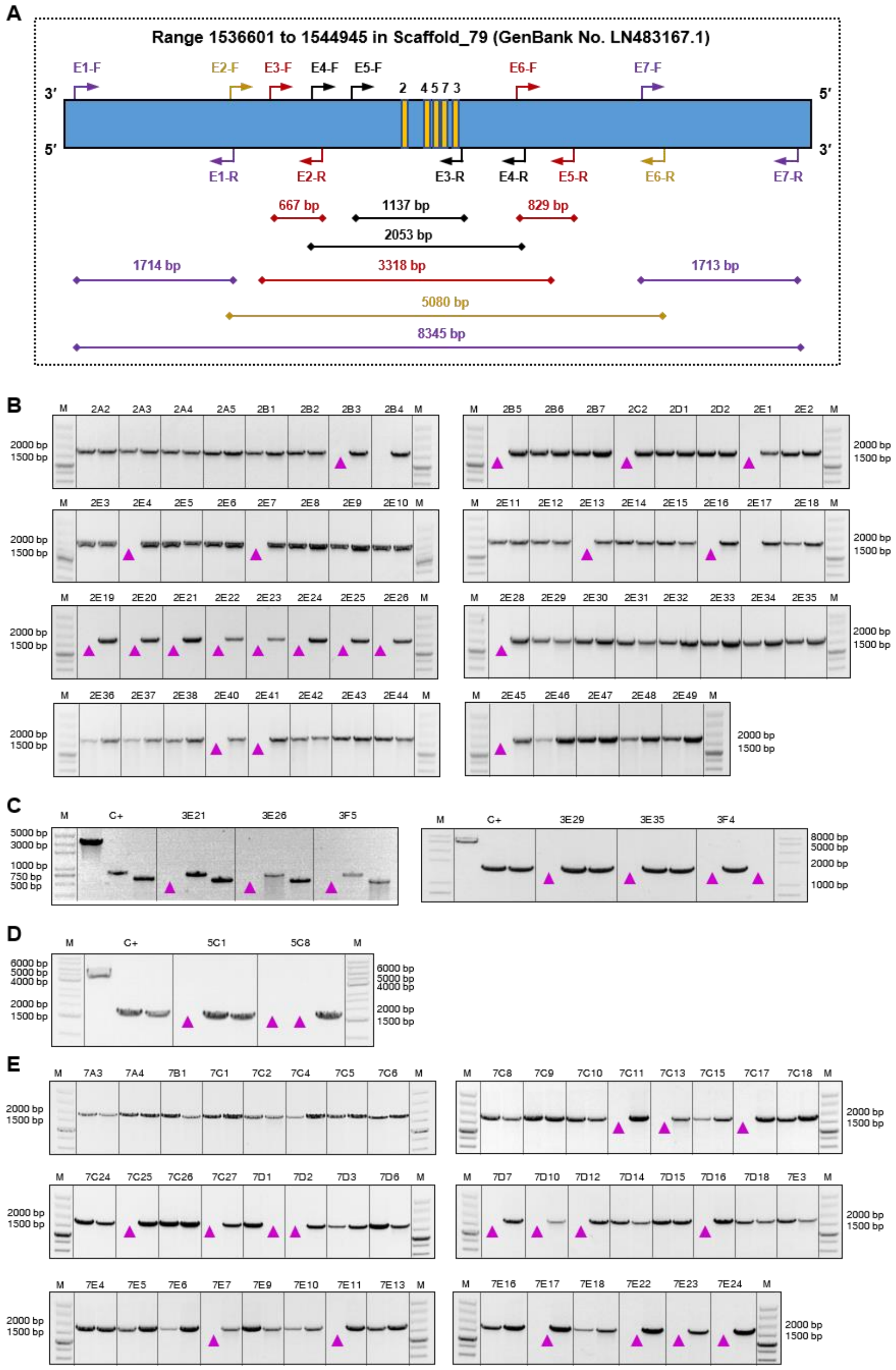


**Figure S2. Characterization by PCR amplification of the DNA repair patterns at *CrtE* sites in edited clones.** (**A**) Primer pairs and *CrtE* target sites. (**B** to **E**) Gel pictures showing amplification of upstream and downstream sequences of the target sites 2 (B), 3 (C), 5 (D), and 7 (E), and DNA regions spanning the target sites (C and D). The DNA regions that could not be amplified do not have bands, as indicated with purple triangles. M, DNA marker; C+, positive control.

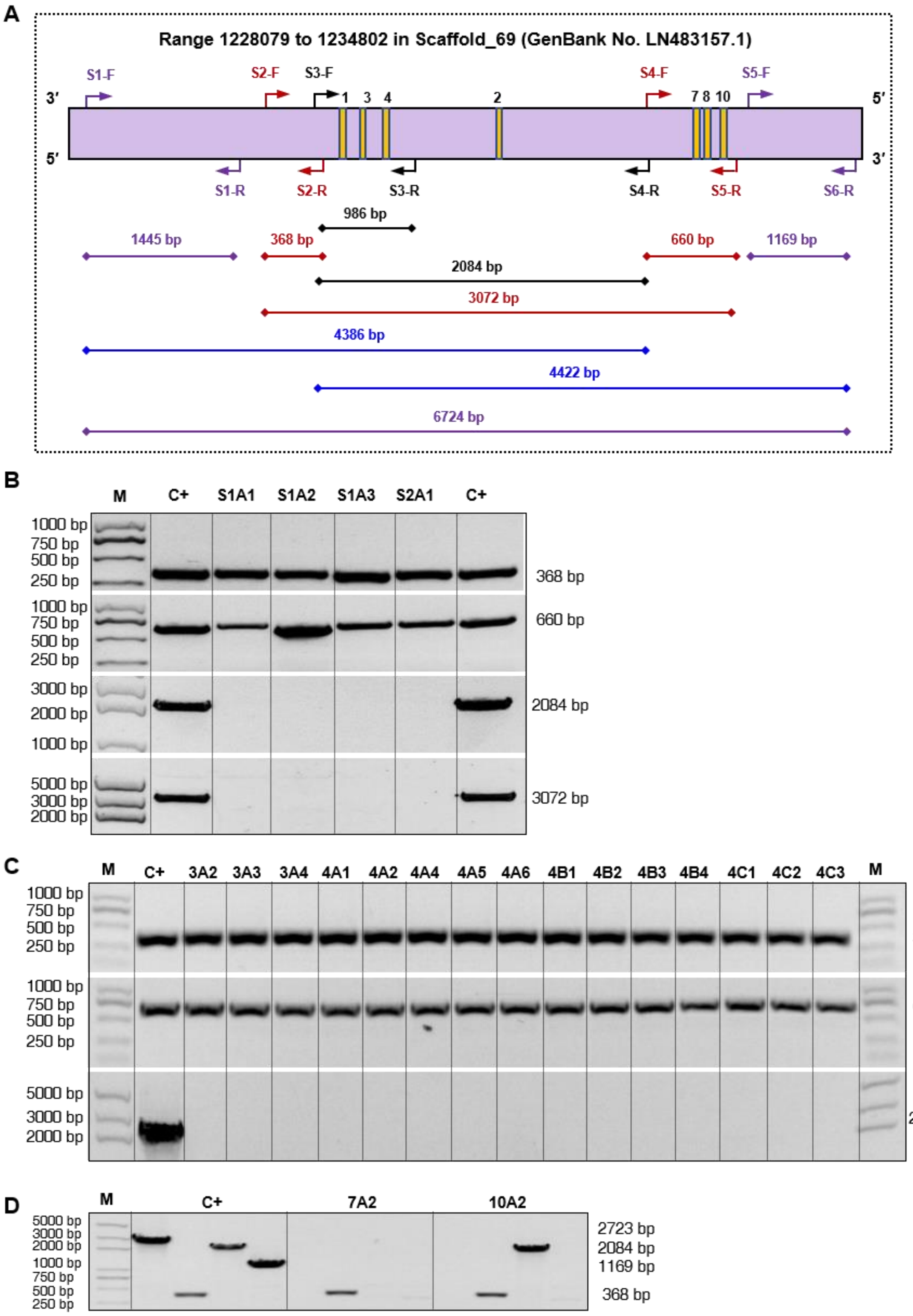


**Figure S3. Characterization of the DNA repair patterns at *CrtS* sites in edited clones by PCR amplification.** (**A**) Position of primer pairs and *CrtS* target sites. (**B** to **D**) Gel pictures showing the amplification of upstream and downstream sequences of the target sites and DNA regions spanning the target sites. *CrtS* site 1 and 2 (B); *CrtS* site 3 and 4 (C); *CrtS* site 7 and 10 (D). M, DNA marker; C+, positive control.

A

| Site | Codon | Amino acid | Colony color |
|---|---|---|---|
| Val88 (GUG) | GAG | V88E | White |
| | GCG | V88A | Red |
| | GGG | V88G | Red |
| | CUG | V88L | White |
| | AUG | V88M | White |
| | UUG | V88L | White |
| Ala89 (GCC) | CCC | A89P | Red |
| | ACC | A89T | Red |
| | UCC | A89S | Red |
| | GAC | A89D | White |
| | GUC | A89V | White |
| | GGC | A89G | Red |

B

| Site | Codon | Amino acid | Colony |
|---|---|---|---|
| Arg84 (CGU) | CAU | R84H | White |

D

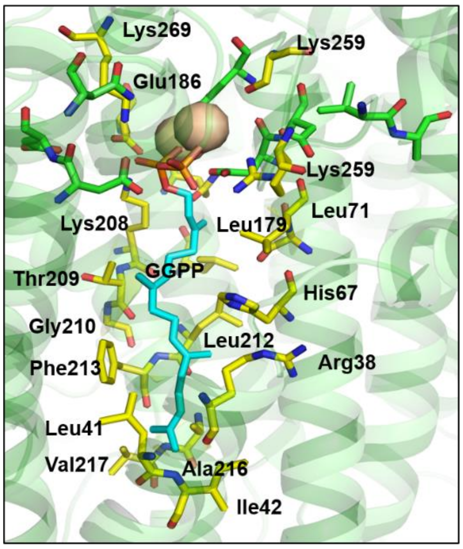


C

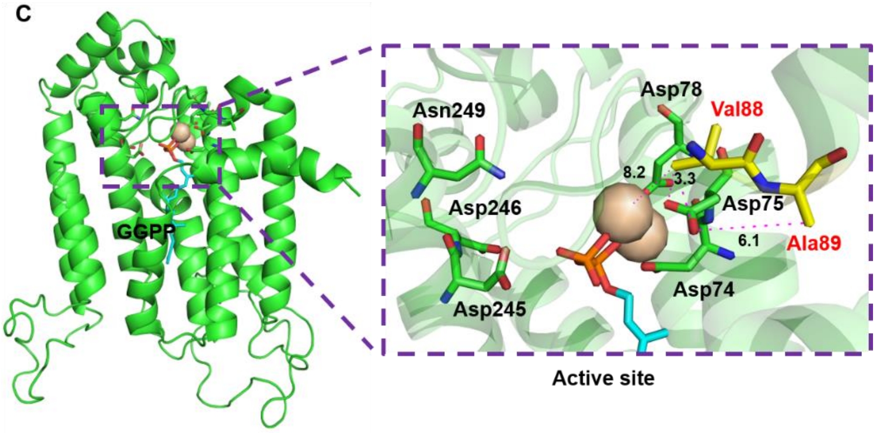


**Figure S4. Analysis of the amino acid residues affecting the activity of CrtE and colony color.** (**A** and **B**) Mutation analysis of Val88 and Ala89 in the *CrtE* target site 3 (A) and Arg84 in the *CrtE* target site 7 (B). The blue letters denote the mutated nucleotides. Red text indicates on-target point mutations identified by Sanger sequencing in CRISPR-Cas9 edited clones. (**C**) Modeling of CrtE based on structural homology and showing the active sites. The residues of active sites combined with the substrate GGPP (cyan, carbons; red, oxygens; orange, phosphorus) or magnesium (wheat spheres) shown in black labels. Residues Val88 and Ala89 are located near the active sites and shown as yellow sticks in the diagram. (**D**) Residues in the catalytic tunnels of CrtE within 4 Å of the substrate GGPP (cyan). GGPP, geranylgeranyl diphosphate. The target *CrtE* sites E4-E9 (Fig. S1D) were designed according to the modeled active sites.

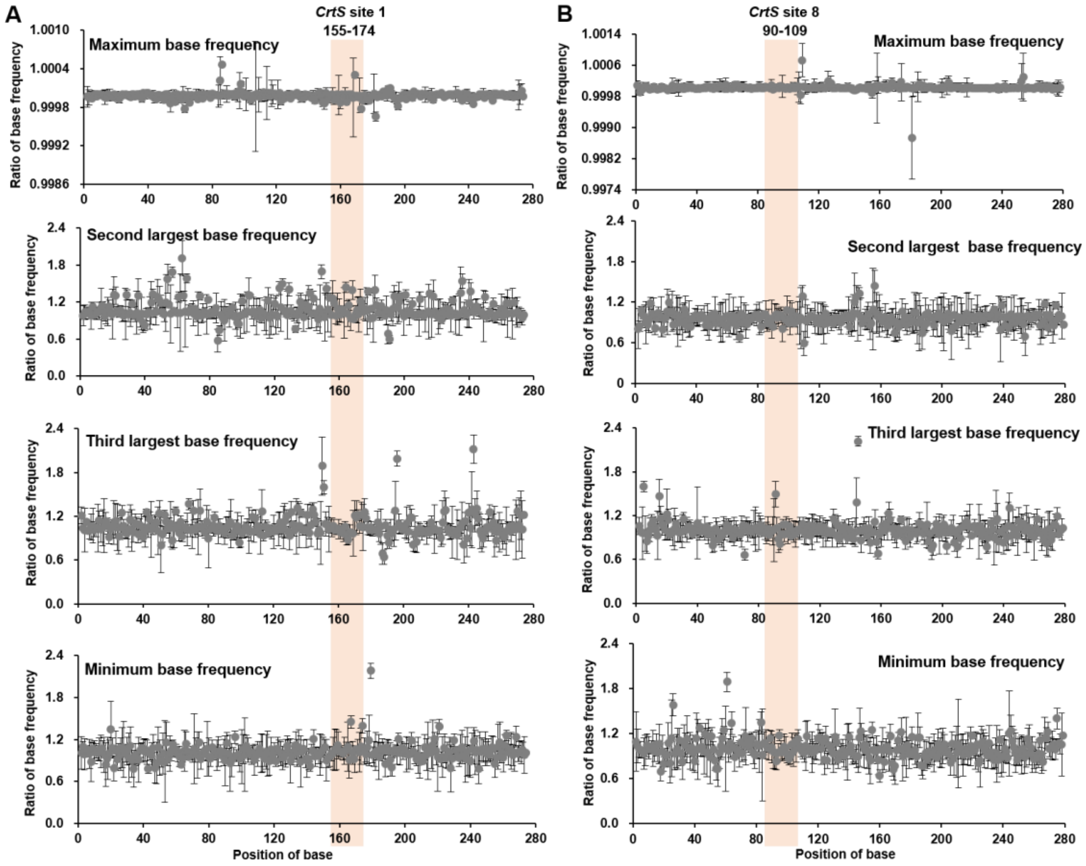


**Figure S5. Identification of on-target nucleotide substitutions in DNA repair induced by CRISPR-Cas9 cleavage using deep sequencing.** The ratio of base frequency of edited clone to control is shown for the four possibilities. The ratio of base frequency at CrtS site 1 (**A**) and CrtS site 8 (**B**) calculated from data from sequencing the corresponding DNA region for five technical repeats for each site. The highlighted area denotes the 20-bp target sequence.

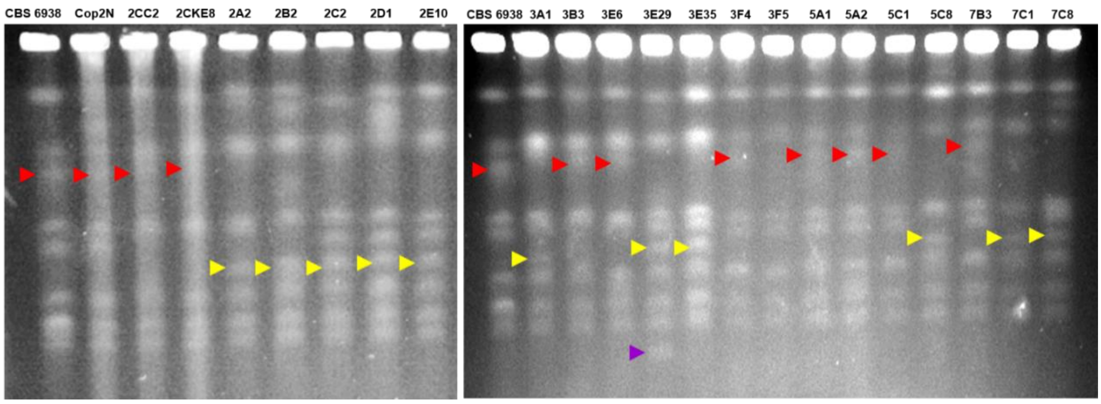


**Figure S6. Chromosome karyotypes detected by pulsed-field gel electrophoresis (PFGE).** The yellow triangles indicate the new chromosomes of equal size that appeared in some of the edited clones. The red triangles indicate chromosomes in the unedited clones and those with deletion or nucleotide substitution repair patterns that were lost in some of the edited clones in which PCR amplification failed to obtain amplicons spanning the target sites. The purple triangle represents a small chromosome generated in the edited clone 3E29 (Fig. S2C). CBS6938, the wild type strain; Cop2N, a strain expressing Cas9 but not gRNA. The lanes labeled 2CC1 and 2CKE8 are normal red clones growing on selective plates in gene editing experiments. Other samples are clones edited at *CrtE* sites 2, 3, 5, and 7.

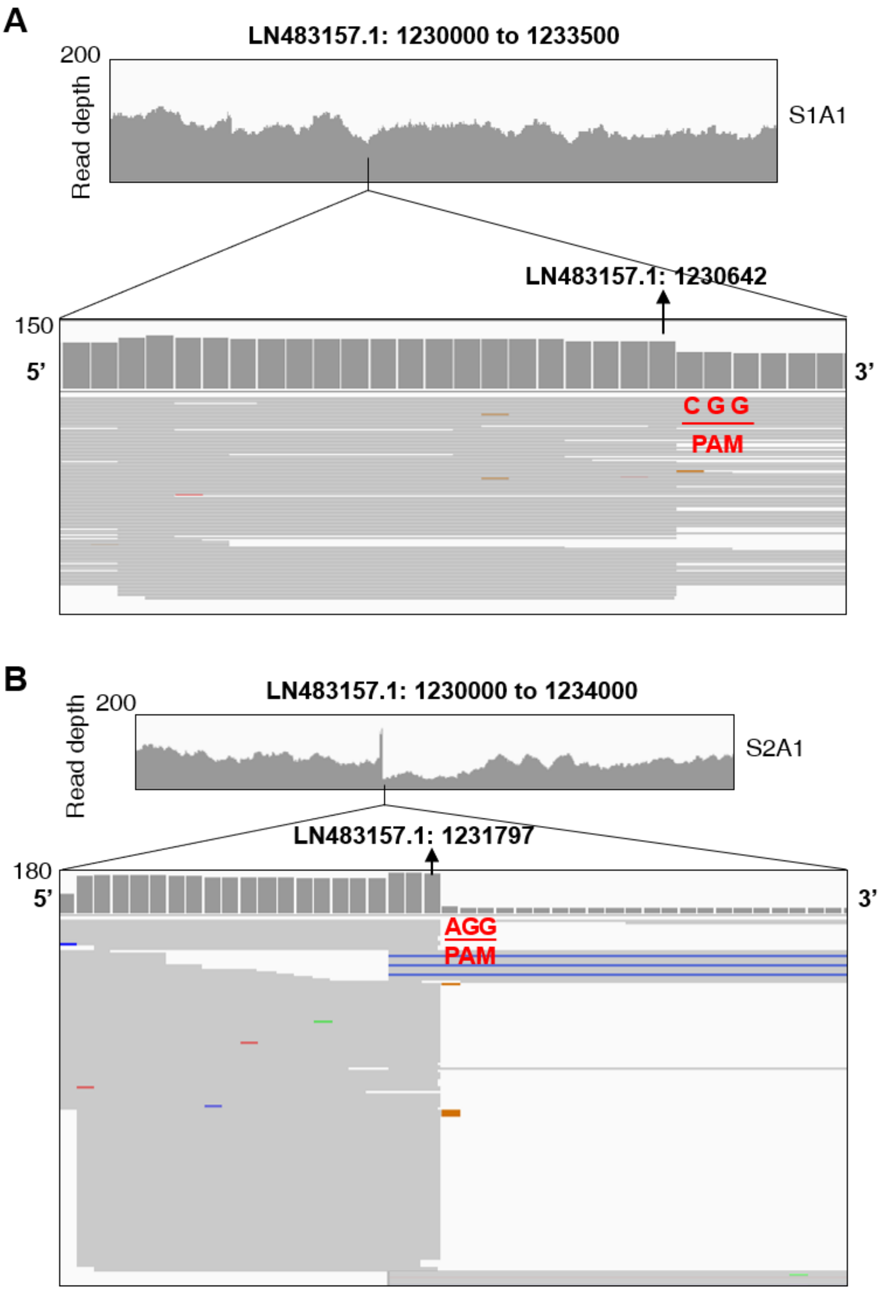


**Figure S7. Analysis of genome sequences showing the break sites in chromosome rearrangement induced by CRISPR-Cas9 cleavage in the edited clones S1A1(A) and S2A1 (B).** Structural variations (SVs) predicted by BreakDancer software package indicated chromosome rearrangement in the two strains, with no large DNA deletions near the PAM region. The arrows denote the possible break positions. PAM, protospacer-associated motif.

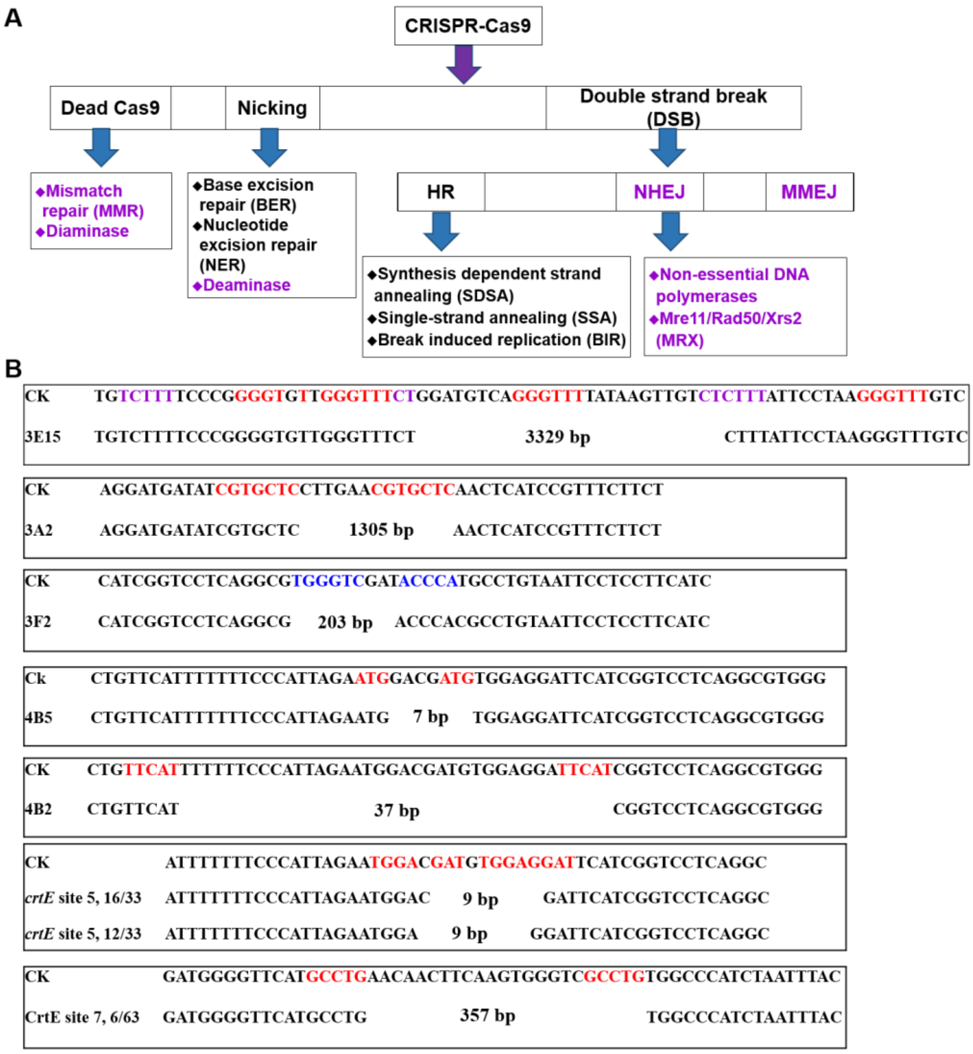


**Figure S8. Pathways potentially associated with DNA repair after CRISPR-Cas9 cleavage.** (**A**) Three potential pathways are suggested by the activity of Cas9. Dead Cas9 does not cut DNA, but can bind DNA. DNA nicking by Cas9 results in a single-strand DNA break or CRISPR-Cas9 cutting leads to DNA DSB. The influences of the pathways in purple were investigated in this study. HR, homologous recombination; NHEJ, non-homologous end joining; MMEJ, microhomology-mediated end joining. (**B**) Representative deletions induced by repair of CRISPR-Cas9 cleavage. Sizes of deletions are shown. Red and purple texts indicate the direct repeats. Blue texts represent reverse repeated sequences. CK, control. The samples 3A2, 3E15, 3F2 and 4B2, 4B5 are white mutants identified from editing *CrtE* site 3 and Site 4, respectively. The numbers 16/33, 12/33, and 6/33 represent the ratio of the total number of deletion repair to that of white edited clones.

**Table S1. Chromosomal translocations induced by repair of Cas9 cutting**

| Strain | Scaffold | Reads | SV types | Start position |
|---|---|---|---|---|
| ***CrtE* site 2** | | | | |
| 2A2 | LN483144.1 | 19 | CTX_after | 161779 |
| | LN483167.1 | 19 | CTX_before | 1539137 |
| 2B2 | LN483144.1 | 24 | CTX_after | 161779 |
| | LN483167.1 | 24 | CTX_before | 1539137 |
| 2C2 | LN483144.1 | 39 | CTX_after | 161779 |
| | LN483167.1 | 39 | CTX_before | 1539137 |
| 2D1 | LN483144.1 | 26 | CTX_after | 161778 |
| | LN483167.1 | 26 | CTX_before | 1539136 |
| 2E10 | LN483144.1 | 28 | CTX_after | 161779 |
| | LN483167.1 | 28 | CTX_before | 1539137 |
| ***CrtS* sites** | | | | |
| S1A1 | LN483144.1 | 12 | CTX_after | 161778 |
| | LN483157.1 | 12 | CTX_before | 1229641 |
| S2A1 | LN483144.1 | 47 | CTX_after | 161779 |
| | LN483157.1 | 47 | CTX_before | 1230796 |
| S3A2 | LN483144.1 | 43 | CTX_after | 161779 |
| | LN483157.1 | 43 | CTX_before | 1229813 |
| S4A1 | LN483144.1 | 55 | CTX_after | 161779 |
| | LN483157.1 | 55 | CTX_before | 1230110 |
| S4A2 | LN483144.1 | 61 | CTX_after | 161779 |
| | LN483157.1 | 61 | CTX_before | 1230110 |
| | LN483157.1 | 4 | ITX_after | 2504540 |
| | LN483157.1 | 5 | ITX_before | 1230307 |

**Denote**: CTX, interchromosomal translocation; ITX, intrachromosomal translocation. SV, structural variation.

**Table S2**. **Gene-disrupted yeast strains used in this study.** The CRISPR-Cas9 edited clones were shown in Supplementary Data S1.

| **Strains** | **Genotype** |
| --- | --- |
| CBS 6938 | Wild type, CBS collection |
| MSH2Δ | *MSH2::Zeocin* |
| MLH1Δ | *MLH1::Zeocin* |
| KU70Δ | *Ku70::Zeocin* |
| KU80Δ | *Ku80::Zeocin* |
| Mre11Δ | *Mre11::Zeocin* |
| RAD50Δ | *Rad50::Zeocin* |
| Sae2Δ | *Sae2::Zeocin* |
| REV1Δ | *Rev1::Zeocin* |
| REV4Δ | *Rev4::Zeocin* |
| Pol4Δ | *Pol4::Zeocin* |
| RAD30Δ | *RAD30::Zeocin* |
| REV4Δ & Pol4Δ | *Rev4::Zeocin, Pol4::Hgr* |
| REV4Δ & RAD30Δ | *Rev4::Zeocin, RAD30::Hgr* |
| Pol4Δ & RAD30Δ | *Pol4::Hgr, RAD30::Zeocin* |
| ADA1Δ | *ADA1::Zeocin* |
| CDA1Δ | *CDA1::Zeocin* |
| CDA2LΔ | *CDA2L::Zeocin* |
| GUD2Δ | *GUD2::Zeocin* |
| FCY1-2Δ | *FCY1-2::Zeocin* |

### Captions for Supplementary Data S1 to S4

#### Data S1

**Primers and plasmids used in this study.** The primers used for vector construction by Gibson assembly are shown. The characteristics of target plasmids are described.

#### Data S2

**Characteristics of the DNA sequences used in this study.** The sequences for construction of CRISPR-Cas9 system, gRNA sequences of CRISPR-Cas9 system, and homologous arms for gene disruption are shown.

#### Data S3

**CRISPR-Cas9 edited and pigment-changed clones.** The red and yellow clones generated by CRISPR-Cas9 editing at genes *CrtE* and *CrtS* are listed. The repair patterns of point mutation and deletion are shown.

#### Data S4

**Deletion sequences generated in repair of Cas9 cleavages.** Sizes of deletions are shown. Direct repeats and reverse repeated sequences are represented in red and blue texts.
