## Supplementary Data S2 for "The cause of on-target point mutations generated by CRISPR-Cas9 treatment in the yeast *Xanthophyllomyces dendrorhous*"

**Section 1: Sequences used in construction of the CRISPR-Cas9 system**

**> Codon optimized Cas9**

atgcctaagaagaagagaaaggtcgacaagaagtactccatcggactcgacatcggaaccaactctgtcggatgggccgttatcaccgacgagtacaaggtcccttccaagaagttcaaggtcctcggaaacaccgaccgacactccatcaagaagaacctcatcggagccctcctctttgactctggagagactgccgaggctaccagacttaagcgaactgcccgaagacgatacacccgacgaaagaaccgaatctgctacctccaggagatcttctccaacgagatggccaaggtcgacgactccttcttccaccgactcgaggagtccttccttgtcgaggaggacaagaagcacgagcgacaccctatcttcggaaacatcgtcgacgaggtcgcctaccacgagaagtaccctaccatctaccacctccgaaagaagctcgtcgactccaccgataaggctgacctccgactcatctatcttgccctcgcccacatgatcaagttccgaggacacttcctcatcgagggtgacctcaaccctgacaactccgacgtcgacaagctcttcatccagctcgtccagacctacaaccagctcttcgaggagaaccctatcaacgcctccggagtcgatgccaaggctattctctccgcccgactctctaaatcccgacgactcgagaaccttatcgcccagctccctggagagaagaagaacggactcttcggaaacctcatcgccctctcccttggacttacccctaacttcaagtccaacttcgaccttgccgaggacgctaaactccagctctccaaggacacctacgacgacgacctcgataatctcctcgcccagatcggagatcagtacgccgacctttttctcgccgccaagaatctttccgacgccatccttctctccgacatcctccgagtcaacaccgagatcaccaaggctcctctctccgcctccatgatcaagcgatacgacgagcaccaccaggatctcaccctcctcaaagccctcgtccgacagcaactccctgagaagtacaaggagatcttcttcgaccagtccaagaacggatacgccggatacatcgatggaggagcctcccaagaggagttctacaagttcatcaagcctatcctcgagaagatggacggaaccgaggagcttctcgtcaagctcaaccgagaggacctcctcagaaagcagcgaaccttcgacaacggatccatccctcaccagatccaccttggagagctccacgccattctcagacgacaggaggacttctaccctttcctcaaggacaaccgagagaagatcgagaagatcctcaccttccgaatcccttactacgtcggacctctcgctcgaggaaattcccgattcgcctggatgacccgaaaatccgaggagactatcaccccttggaacttcgaggaggtcgtcgacaaaggagcctctgcccagtctttcatcgagcgaatgaccaacttcgacaagaacctccctaacgagaaggtcctccctaagcactccctcctctacgagtacttcaccgtctacaacgagctcaccaaggtcaagtacgtcaccgagggaatgcgaaagcctgccttcctctccggagaacagaagaaggccatcgtcgacctcctcttcaagaccaaccgaaaggtcaccgtcaagcagctcaaggaggactacttcaagaagatcgagtgcttcgactccgtcgagatctctggagtcgaggacagattcaacgcctccctcggaacctatcacgacctcctcaagatcatcaaggacaaggacttcctcgacaacgaggagaacgaggacatcctcgaggacatcgtcctcaccctcacccttttcgaggaccgagagatgatcgaggagcgactcaagacttacgcccacctcttcgacgacaaggtcatgaagcagctcaagcgacgacgatacaccggatggggacgactttcccgaaagctcatcaacggaatccgagacaagcagtccggaaagaccatcctcgacttcctcaagtccgacggattcgccaaccgaaacttcatgcagctcatccacgacgactccctcaccttcaaggaggacatccagaaggcccaggtttctggacaaggagactccctccacgagcacattgccaacctcgctggatctcctgccatcaagaagggaatcctccagaccgtcaaggtcgttgacgagctcgtcaaggtcatgggacgacacaagcctgagaacatcgtcatcgagatggcccgagagaaccagactacccagaagggacagaagaactcccgagagcgaatgaagcgaatcgaggagggaatcaaggagctcggatcccagatcctcaaggagcaccctgtcgagaatacccagctccagaacgagaagctctacctctactacctccagaacggacgagacatgtacgtcgaccaggagctcgacatcaaccgactctccgactacgacgtcgaccacatcgtccctcagtccttcctcaaggacgactccatcgacaacaaggtcctcacccgatccgacaagaaccgaggaaagtccgacaacgtcccttctgaggaggtcgtcaagaagatgaagaactactggcgacagctcctcaacgccaagctcatcacccagcgaaagttcgacaacctcaccaaggccgaacgaggaggactttccgagcttgacaaggccggattcatcaagcgacagctcgtcgagactcgacagatcaccaagcacgtcgcccagattctcgactcccgaatgaacaccaagtacgacgagaacgacaagctcatccgagaggtcaaggtcatcaccctcaagtccaagctcgtttccgacttccgaaaggacttccagttctacaaggtccgagagatcaacaactaccaccacgcccacgacgcttatcttaacgccgtcgtcggaaccgctctcatcaagaagtaccctaagctcgagtccgagttcgtctacggagactacaaggtctacgacgtccgaaagatgatcgccaagtccgagcaggagatcggaaaagccaccgccaagtacttcttctactccaacatcatgaacttcttcaagaccgagatcaccctcgccaacggagagatccgaaagcgacctctcatcgagactaacggagaaaccggagagatcgtctgggacaagggaagagacttcgccaccgtcagaaaggtcctctccatgcctcaggtcaacatcgtcaagaagaccgaggtccagaccggaggattctccaaggagtccatcctccctaagcgaaactccgacaagctcatcgcccgaaagaaggactgggaccctaagaagtacggaggattcgactcccctaccgtcgcttactccgttctcgttgtcgccaaggtcgagaagggaaagtccaagaagctcaagtccgtcaaggagctcctcggaatcaccatcatggagcgatcctccttcgagaagaaccctatcgacttcctcgaggccaagggatacaaggaggtcaagaaggacctcatcatcaagctccctaagtactccctcttcgagctcgagaacggacgaaagcgaatgcttgcctctgccggagaactccagaagggaaatgagctcgccctcccttccaagtacgtcaacttcctctacctcgcctcccactacgagaagctcaagggatcccctgaggacaacgagcagaagcagctcttcgtcgagcagcacaagcactacctcgacgagatcatcgagcagatctccgagttctccaagcgagtcatcctcgccgatgctaacctcgacaaggtcctctccgcctacaacaagcaccgagacaagcctattcgagagcaggccgagaacatcatccacctcttcaccctcactaacctcggagctcctgccgccttcaaatacttcgacaccaccatcgaccgaaagcgatacacctccaccaaggaagttctcgacgccaccctcatccaccaatccatcaccggactctacgaaacccgaatcgacctctctcagctcggtggagaccctaagaagaagagaaaggtctaa

**>sgRNA**

tgtcgttagtgtcatcggattgatctctatcccaggcctgatgacaggagcgatcatcggtggttcttccgtcgaacaagccgcgaaattacaaagtacatctcgaaatttcgatactttgaactttgcctttaaaacttttagatcgactgatatactcttcgtttcctttaccaaaagtgatcttgatgtttatgatcagcgcttcttctgcactatccggttagtccttcttcctcctatcgatcgcctagtcccaatcagaactcctttgaattgactcctgatcctttgttttccaccttcaactctcgccaatagtgctggcagccatgatattcacattctcggtcgtattcgaccctcaacatcgaatcagagacgacaggatatattcgtccgagagcgtgctttctggtgggatcagaagtgttgggaagatcggcgtgttgggttggaaatcgatcaaaaatctaggaggaaaggagcgaaggccatgagagtaagaacgtggggacctggcgtagtggtagcgcccagcctttgcgtggctgtagccccggttcgattccgggggtctccaaccccgtaaattagatgggccacgttttagagctagaaatagcaagttaaaataaggctagtccgttatcaacttgaaaaagtggcaccgagtcggtgcttttttt

**Section 2: Sequences of CRISPR-Cas9 target sites and primers**

**>CrtE_LN483167.1 (1539235 to 1541284)**

atggattacgcgaacatcctcacagcaattccactcgagtttactcctcaggatgatatcgtgctccttgaaccgtatca

ctacctaggaaagaaccctggaaaagaaattcgatcacaactcatcgaggctttcaactattggttggatgtcaagaagg

aggatctcgaggtcatccagaacgttgttgggtatgtcatctctcttagcttcttgtcggttggggttgttgcttggagc

agaacggagttagtagaccgagcaggtcaaacgccagatgaggttcatgcctgaacaacttcaaccctcggcacggaacaatgcagcactgacggtcgttatttcgttctttctcgttcttgttctcgttggtaatccgattcgacttgcttggtctgct

ataaacagcatgctacataccgctagcttattgtgagtcttttcctcatcctcttgcctttctacacatctcccatcatg

ctggaccactggtcgtctagttacaacagtcctaaagatgcgacggtaaactgacatctttctcttgtgcgtttgctccg

tcatttgtgtaaacgtcccttctgttcatttttttcccattagaatg**gacgat**gtggag**gat**tcatcggtcctcaggcgt

gggtcgcctgtggcccatctaatttacgggattccgcagacaataaacacgttcgttcatttgctcattccatcccatct

ctgtgtgtgttccgaaagtcctgacttgtctattctgaccggataaccactatggattatagtgcaaactacgtctactt

tctggcttatcaagagatcttcaagcttcgcccaacaccgatacccatgcctgtaattcctccttcatctgcttcgcttc

aatcatccgtctcctctgcatcctcctcctcctcggcctcgtctgaaaacgggggcacgtcaactcctaattcgcagatt

ccgttctcgaaagatacgtatcttgataaagtgatcacaggtaagcttcacttgtactctagtctttcgattatatgtat

tcatctgaccagcattcgattcttcgggtaaatagacgagatgctttccctccatagagggcaaggcctggagctattct

ggagagatagtctgacgtgtcctagcgaagaggaatatgtgaaaatggttcttggaagtgagcgaatggtcttttgtcct

ggctgaaatcattgttgtctattgctgaccatgttcgggatgaacgtccatctagagacgggaggtttgttccgtatagc

ggtcagattgatgatggcaaagtcagaatgtgacatgtatgtgctcaataataactacatccgtctatgttttcttgtac

agcataacgtgctcaactcatccgtttcttctgggaacccttaatcgtactgggcacatatcattgtgtagagactttgt

ccagcttgtcaacttgatctcaatatacttccagatcagg**gatgac**tatatg**aac**cttcagtcttctgaggtacttatcg

tttctctgagtatcacagtcgaaacaatctattctaatgtctcctggtgctttgactgcaaacttttatcagtatgccca taataagaattttgcagaggacctcacagaaggaaaattcagttttcccactatccactcgattcatgccaacccctcat

cgagactcgtcatcagtcagtttcttctcttcatttctcaccatgattcatactctatttgtagatgttgagcttgtcat

tcctctttctctttcggcgcgctatagatacgttgcagaagaaatcgacctctcctgagatccttcaccactgtgtaaac

tacatgcgcacagaaacccactcattcgaatatactcaggaagtcctcaacaccttgtcaggtgcactcgagagagaact

aggaaggcttcaaggagagttcgcagaagctaactcaaagattgatcttggagacgtagagtcggaaggaagaacggggaagaacgtcaaattggaagcgatcctgaaaaagctagccgatatccctctgtga

**>CrtS_LN483157.1 (1230024 to 1233189)**

atgttcatcttggtcttgctcacaggtgctttaggcctggctgctttctcatgggcatccatagcgttcttcagtcttta

cctcgctccgaggcgatcttcactgtataaccttcagggtaagaattgagctctggaatcatgcttgtgtaaatcctata

atctcattcatcctattcctcttcttcatcctctcttcaggcccgaatcataccaactactttacaggcaattttttaga

catcctctcgtgagttttcatcattggctcagtcgtccaatcttaacgatcatcgctaacgacctttcggacgcgttctt

ctttctatgtgaaatctgatctttggtttgttacgagagcacagagctcgtacaggtgaagagcatgcgaagtacagaga

aaaatacggaagcaccctccggtttgctgggatcgctggagcacccgtcttgaactcgaccgatccgaaagtcttcaacc

agtttgtccatccgaaccctcatcctcctctgctgatcaattcaactgtagttaacgcactttgaatggacagtgtgatg

aaagaagcctacgactatccgaaacctggtatggccgctcgagtgctcagaattgctaccggagatggtgttgttacggc

ggaaggtgcttttcaagttctcttatatcacatctaatccactcggcgcgattgaactcaacatttctgacgagcctgtc

accttgttttcacttcatggtctcggtgcatcttgtctcatctcataggtgaagctcataagcgacatcgaaggatcatg

atcccctctctgtccgctcaggccgttaagtcgatggtcccaattttcttagaaaaaggtatggaacttgtcgacaagat

gatggaggatgcggctgagaaggatatggccgtgggagagtcggccggtgaaaagaaggcaaccagactcgagaccgaaggagtcgatgtaaaggattgggtcgtgagtacccgcctattccttcaccttgatggacgaagcatatcaaggaaaggttcattgactgacaaacactatcttaccagggtcgagctactctggacgtcatggctcttgcaggtcagtctactctctcttat

aaatgctccacatatgtatgcatgtactgacatgctcttcctatattcgatacgacgtcatatgtccaggatttgactat

aagagcgactcgctccagaacaagaccaatgagctctatgtcgcttttgtcggacttaccgatgggtttgctcctacctt

ggactcgttcaaggctatcatgtgggattttgtaccttacttccgaactatggtatgtctgccattctttgatatccaaa

gattatggataggttacttgctaaaatttcacctatcgtgaacagaaacggagacatgagatacctttgactcaaggatt

agcagtttcccgacgagttgggatcgtaagtgccagatcaagcctctctgaatattcttggtcatcatcttaacctccta

ggctcattcatccatggtgcgcaataggagcttatggagcaaaagaagcaggccgtgcttggctcagcttccgatcaggc

tgttgataaaaaggatgttcaaggtcgggatatcctaagtctcctaggttagtaacgtttttaaacgtatatacagagcg

gcgacattctttccctgacaactgtcaacatgctcgttactagtgagagcaaacatcgccgccaacctgcctgaatctca

aaagctgtccgatgaggaggtactcgctcagatcagtaacctgttatttgctggatatgagtgtgtatcctttcccctct

ctatccttagctgattaaaagcactaatagaggtctttatgtttcctgtttgatcagaacttcttcgacagtcttgacat

ggatgtttcaccgactctcagaagacaaagccgttcaggataaacttcgagaagaaatttgtcagatcgacacggatatg

cctacgctgtgaggatgtttttgatgctaaattacttcttcttgcaaatgactaaaacggccttccattcttgatccatt

ttagagacgaacttaatgcgttgccttatctcgaagcggttggttctcgattcttggtcttgtcttccaaatacaatacg

gattattgctcatctgatttgcgtctacgggctgtggaatttaactagtttgttaaggagtctcttcgtctagaccctcc

tagtccgtatgctaaccgtgaatgcttaaaggatgaagacgtatgttggcttcatcacgcataattttcatttcatattc

ctttgtacatacgcatacaggctgaccgagctcaaattccggcttcctcttctgtgcttctttttctggcctttcttatc

ttcattcttcaaccaaaatttgtcacagttcatcccacttgccgagcctgtcattggtcgagatgggtcggtcatcaacg

aggtccggatcacgaaaggaacgatggtcatgcttcgtaagttttcctttatttcatctcgtccatgaaatagtttctga

tagacgcggaccaattcagcgttgttcaacatcaatcgttcaaagttcatttatggagaagatgcagaagaattcaggta

caattcgttttcttttaaaagccaatcggtttcgtatcgtaattgaccgggctctcttttaatttctcgaaagaccggag

aggtggcttgaggacgtaacagactcgctcaacagtattgaagcaccctatggacaccaggcgagctgtatgttttattg

attttatctttgtgaattttgcaaaacgttgaacttcgcgcttcccttgttgttgaaatcccagttatctctggacccag

agcttgcttgtaagtttcttctcatctggcgccttagcagtatccgatcagccatctagttctttgtacgattgtttctg

actctctcgactttcgcagtggttggcgatttgctgtcgccgagatgaaggccttcttgtttgtcactctccgtcgggtc

cagttcgagcccatcatctctcatccagagtacgagcacatcaccttgatcatttcccgtcctcgaatcgttggtagaga

gaaggaggggtaccagatgcgtttgcaggtcaagccggtcgaatga

**Section 3: Sequences for disruption of genes by homologous recombination and primers for diagnostic PCR.**

> CrtE gene and up down

tcgccagattcgttagatgtagctcaggtgagtgaataatatttctacaattgaataacacattttccttctttttctcg

ctctgtatactgacctagagagggttttgtgcctcatggcagctccgaggggttactgtatacgaaggctgctcggcaga

agatgagttgattgagtaagggcttaattttcctcttcgaccttgccttc

tttcaccggaagtagctcatctcctgttggctgtcttcaacgatctgtgagtgtagatcgttctgggagctcgtctctgcttggtcgaccgaacaacaacaaaagcttctggcgttcgttgtacgtatgatgtttattttagctttatcgacgaatgcatctttgacgatttgaaatcggttgtcttcgc

ttatgacatcagacggggagcgacagagtgcctgcgaccgggctgatcaacatgagcttccgaattcaagctaccagagt

gcccgagacttatcttccctgtacgtgtcttgagcttgaactctgaaccgacctagatatcgcatcctgaccatttgttg

ttcaactatgatcacttccctcctcgaaagcttcgcacacatgtttcaatacgctctgcttgcctaggtatagatcaaag

acagtgcttcagaaaaagcttgaagaggcgctcacgatgagccagggtttcgggctcaaatagatatcttgttggcctgt

cattgtatccattgtttctttattctcctatctctataattatctgtttctgtaccagactgttcatacatgtccatcgt

gtcgatatatagaatccctttttaatgactcttgccggcgatttgacttacgtaattgcgctcaatccggcggtcgaaat

ctctttgcccgagcccgcccggttcggaggtggttcgacctggttcgatggccggcgacttggccgatcggcacggcctt

gcgtgttctcctggtttctccatctcgctcgatcgtttttcatctggttcggaagatacgatgaaccagaaggatactcg

accagcatataccccactcgttcatcgcacacgtagcatacacccaatttaaagtgcactcagccatagctaacacacag

aactacacatacatacactcatccggaacacatagg (**upstream homologous arm fragment**)

atggattacgcgaacatcctcacagcaattccactcgagtttactcctcaggatgatatcgtgctccttgaaccgtatcactacctaggaaagaaccctggaaaagaaattcgatcacaactcatcgaggctttcaactattggttggatgtcaagaaggaggatctcgaggtcatccagaacgttgttgggtatgtcatctctcttagcttgtcggttggggttgttgcttggagcagaacggagttagtagaccgagcaggtcaaacgccagatggggttca

tgcctgaacaacttcaaccctcggcacggaacaatgcagcactgacggtcgttatttcgttctttctcgttcttgttctc

gttggtaatccgattcgacttgcttggtctgctataaacagcatgctacataccgctagcttattgtgagtcttttcctc

atcctcttgcctttctacacatctcccatcatgctggaccactggtcgtctagttacaacagtcctcaagatgcgacggt

aaactgacatctttctcttgtgcgttcgctccgtcatttgtgtaaacgtcccttctgttcatttttttcccattagaatg

gacgatgtggaggattcatcggtcctcaggcgtgggtcgcctgtggcccatctaatttacgggattccgcagacaataaa

cacgttcgttcatttgctcgttccatcccatctctgtgtgtgttccgaaagtcctgacttgcctattctgaccggataac

tactatggattatagtgcaaactacgtctactttctggcttatcaagagatcttcaagcttcgcccaacaccgataccca

tgcctgtaattcctccttcatctgcttcgcttcaatcatccgtctcctctgcatcctcctcctcctcggcctcgtctgaa

aacgggggcacgtcaactcctaattcgcagattccgttctcgaaagatacgtatcttgataaagtgatcacaggtaagct

tcacttgtactctagtctttggattgtatgtattcatctgaccagcattcgattcttcggttaaatagacgagatgcttt

ccctccatagagggcaaggcctggagctattctggagagatagtctgacgtgtcctagcgaagaggaatatgtgaaaatg

gttcttggaagtgagcgaatggtcttttgtcctggctgaaatcattgttgtctattgctgaccatgttcgggatgaacgt

ccatctagagacgggaggtttgttccgtatagcggtcagattgatgatggcaaagtcagaatgtgacatgtatgtgctca

ataataactacatccgtctatgttttcttgtagagcataacgtgctcaactcatccgtttcttctgggaacccttaatcg

tactgggcacatatcattgtgtagagactttgtccagcttgtcaacttgatctcaatatacttccagatcagggatgact

atatgaaccttcagtcttctgaggtacttatcgtttctctaagtatcacagtcgaaacaatctattctaatgtctcctgg

tgctttgactgcaaacttttatcagtatgcccataataagaattttgcagaggacctcacagaagggaaattcagttttc

ccactatccactcgattcatgccaacccctcatcgagactcgtcatcagtcagtttcttctcttcatttctcaccatggt

tcatactctatttgtagatgttgagcttgtcattcctctttctctttcggcgcgctatagatacgttgcagaagaaatcg

acctctcctgagatccttcaccactgtgtaaactacatgcgcacagaaacccactcattcgaatatactcaggaagtcct

caacaccttgtcaggtgcactcgagagagaactaggaaggcttcaaggagagttcgcagaagctaactcaaggatggatc

ttggagacgtagattcggaaggaagaacggggaagaacgtcaaattggaagcgatcctgaaaaagctagccgatatccct

ctgtga

aagaacatattctctctctcgtctgtccgtttctatcagggttttataagttgtctctttattcctaagggttt

gtcagatgattggacttgatgtgctctattgcccgttcatctttttcacttcgacttttttctctaccgtgcatgcccat

tcgcattctcttgttcatcttgtgtttaatttgttcgacataacattaatcatcgtgtcttcttcttttcgaagaaatct

cgtgacttgttgaacttcaactataattaatcatattcatatctcaa

agtcttcgtcttctcgcaatgtgattcctccttccagttccctctttgatttccttctcattgatcggtttctttttcttttttgctctcctgtctcttctttattcgccttccgtctctctgtctcgttttctcttcacttttttttttcatcttctctcggtcaacttgtcatttaatctctctagggtct

catgtcaacacgtgccaagcatgtcatacgtgtgcagggtgatgtacagtcattttgccatccctcttcgcagggtctca

tctatcttgtctatcgacttttcctctttttgaatttcctcggagttttatcttggtataagcaatggagaagagcgcat

tcttccgttggtcgtgaactaaacacagacgacaataacagaaaggacattgaatatgttgcgtcgagctatacgagtga

tagaaattgatacaagatcggcggaaaccaggatgagaaaagaaagagagtgtctgcaaactcaatagagaaaagatgaacaactcgacgggtgcaatatagctggagaacagaagaatcagaatgctttctggccgatgctggagaaaaactttgcggc

gagcgctcggagaatctgaagagaaccaatggaaatcgagagataaaagtgagcaggataaggatgggcaggatcaacgagcgaaacttctttacctctgcctttcggtccgagccgagactctttgtttttccagtaaaagtggtattataagaacgat

tgtgctttggtcagtttccccctggagaaagaagaaagaagcatataaaggaacatagaaaacgtaccatgaagacctca

ccaacggcctttcctttagcatcccggacggcacctagacgatttccagcaaatcctctgaattgaaaagaaagaaaaga

agaattctaatcaagtcagcgctctttgttctatatgcacggcgatgatagtttactctc (**downstream homologous arm fragment**)

gaatccacagaacgtgagaacaaggtgatttagaataaaggacaactcgtacccggccatggctcctggtatgtttccaa

tcacaaatcctacagccgcacctcctgctcctcccaccaagctccacattcttcctgcgcgctctcttccaacctccggc

**>MSH2_ LN483345.1**

ggaactctgttagggttaccttctattttctccttctgttcatttgagatatttgacgcgtttccgacgcgacgcgtttc

tctacgcagccggtctttcaacgatttgtgagctttcacttt

cgatctcatagccatcttagtgcctcacacatactacgtattccttcttcatctgtttagagaagcaagatgagtctc

atgtatggaaaggagccagccggggagaagacatctcttgatatgggtaggttctgtgttttctcgagtgttctctcaac

aacaagctgacccaatgcgtcatatagataatgccgccgaggtcaactttatccgacaatttgagaatcttcccaaggtt

agttggtctggctactttttgatgatggagaaaaggtctcggcatttgtcgaccggttcttacaatccgataacacaata

gaaattagatggaacgattcgcctattcgaccgcgtcgtaagtgtctatcgtgtctttaatgctcaggagtgccagatca

ccggagttcatcgtctgttcaatgtctttacagaattactattcctgccacggagaagacgcaattcttatagcgagcat

cgtctacaacaccagtaatgctataaagttcatcggttctaaaaccaagaagtggccgttgggcttgccgagtgtgactc

tcaatcagaacgccgctaaggggtttctcagagatgctctcaccgccagacagatgaagatcgaggtcagtcaatctgtc

aagatgaaatgctctccaaatagcgtctgactgttcttttttatttttatttttttacccttgtcatgacagatttatag

tcaagagggaacaggaaggccaaatcagccttggcttcttactcgtcaagttagtctaaatatgactactaatcaacgat

gtcgcccgaactcaatagcggttgacctaatcattgatataacacataggcttctccgggaaacatatcgcaacttgaag

atcttcttttcaccaatcacgatctgctgacttctcccgtaaggctgtctttctgatcgactgctctggttcttcatact

tcaactgatatctgtgtgcactaaaccttctgaaattcagattgtcatggctatcaaggtgatgactaaggacggggttc

gcacagtcggggcagcctttgctgatgtatcaagtagagaacttggagtagcagaatttgccgataacgatctgttctcc

aatacagaggtctgtacgctttcctgttatacgatcaatatctcgtgtatcgagcaaaacctgatggtcttttctaatta

aaaatatcgtatgtctactacataacgacctattcagagcttgttaattcagctgagcgtcaaagaatgcatcatccagg

cggacgataaacgaacagac (**upstream homologous arm fragment**)

tatgatcttgcaaagatcagaacgatgctcgagcgatgtggggtaatcgctaccgaaacc

cgcgccagtgagttgtttaccatctctgtgccaagttgaacgtctgctgatacttagatcgtaaacaggtgaatttaatc

caaagaatgtagaacaagacctgaacagattactcaacccaaaacacgccgctcagtctcttcgtgagcgatcctgctaa

ttttgctctgttcattgaccatcgactaaaattctcttaatatgttcacgatatctcgttcctctttagctcaactcaac

ctcaagaccgcgctctactctgtctcagccctcatttcccatctctcactactgaccgatgctactacacacggaagctt

ctcattaagaacccatgatttatctcaacacatgaaattagacgcttctgctttatctgcgctcaacttgttaccaaccc

caggtgacattggaggaaaaaacagtagtctgtatggtttgctaaacaggtgtaaaacacctcaaggtcagcggctttta

agggtttggctgaagcagcctttggtcaacaagcatgagattggtgggtttcatatctgtcttctcagcaaagaacggat

aagaagagagcaagactgaatgaaaagactttgtctctctgtgccttgtatctctctttggcgctaaaagagagacggca

gagcttggtcgaaactttcgttgaagatgcagaagcaaggcgcacgcttcaggtaagtctacttgtttttcccttcatcc

ttatgatcgaacttatctgacgacgtcttctggtgaccctctttgtcaacaccactaacttgaattcaatccattagtct

gtgtatctcaaagtgatgcctgatttctctcgcctaagcaaacgacttcagcgtagaatggcctctctggaagacgtagt

cagaatctatcaggctgtactcagagtaagccagcacaatttcatctttgaacacatacttatgatcgtatcaccagaga

tagactaagctttctgttattcagttgccggagctcattgatacactggaaggtattcagcttgataaagaagactgcaa

gataattatcaacgaagtgtttgtcaatcaacttaaggttagaatttcgtgtctccacctcatctcttccctactcgcat

tttatttaccaaaccttctctcgtattgtcaatcttcattacggcttccttcaggaatattcgaccgctcttgagggtta

tattgagctggttgaatctaccatcgatttgaacgagctaggatctcacaactacgttatcaagtctgacttcgatgaca

ctctacaatcaatcaaggatcaactgattcaagctagggatggactggactccgaacatgaacgagtcggtagagaatta

gggctagatatcgagaaaaaattacatttggaaaatcaacaggtttatggatactgtctaagagttaccaaggcggtcag

tatgaatccatctttcccttcttttgaccggtgtcaaatcgttgataccaaactttgcgttcatctggatttaaaatctc

ttaggaagcttctgtcattcgaaacaagagagggtatattgagcttggtactcagaaatctgggacatacttcacaactt

cagcgctgaaggagttcgcgtcatcgtaccaggatttgtcccggtcatatagtaaggcgcaagacagcttggttaaagag

gttgttgaaatcgctagtcagtactccagtcttgcttctttcgagcgttgtttcacaaaacagaacatcaacagaaacct

atcagtagctgatcaagagttgtaattgggtttctcaggcacgtacatccccgtgtttgaaacgttagacactgttattg

ctaatctggacgtgattgtcaggtaagtcagatcttccagtgatgccctcgttgacgcttaagatcaaatcacgttccta

acctaatcttgttttttctgtagttttgcggatatatccattagcgctcctataccttacgtaaagcctataatcaagga

gatgggtcagtcttcgatgctacctctgtattttcattctcatgtacattcgaatgcggtcgtaactgagttctatcgat

caaacttcctttccactgttttcaggcagtggcaactgcatcatcaaagagggtcgacacccatgtctcgaagtccaaga

cgaagtcacttttattcccaatgacgttgaattcttgagaggtgcgcttcagcgttcttttatcgcaaataaggcggtcc

cattagacttgcaggataattgatatcgagaacccgttctgataatcgtgatgacagataccgccgaattccagattgtt

tctgggccaaacatgggaggaaaatccacatatcttcgacaagtgtgtcttctatctctgttgcgaatctcttccattct

ttcttgattctttatatttacgcttcgatctgctcactcttcaggtcgg

cgtcatcgcgttaatgtcgcaaatcggctgtttcgtccctgcgacggaagcagagatgccaatcttcgactgtgtattggcccgagttggagctggagacagtcagttgaagggaatctcgaccttcatggcggaaagtcagttgactcatctcttcattcagtattgtccgttcaaggactagtacgagattgtacttactctaatctcacttaatcaacctctcagtgttggagacggcgaccattcttcgtgtacgtgtcaaggctat

cattgtcgtatcatgtgattggccttattcataaacttgccctattccgatatagactgccacgagctcctcgttactgc

tgattgatgagttgggtcgtggaaccagtacatatgatggatttggcctggtaagtcactcagcaaattattcgcttctc

cgtcgatctaacaagtcctttctccttcacgctacacacaggcctgggctatcagcgagtgagctttgtttcatttcatt

tcatcatactatgtcgtttggcgctcctgatcacgaacctcaaatctattctctacagatatattgttacgaagattaaa

gcattctcgctcttcgctacacatttccatgaacttaccactttggcacaccaaaacccaacggtccgcaatctgcaggt

caaagcgaggatccaagataaagatattaccttcctctataccgtggagcctggtacgtaacctcgaattattcttgtta

cttgcgcgaatcatcgtctttgactatcatgatctgactaatctcttcatctttgtataggtgtaagcggacaaagttat

ggtatccatgtaagtagatgaacaatttgtcaatgaatggtcatcatatccgactgaccggtctctatcatggtgctcag

gttgctgagctagccaaatttccagaaagcgttgtcaaggtttgtcaacatcgatcgattaattggccgatattaccact

tgtctgagtctcttgatgtgtcatttgggaatgtagatggcaaagcgaaaagcagatgaactggaagacttcggaggtat

gttggcgtgaaccctttgtacccggggttactctatttcgttttgctgatctgtcgaaaccccatatttgtgatttctcg

atttgcagacgatgcggaagttactccagacgattcaaacccgttcaccaagtattctcgagaggaaacggaagaaggaa

cgaaaatcgtcaaacagttcctcaacgcttggtcttccagagtaccttcgtcatccaaagactcaggggttgatgagaag

agccaggcaacgttagaggaggaattggcgattctacgcgagatagcggcagaatacagagatcag

(**downstream homologous arm fragment**)

atcgaggccaacccttacactaggtcactccttgatagcttctaa

**>MLH1_ LN483166.1**

ggcgggtattaaagaaaacgaaggagaggattcaagagggagaaggagaacaagaagaacgaaaagacggtctagaataa

gagaaggatgaagga

tatgatgagaagcaagtcggtccggattgggccagatggttaaaacagatcacgttcgcccaaga

agagaaggccggttcgtcacctttgtggcccgtcagccagatattcatagcaacaacaccgagagatcaaacaccg

atggaatcagaacaggatacaccgttcgaaccaaagcctatcgtcgcgttggatgagatcgtgatcaaccggatagcagc

tggagaggtaaagccatgtatacgatatagggcagcagctgccgatgaagccatgtttctaacgcgtctgggtaaatgtg

attcctatagatcatccaacggccggcaaatgcactcaaagagctcatcgagaactgtcttgatgctggagcaaaatcca

taagagtctctgttaaagatggaggacttaaacttctacagatccaagatgatgggtgtggtatccgggtatatgcaatc

cctctatctgttctttccaggaaggaccgcatgtggactgatatcgagcatcgtgccctcttacctgacagcacgccgat

cttcccatcctctgtcaacggttcactacgtccaagctccgaacgttctccgacctcgaaacgattgcaacctatggatt

ccgaggcgaagctctagcatctatttctcacgttgctcatgtttctgtcctcacaaagacgaaagaaggcaattgcgcgt

ggaagtgagtgctcgtctccgcgatgctttctcctcgaatagatgtcaatgacacatgtttctgactatgtgaacggtgt

tgccggcctctatcatacagagcaaactattcggatggtgtactcatacctcttcgcccgggagagagcccagagccaaa

atcagcggccggaaatcagggaactactataatcgtgtgtttcagaattgttctttatctttgtagtctcagtagtttgg

tctcttctgacacattcggcgcgttctctctgtactttccaaacgtctataggttgaagatctgttttttaatgtgccaa

ctcgacgaaaggcgctgaaatcaacctcggacgagtttagcaggattttggacgtggtaaccaaatattcgatacacaac

ccttccgtctcatttatgtgcaaaaaggtcagctcaggtctttgtcttcaagagagttccgtcggcctgtcattttgttt

gtatgaagctctcaagttgctgatcttcccaatacatacctctgtatactgtgtaggctggccaggctgcccccgtactt

tctactccatgcctctccacttctgcacaaaccatcaggatagttcatggcccaactctggccaaggagctccttcatct

tccgaaaacaaagaacacccaatttgagtttgaatttgagggttggctgacaggcgccaattgg (**upstream homologous arm fragment**)

aataccaaacgcagtggagggtttttggtgttcataaatcgtaggtgatttcgaaatatccgcaatcttgaatatatgctcacttacagcgtgtaa

aatcttggaatcgggctgggtagatcgcttggtagactgtccaagactgaagaaagccgtggaagcgttatattcggtta

ttcttcccaaaggaacgtatccttgggtctatatagagtaagtcttagacgagtatcaattgatgcgctccatcagatca

ttctcaacgaatggtttaaatgtataagtagagtatctaacacggcctacgtagtctgctgattaagcctgagaatgtcg

atgttaatgtccatccgacgaagagtgaggttcgtgaggcattctacaatgtgtccttcagcatccttgttcttgctcca

gggccttacctgtttacatatcctgctttgtttcaggtgcattttcttaacgaggaggagataatcgaagctgtctgcga

aacagtccaggaggtcttgatcggagcaaattcatcgagaacatacagtactcaggttagcccggttctaattgaaatcc

accaataattgtccatgatagctaacaatatctctcttgataccccgaagcctcagaacaccgtgcaacagggttctgat

ccacaccacgattccgacccagagtttgaagaggaagaaccaagagaggtcaaaaatcttcctactttcggtactaacag

tggtacttatggagggagtaaaccaagttctactcgaatagtttcaagatgttagttgtccctcaatcttcaactccttt

tttgtgtgtgtgaaactcgatccagtctaattcattcccaatgtcgtcttacttatgattcgcatagcatccactcaacagtctaatcctgcgcataaagtccgggtagatccgaccgtccggacgctcgactcgatgtttgctcctctcaactcaaagatatctaccatcgtacctcccgagtctgctccgattaagcggcgtagaatagatgaacgcgttgagagtgaggaaggggatgaggatgagaacaaatcgaaggctaacattgaggatgacgcagatatgagtgaagatgaagtagagtgggtcaatgaggag

gtcaatacgcgcacgatagatgaaacaaaacagaaatcaaacagagaacgagggatcagtggaggaggcctagtgagaa

cgagcgagtgtaggttgacgagtatagtgaggcttagaaatctggtcaaagagaaaaggcataattgtaagttttttgtt

attcggtctttccaactttcatgctgaacagtcaattttgtccgattcctttgacatctaccatcggacatacatagtgg

tttcggaaatcatcaaggatcataaattcgtaggaatcgtgagttgtgaacgaactcaatctttgatacaacacaatacc

aaactatacttggtcaatcacgctgctatcgcgtaagttgtcattcctgtcgaaacatactttgcgctagggctgaacat

gcatcaagtttcccatcatactctttctgctatgtaataccatgaagagatgagctgttctaccagctgggtctctgtca

gtttggggagatcggaaagattcgcctggatccacctgcggagatcaaacctctgttgaagttagcgatcgttctagagg

aagatgtgcccgaggagcgaagggaggctaccattcaggttcgtttgacgtgttcaatttcaaacatcgagagagccctc

ttcctcgttgtcactgacacatgagattctgtaaacagatgattgaacagcgattgatcgaacgtcgagagatgctggac

gaatacttttctttaacgatcacggaggacggtaggattgagtcgattcctctgctcctagagggttatacacctgactt

gaatcgattaccagaatttttgatgcgacttggacctggagtgagtgttgaaatcttttgttttgctaatctttgtctcg

cgtagttaggatttggagacttggactgagcgcgttccttcgggagtatttttcaggttgattggtcagatgaagaagca

tgttttgagtcgtttctccgggaactagcttacttttacatcccccttcctcttttaccatccgacgagacggatgctac

aagagacaaagagcgagagctgggtatgttctctcattatgtatcttctttcaatcgggccttcttcctatagtcgccgc

tttaaagaatatggactaaattttgcctggcatatacttcattaaagaatcgttccaaatcgaacacgtcatctttcccg

cgctccgacgacatctccgtccttcaaagtcattgctaaaagggggccatatcactcaagtggc

(**downstream homologous arm fragment**)

taacctaccagatctctacaaagtgtttgaaaggtgctgatacag

tttagatttgcagacatatcctccttttttgatctttcatcttagtacatatatatatccactcttactatatccaccaagttgtacacccccaaaacttaccacgggttcattcttaaattgaatcttcgaatcaatactggtttatttccaaatagtgatctcttatag

**>Ku70_ LN483332.1**

aacttcgatagctcggatctaaataaaatcatcgaaagatacgttgccgtacacagctatttgcacagcttcttgtacat

cgcgcgaagcgaacagatgctgtgcttggttacatcaaacgtcattcctgc

cagcatctttctcatagccagatagtcagttgagataatctcctccatctttgtagctttccaatctctagagcaaacatatc

atgtcctcgacaccgccgatagcaaaagcaacacgtcgtagggctcacaagttggcacgagcaaaccctccatcgctgtc

caggttcaaatcgttgatagcctggtcggtttgtgtacctctgagtgagtgatagacaatcggaagacttttgacgtttg

cctgtcttctatcacaacacaacacaacctctcacaccttctgcctaccgtgttcagtcgtctggacattctcttcctcg

acagctacgggagctatcaagaacggagtggctgccgtgacgtctggttcgtctgtctcctccgccggacatccgaccat

gccgcccattgcgccatctccggtcacccatcaatcgaacccaccggtgacgctgtcggtcgaacccgccaaagacaaat

gggatgatctggtagaattcatagcccatgagaagcgccttttaaccagtcgtcaacactttaacacggagcagaagaaa

gtcgaacctatagaagctgctctgacagagggttcagcggaagatacagccaagaccgcctcgaaacgtaagtgtctagc

tttttcgtctctcttgacacatatcactcaaaccgacgtctcttttaaccttccctagagatactacttgctgcccttct

tggtctcggcagcttttttctctccctgttgagactgctctatatcatattgacatatattctcctcccaatcaccctcc

caattggctacctctttcacctatccatcgtacttccttaccgcatatcatactcgataatagaagtgctttcacctcta

atcgcactcataggaggttgtgcagctttcggagtgatgctcggaggaggcgctgccggactcgttaagggttatggtag

gttggctttcggacccttaggtaaagtcgaccgaaagactgaaaacgaagatgggaagaaaaggaatcattaggatgatc

tagatcatatatatgtatacgtatgttgtaatcacaagagaaaactcatcatgtgggtacagagaccctatggacacact

tgctatgagagcccgttcgattttagatctcgacgcgtctttaaaacgcgtcgaggcgtatctccgttctcatctcgctt

atcatctttcgtatattggcctggccatcatctatacacctccggaaaacttgatcaccatcatcactggccagtgctgt

actggtaaagaaggaagagagacagatggccttacgatcggacgtctggatagatgaaggtgaacctgatgatatagaac

tcatggaagaggacgtatgtttcccgttcccctttctccactcccttcctctcattcccctcctgtcttgccagtcaggt

ctcactaaacaaaaccttccctaaaaaaaaacaacaacaacccacgaaccgcaagacagtacagggcctcttcaagagat

gctgtgctgatctgtatagatgcttcgtccacaatgcatgaacctcaatcgg (**upstream homologous arm fragment**)

accaagatcaagagaccaaagcagaacaatatcggccagaagcctcgttcttcgaaggctcgttgaggtgtgctatagagttaatgaaaggaaaagccttaacctcaccatcggatctagttggggtcgtgattttcaacacggtttgttaactgtctagctttgatctaatctgtagcaaacgaatg

ccgggtctgacgattatctcctgcctcggaacatttggctctgttcaaacaaagaggaaagaacctcgcgaggatctgga

tgatacagaacaacacatgaccgaccccaaagggatctatacgctcatcccaataggttcgctcgacctagagcctatca

aatacctgatcgatttatcagatcgtaaatctttaaactagtcttgaaatgacgagatcgatgtactgatgtcccggaaa

acttaactcgcatggcatcagatcaagaaagtaacccggacttcttccaggagacgttccctcctctttcttcagatcaa

ccatcagtttctccaggacaactctttagtcattgtaacggagcgttccgatcagcgtgagcaccaacatctttctctaa

ctcccttctgtttttttagcgcgcatctttctctctctgctgactttcatatccttaccgatgaacacaggaacatccgc

acaggatctagaagagtgttcttagtcacggacgatgacaacccgatgaagcatctagacgatccgaatcaggctcgtca

agctgtaatggccgccaagacgcatctgcttgatcttcagacgataggagctagcgttgtgccgtttttgcttgggagcg

aagaaaagttgttcgatgtggatttattctggcaggtacatttgatatctttcttctttcttgtcttttccctgttttct

agaatgtttgacattaatgttcgccaccttttcttctgtcttctccatctcagagcgtattatcaccaccggggatagaa

cccaatctgagtatagggtcaacaggtctgatcgaaaccgaattcatctcggcctctgaaggcctagtcgacttcagaga

gctcttggcacagatgacaattcgtcagactagcaaacgagcaatattttctctcccaatgatcattggttcaggcaaga

aagattctggagattcaagctcacccagggaggtcagaagacaactcgttatcggtatccaagggtatagtccccctttt

tcctagagaaagaaggtaatactacagccgagaatatgctcatcagagagggctttcttgcttggtatgaaaataaggta

tgcgcatatcgtccaggctaagaaatcagctacaaagatgacggacgcgtcaggcgaagtctgcttggaagcaattcctc

aaattgaatattacgactcgctagatacagaggttagtgctcaggttttcctcactcagtcatctgcgcctccactccaa

gtttcttgtccgttgtcaacaaacctcctccttcttatctcggataggtaaagaatgcatttgatccggagaaccttcgg

cagacgtttagacataataaactgggcgacccagcccctccacttcaacgagattatgcagcccagtcagcagcccagga

agactaccttgagcacaaaggcttcggtacaaaggagctcatcgaagagtccgaaaaggaaaaaaagtcagatatcgtcg

atagagggaaaggaaaacagtcagagtctgatcgtccggaggtttgttcttctatttgtgcccttctctattctcattct

gttctacgcattggtcctggctagagagtaactaattatcgtcttcatcacggtcttgctaccttctagcaaataacgta

ttcattggaagaaataagccaactcaagtctttaggtatatcacccggtaagttgcatatctaaatactttttcaagtga

acgtacgtgcctcaatagttcattggatctgtcccggcgaatgatacacatgaccagggataagtatcctagggtttgtg

agccgatcattcttgacatttgagatgaacatcaaaaacagctacttcattttcccgaacgagcatgtacgtgatcacca

cctcttatgtaaacgcgcatgtgatatccacttatcctgttccttatcattatcgttactgctatgttaattctatcttc

agacctatactgg

ctcccgaagaacatttgcctcgcttaacaagacattattagagaaggataagattggtctggggctt

tgtctcttccgaaagggctcaatgcccatcttttgcgctctaattccccaggtataaccttgttcaattcgactcgatta

cgatcgaggcttacacgtcctatttatcttctacctctctcttcacatctttcgcgagcataggaggaaacatttgacga

agacgaaagacaggacaaagcacctgggttccatctagtccctctgccgttcaaggacgatattagaggaccaccaagaa

agattgaggaggctcaggaaaacgcttcgtatgatggtgcgcgcagatctttctgttccctcgcaggtctcatactactt

tcttcaatgttcttggaataggctcaagtgtgttgttgttggatttgttcagcggatgagatgcacgttgaaactatgcg

aaacatcatcaggaaactccagaaggatgatcgatatctacccgaacagttcccaaaccctagtaaggtgtccaagctcc

ggtctgtcaactcatccttgttctgatcacctgattttcaacctatatatatcatatattcctagctctatccaggtata

atgcacatcttcaagcaatcgcattcgatgagccattcggcgaagaagaggctgagcggattcaggataagacagaacct

ttagcgaaagacatgattaaggttcgtcgtctatttcttgtttcgtaccccagtgatcaacatcaaaagcgaaagcgatc

aaaggaagtgaaactctccctcttcacgacgtctcggatctgatgtaccgcattcgttctgtatcgatattcactagcgg

gctgggaaactcatgaaacaattcaaggaagggctattcgaagacgaacgaatacttgagacggttgaaaagcctagaaa

aaggaaagttggaatcgaggatgacagtgacgacgccctcgtttcagctaagaaaaccaagagtgttagtcaattttctt

ctcgatcccatgttcgggtctttgatgggatccgataaatcggctggtttgatccgtgccgttgattcatttatcctttt

cccggtagcttgctcgggagatggacatagcttcggttcgtgaagaataccgagcgggaagactttcatcaacagtgaga

accacaatccgttctttgtgttccgtctgttttctttcttttctttcatcaaagggcttccttgactctgaaagttttta

tgaaaataaacaaacggggaaaaataaccacgttttcagtatcgagttatag (**downstream homologous arm fragment**)

atttggtatcatttttgaaatcaatttctgtctcgtaagtatcttcccctttgttcctctccgtcggctcccctttcaat

cgggagccgatctgatcagaacatatattaatctatctgctctttgatttctttgacgtctagatgcccaagcaagaacaaagccgacttagta

**>Ku80_LN483142.1**

gccttgattaggcatcaaggtttggatgcctgtgcctgtgcctgtgccttttgaccgttccttttgaccgtgcccctgat

cgtgcctgtgaccatcggtcgcggtccctcggtaaccattcagcctcaat

catccaatccgatccttcatacacacacatccacactctgcccagtcaatactccgttcactctttcaccccgttcctctc

atggcagacagaggaggatacaccacccatgtgtttgtcatcgattgctctcagccgatgggtgcactcgtccaagaccc

acaacagtcgctcccggaaaaacgttcgaatccgcatctgcactcagccgtcgtttccaagtttgagctcgccagagagg

ttgttatgagaaagatcgtgtctgtggtaagtaagattggctcactcaagcaagccatacgcacgcacacacacacacac

acacacacacatatatatatatgcgcatatacatgggatttatagattggatgtcccagaaactgatatatcggatttcc

atctcatgtatatgtgtgtgtatgcacccaaacgagatagattatgagagaacgaaagaccgaacacgtcatggtcgttg

ctacaggtgaccgtgagtaatgctctcgtttggtctcgccttgccatcgaccagcttattctgactgacagaaggtcgtt

gtcgttgtatgcgttggacatccgcgcacccacacacctacccactctccatcactcaggaaccaacaacccacaaactc

aaggatatgaaggtgtagacgttctcttccaaccggccaccgccgcgaaatggatggttgacaaggtcaaatccctcaag

cttggaaacggcacgaatgatggggatggcaagtttcttctcttcctgcactccttcagattcttgttctgatgtgcgcg

ctctctttctcctcttcctctccactctagtccttctcggactcaccctggcgttgagcgagttagacaggacggctgct

aggaacagatcaacctggtcatacgaatttttggccttgacagacggtaaattctccgtcaattcctcctttttcttgtt

cccctcgaaccatcctaacttttcgcctttgtcttttccctgtccgccacccagcttctcaaggcgctctggcggacgat

ctggttgaccatattgctaattgtctctccaagataaacgcgactgtgaacttttcgtgcgttcccaacattaatctttc

caacaaactcttctttagaattacaaaccctcgcctcagtctcatctcatccttctatatagagtctatggaacagatga

cccagaatgggaatttatcgctggaccgagttcgtttgagaagaccaagatgtttgtgtgtactcccaggctctctgcga

agaaaaagagagctaagccagctgacctttcgaccgggggttctctcttttaaattcaagattaccaacgagcaaatatt

gtctgggctcattgcacaaaatcccgaacgtatcatccgaacgtccctcgatcgaagtgtcctcgatgcccgcgcgccaa

gcaccaagacagtcaagccgatgatgaacagc (**upstream homologous arm fragment**)

atctcactcattatcggtgatccggatatggacgtcaaccatgaggccgtaaccagtcatctcgagtttaacgtcgtaattggaaaaaagactgctctcgtcaagcccccagggtcgaaaaagatagcggtccgacagagacccggagcgggagtgagattgggcggaagccaaatagtcactcaggactcacaccatgatgaccctgaacagaatcccaccgtcaagatcgaacaatctcaaacctctactttccccgccggatcatcctcccaggcagtccgatcggtttctcaatccggctcttcttcctcaacatacgacggatcattccagataacggctagcctgtatgatgtgatctccaa

cccgctcaaacctatcaaaaaatacatcgtcgaactccccagagaaactgaaactgacaccgaggctggcaccgagaccgagaaagattccgtcaaggtcgacagggcggttgaggatttgactagatcgcacaaattcggaaacacctggttaccaatc

tcgatcgaggattcagacataggaaggttcgagactaggaaggcgctctggctttaccggtttatgcccgctcataaggt

gtgttccactttttctgtttttgccgagtcctgaagtgagtgtcctgaacaggatgtcttcttcctgcttccttctccct

tcctctcccttttcatcctgcttccttctcccttcctctcccttttcatcctgcttctcctcatcctttccgctttagtt

cgaacgagaactagaaatgggcgaagtataccaagtctgggcgaatccgaaatcaacaaaatcacaaattgaaatttcca

gcttggtgaaagcgatgagaacgtatccaaagggtgaagtctacgctctgttcaggtacacgtaccaggaaaacgctgcg

cccaggatgatggtaggcaaaccggtctaccaggacggtcagcacggcttacaactcgttcaagtcccgttcgcggatga

ttatcggaggtactggtttgactctctgctgcatctaaaggacacagacggaaatatcgtaaaggaacacccgtatctgc

ccacggatgagcagaacaacgcgatgttggatctttctctttccatgcgtttctttccccccccccccccttt

ttcctggtggactgcatatctgaaaacgcccctttcttgttgtcttggttcgttttctttggcacaaaacgtcctgaaaa

caacatcagagagagccagccatggtaccatccttctaaagtctacaacccaggactacatcgagtaaaagaagcgatgt

atcatgaattcactgcctcccccgatcaaccgatatgcgcttcccatccaagcctaacga

gtcatctgtcaactccgagacagctggttcaagcagcggcaaagaccgtccagactgctgtcgacgtgtttgacgtccataaaggtatgacagtccttctctttctctgtcttgttctcctgtcttccaaccatgtcagaggatcaccagtttgacagaatttccttctatggttgaaca

gtcgttcgtgtaaagagaagaaacccagatgaagaagacgaagaatctatagagtatgtttcccctatctttctccatct

tatccttcgcgcgacatcatctctttctctttgtattcgaaaaaaattcagactgatgtttaatttgatgaatcttattt

cacgtatagtttgggagaagggttagatgttttgattaagaacagagcagccgcaagagcaaacgcttcttcgtcgatgg

ccaacgcaaaggcaggcgaaacgtttgcacaatccactctggatcccatccctccgtttctcccgctcgagcgactctca

gggagacgtcgtccggttggttcttccgaggtgtctatcaagaaacagcgcgggtcgacagccgggcctgatctctcgcc

cgatagaactatcaagattcccacaacggcagtgggagatgaagcaagcgatggttccatcacccaggatgaggacgaggatgaagacatggatgaagaggatggaagggaagaagaccgggaggatcccatcagcgaatttgataagctgatcaagcgtgatggagataatactgtttttggtatgttccttttattcatctttctgtctctactccccatcagatccaatcaagctttgttcaccttcgtcttctaatctttggaccgtgaggacaacagcattggagcgtctacagaaggatatgttcagactcgttcaagcaggatccaacgaacgcgcgttggattgtttgttgaaactcagaaaagaatccctagaggtttgtcttttcttgttcatgttgctacaaagagtccaggaaaggaatcgcgaggagagaacggacgcgtgttgacgtaatgttcgagtggtttcttctcctgatgtcggttggattctagggcgaagaatccccccgattcaatgggttcgtctccccgttctctctacctctctt

ggctcgaacctaacctgcgcgatatcttatcttttccttcttcacagattcattcgttcactccggtccacatgtctctc

atcaagcacggcggctacgaatccgaacttgcccggtttttggacgttcttcctatccgaagcgaggtcgagagtgaacc

aagggcaaggagaggagattgggttgatctccgtcaacgaggattcagaattcagattgagtgatgtc (**downstream homologous arm fragment**)

tcggaagaagaacgtgtacaggtgtgttctggttccgttctggttgcctcttgagcccttttatccttcggtgattcactcaaaccgacaat

tttttgtcattcgtctctagttcttgaagtga

**>Mre11_LN483166**

gaatagtgacaaatttaactctgttattaaggtctgtgatcgtatggtatatgtatggcctataaacctaggaagagtgt

gtgagcagaaggaatgatgagagatagaaatcaaaagcgtgtgttgatcgagacgcgccagccgacagacgcatttgaga

cgcgtagcctcatttaggacgcgtcgcgcagtaacgctgacgctcgtccccctttagaccgccattgtccgatctctcgt

tcagatccacgccgtgtccaggtagcagttgtatagacacatgtcgtcc

ccaccagtatcatctcagctatcagagggagaaccgccgtgagtctatgtggctgcaagggtaaagagtgg

agagactagaagggaaagaagacatgggctgatgtcgtcttcgtttctgagagaacagacttacaagaccaactcctttg

ctaagggatatagaggacgatgggtcaaacactatacgcatcatgcttgctaccgacaaccatttagggtaccttgagaa

ggatcccattcggggaaatgattcgtttgatgcctttagagagattctagagctggccagaaaagaggaggtgggttgga

ataatctctccttatatccgtccatcttcaatggcacatagccagctgatgtctcttctttttgatggcgacaggtagat

ttcatcttgctggcaggcgatctcttccacgagaactatccttcaagatacactctccatcagacgattgctcttctcag

agagtactctcaaggggataggcccgtccagatagaacttctgagcgatccgaatgaagggaaagccgcaggattcacgt

aagtctcttagccaaatatcgacattattcttctgcgccttctctatatacgtgctgatgtctatcaaaaatcattcggc

ctgttagatttccagcgattaattacgaagactataacctcaacgtcggcattcctgtcttctctattcatggcaatcat

gatgatccccagggaacgtcagctgtgcgctcctcattcatatcatacccataccaggcatcccctgagttttaacaagc

gttttttttttttgcttctcaaactatccaggcaggagctctttccgcgattgatattctgtccgtctctggtcaactga

actactttgggaagactgatctgggagcagacgaagcgcgtcccgaatcagccaagcagggtttgacgatctcacccgtc

ctactaagaaaggggacgaccaagcttgcgatgtatggaatcggaaacatcaaggatgttaggatgagtcatgagttacg

gaacggaagagttaggatgttgagaccagaagaggatatcgataactggttcaacattgtactggtgcatcaaaacaggt

ttgtcctagttcatctttccttctactttgtcggatattgacgggtgttacctcaattttttttctttgtcaacacaatc

agagtggctcataaccggcatgagtatatacctgagaacatgtttgatgattcgacagatcttgtcgtctggggtcacga

gcatgattgccggatcgttccagaaatggtagcagacaaacaatactggatcacacagccaggctcgagtgtggccacta

gtttagcagacggtgaagccattgacaagtacgttctctcgttccttccgacttgctatcataccctgctgccgcgctag

acaaccagagtcgagactgatcgcttctctttctggtttacatgcaacatgatagacatgtcggacttctggagatccaa

ggtaaacatttccagatcactccgatgcctctcaagaccgttcggccgttcgtgatgaaacatatcgagctgtcagtcgt

agcggaagaaaccgggctgaatttagaaaataagccaaaggtaactgagtatttgagac (**upstream homologous arm fragment**)

aacaggtacgcctcgttttctttttctttacatcgacaatcagtcgtatgggtacgagtttgacgtccatgtctgaatgttttcctctggtctcacccctcttcatatttaggtattgaagtgtgtggccgaggcgaatgctaaatgggatgagaagaacttgtccctatcacctgaggtagacggtc

ccaagagacctaagccattaattcgtttgaaagttcgtctaaccgtccttctttgacgatttacctatgtttg

gatattcccgatccttatttttgactcttctacacaggttgaaacaacgggcgtattagagctaacgaacgtcacccgtt

tcggagaaggcctcaaccagctggttgccaacccaggaaatcttctacagttttatcgacggaagaagtttgccaaacgg

acagataaagtttctgtcgatcaacctgatctggacctctccgacgatgacgggctcgcgcccggtcagcgacaagggaa

gatcaagatggcggatttggtaaaggagtatttgacagctcagaagctagacgtcttacccgagaacggattagaggatg

ctgtcgaaaagtttgtagaaaagggcagtaagacggcgatcaaagagtttgtcctcatagatctcttgcctgatcgatca

gataacaaaggaagcttatctcctctttgtgcgcacttaaagttttgtcgctggaacgctgaaagcattcaactcaaacg

cacgaatgaagaacctgactgaggatgagatccaagatgaggtaccgtggactggttactcaatgtttcttataggcagc

ttgatctcattcatcgcctgtttgtttgccttatttttcaagttgatccaacaaaaaaacttggccgagagccaattcgg

agggaagcggcccgctgtgctatccgaagggaaagtcagagttcgtccaaacacctggtttcctttgcctttgttctttg

atctatcgacggtatggacattgatatttgtttaataaggcgtcgaatgctccagactcggacgtaatgatggcagagtc

gatgggtgaagactcagacgactccgtgttcactcgggcgacaggaagagccaaaaaagcaacgacctcaactgcagcaa

aggccaagaaagtcccagccaccaagaagcctccagctgcagcaaagaagaagcaggctttagtacgttggtgtagccat

cctgaacacggatatcaattttctgaacttaactcctctgctatctctagttcaacgacagtgaagaggaaattttcata

cacgatacggacgaagatgactgtcgaactgaaactcaacggacaaaaggtcatatcgatgatttcgatgatgacgagga

tggggatgaggacgaagagattactattgcttctcctcctagacgaagagcagcccctcagacgtccgttcctgcctctt

tcacgctgtatctatctctctcttcttctttctccaattttgacaacatacccttcttctctttctttctcttgatcttg

attttcaaatattgattgaacggttgtaggtcaagatcgagagcgccgccaaaggcggcggcgtcggcaaatacctcgag

aaggacaaagaggacggtagcagacacttctggttctcgttctcaacaaagtcaactcaagtctgtctttccttaccttc

ttctatttttgcatctatcctggttttcgtttgataagaaacggacactaatgttctgatgatcattaaatggaaactga

agtctggaaccagtcccaactagac (**downstream homologous arm fragment**)

cggctcgtccctctcgtaaaaatgtctgtcccgccctctgctctcttcaccttctttctttcgtttcgctctaattttgcatcattctttttaattcttgatctgatgactgacgagagatcttgttctcttctcttctccgctttcttttctctctgatgtctatacttgttgggttcaggaatga

**>Rad50_LN483157.1**

catcgtcgataccgtcctctccatccagagttgctggtacacatcatgttcgtgtttagccaagttttactgtattcaga

gacagatggataggcagatgaaagaacgcgctgctcatcttggactctttctggctcgcacacacacgcatttcatcaat

cacatcttcattaactctgctaggtcgtccctggatcgtcttgtcatcagaggaatcaggagttttgagtatgtccgaaa

atcaagattgggactgtatctataaggctgaaaattcataaagactcacctttctttctcgttcgtag

taacaactcgtccgccatggtagagtttcaaactcctgtcaccgtcattgtgggcgctaatggaacgggcaagaccgtcagtattttcacat

aagccattacaccagcccaacttaggagcttgagtcttgggagtcaaacacttcgaattgtcttcatatcagttattgat

atcctgcgggttctgggggttgattcgaatagaccattatcgagtgcttgaagtatgcaacaacgggtgaacttcctccg

aacacaaaaggaggcgcatttgttcatgacccaaaggtaagtttttgacttccaaaaggaagaacaacattcagagatcg

gaaaaaatctcattcatgctctcttatttggacaaataaaagatggcaggagagaaagaagtcaaggcgtcggtgaaact

gagatttcaaaactcaaacagcgagcggatggttgtcactcgtaacctatcggtctccgtcaagaagaccgggttgagca

tgaagacactagaagggattttatccaaagatgaggcaggcgaatcgaatcaaaaagtatattattttcactctttacct

cctggaagaagagttgcgctgaatgttggggtgtgactcggttcaagtacagagaaacacgatatctacgaagtgtagcg

agatcgatgacgaagtaccacttttgctaggtgtttccagagcgatactggataacgtcatcttctgtcaccaagaggag

tcaaactggccgttgagtgagcccgcgacactgaagaagaagttcgatgagattttcgaagcgacgaagtgagtggatgt

atttattcccctaatcctcttaaagctcattgcgcattgatcagggtgattgactgatcggcatacatgttggctgattt

tggtgatggacctccaggtttacggatgcgcttgattcgatcaaaaagatcagaaaggatagggcagctgatctcaaggt

agagaaagaaaagctcaaccatttaaaaatggagaaagatcgatccgaaattgttcgtcatccgattctctggtcctcta

atccctcttctatcaacacatgaacaccgagttgtctttcatctcctctagcttcgaacaaaattagcaaaggtcactca

atatatcaaagacaaagaccaacaagctgaggatctag (**upstream homologous arm fragment**)

gccaaaaagcccaggaaatggcggttatcaatatcagcttcagagaccaggccgaacagtttgcagagacgttcaacaaacaggagaacctgaagcatcagatctcggtctgggaggaacagatcatgaattatgaggagacaatgagtttattgcaaggttagctccggtattcttctcttctacccaaatgtctttcacacagctattgagagactcttaaagctgacgagtgtgttacatcgaacatggcatatcaattcacagaaagcgaagaagagcttagagggcagcttcatggatttgatcaaacgcttgctgataagaaacagctccgaacagcacgattgaatcggagagga

gacgaagaggatgtactgcgagcacttgaaagagatttcgcgaatcttgctcaaaagaggggaaggcttgatgcagaatt

tactgtttgttcagtctttctttggtccaatatccctatatgggtacgtggaaattggtactgatgagttcttctctttt

gagagataggctcaccaaacggcactctcaacccaagaaaatttggcggcaaaaatctgcaccaaattcgatatcgggta

ttcagacgacactcccttggtaagtcctactaatggtaccatccttaacttgacaacagatatatatcactttgctaact

ccggttttcttatgtctcccttatcaggaccaagcaaaactggacgagctgaagaatactttacagaacctaaaaagaga

acaggaacgaatactcaagaacaagagggtacgcttccacgtctatgagagagatcttctttctagtcatcctcatgtcc

gagccttagtgccaagtttactcatcgtaggttggcttctcaacgtgacttgtaacgttgacaggaggaaacccaggcgg

aagaagctcggatgcagaccgagttgataaacttgaataatgtgaaagctgccgaatcaggcagtagaaatagctttaag

agtcaaatagtcagttcatgtcttcagtttctgaggtggtcccgtctctttgcgcacccatcatgacactgatcgtatcg

atctgctttgcctctcaccattaggatgagcttgagaatcaggttcgaaaagatgccagaggggtcactcaaactagctc

ctctgtcgaagctgaactctcagctctccaagcggacatagccactcagcaaacaagcttggatgaagttgagcttgggt

tgaagactgcaaactatgaagaaaccaagcgaaggatcagaggagaaatcaatgctcttgataatcgccggaatttaatg

acggatgagctgagtgaacttcagcatcattcggccactcagacaaagttggcgatcaagaaggaagggctgatgcaaaa

ggaggcggaagcagcatcaatgtgagtagatcttctggcaaagctatcctctgcttatcaaggaaattctgctagatacg

gttcagccactgattttattcaatactggtggtatacagcacaacggaaaatactgaaaagtttcaaagattggtaggga

aaagtatcgttcagcgaacgatgattcgagatgttgttaaagctctatcgttggtctcttcactgttttgcacctctttt

tgcgagcagatctaaattgtttcatactctttggttgtgcaacagggataaagacggtgaagtggcagacgccgaacgag

acagatcgaaagctgacagagtgttcgctcaagctgaagcagccgttcagcgggacaaggacaagctattagacgcaaag

catgaactgtctggtgggtcatcactataaatcgaacttcctatcagagacttgacttcgagttgtctttttgggtctga

acaaacagatctaaaagacaaaattgatgcggctatacaggaggttgatggagctgaggatctaacttctggaatcaaga

tctgcttggaacagattgagcttgcacgagggtatgtgatattactatctataccaccataatcacatcttgaacagatc

acagatggatcgacctcggttttttaataactgttcaaatgttgtcatagggatatcgcatatgctggacaacttcctgc

actcttcgagatggttgaaaaatctatcaacagcaaacacgtatgcttaggttgtaatcgatctgtggacgatcgagaca

tgaagaccatactggcctacgtaagtcccatttcatggaagaaacgttttaacattctgctttgcttttgttgcgcttgc

ggctcaatgccacttttcatatcttgggtttgtttgaattcaaaacttcttaggtccacaaacagaagactaggagatcg

gtcgaagactccacagaaaagtaccaagaggacctagatggatggaccgatcaattagggaagctcaacgagttgaagga

gccagaaagacggatacatgctctagaggatgaagagttggagaggctgacagagcagatcagagaatctgagaagcgaa

gggaacaaaccaagtcagatctagaaaaagtacccctgagttctactcattcctttgttattcgcaacttttatttattt

ccacaacaaatgatactgatgtccttcgcccttccgattctcaggcccttgcgtctcttgaatcactcagagagaaagag

actcaactccaagagctccgggaagtttccaaccaactcgccagacttctagaagaaatttctagcttaaagcagggcgt

agaagaattggatcgaagtttgacagccagtggatcaacgagacctgtcgtcgatgttgaggatgacctgcagaaggcaa

atatcgacctgtttgtatttttctcattaacttaatcatctttccagtcgcataccaagctgaattgaattcatttccct

ttcccgttttcaaaagaaagagactccgtcaagaactccagcagatcagcgtggacgaggaccacttatcgaaacaaggt

cagggataccgtaatcagctcaacaatcttaacctccgtttggttactcttgaagcagatcttcgacaagttcagacagt

tgaaaaacgacttgtcacatctcgtgagaagatagacgatctcaggcaacaacttgcagtaagtcaacctagtacattct

atattgacaagtcttatataacttaataatttcatgtaatctttaccataggatgtcgaccacaaggtcgctgctgcaga

tgtacccattcagcgacttcaggaagaatttacaaagttccgaagtagcttcacgaatgagctgtccacgatggaggagc

gtttagcatcacttgtccaagccatcgcgaatcttgatcaatcatcagaggcggttgaagagtacgtcctcatcacacat

gcttcagtgatagttatctggtcatatcatttatccactctgtctgtcttttaggtggacagcaagcgaaggtcctcgtc

gtttgtcagatctgaatgatcagatctcgagaaaggagggggatctgaagcggtctcgaactttaatcactcaccatgat

gaggtcatcaagcagttcgagtctgaaatctatcagtccgaccgctcccgaaagaacatcgaagataacattcgtcatcg

tcaattcaaggatcgagttaggaacgccaagatagagatacaaggactcggtttggaggaagcctcaagagctagagatc

agtatgctaggaagtacaagaaggcaaaggagcaagagcagcagttgacctctcaagtaagtcttgagttttttaggtag

catacaatcaaatctaatttttttggattttctttctgtcagcacgcgactctattaggagaaatctcttctcttcgagc

gcaggccttggagatgaaggacactctgaacacagagtacaagaacatcaatccagaatttatgagcaaacttatcgaag

tcaaagtacgttctagtgtacatgtcctcaaataaaacatcttatcgtagaccgatctttcctaatttactgtacctttt

cgtttagacttccgatcttgccaacatcgatctagaaaaatacggcaaagcgctggataggtattcaattttgctatgtc

taatcgtcattgggcaatctgctgacttctatcgattgtacagtgctatcatgagataccatagtgtcaagatggcggag

atcaatgacaacatccgatatttatggaacaggacttatcaaggaacaggtaacgatcacccgtcatgttgaaaaaatca

tccggatatatcagctgaaggcttcgtgtttggtacatccttcagatatcgatcaaattctcatcagaagtgactctgaa

gaacctaagacggctacgaccaggaaaagttataactatcgagtatgtcattattttcagtctccggcgtatcagaccat

gtattcaaaaggacttgcattgcatttaggtcgtcatggtcaaggatcaggtcgagatggatatgcgcggtcgttgtagt

gctggtcagaaagttctagcttcgatcatcatccgactagcactctcagattcatttggccacaattgcggaatcattgc

cttagacgagtaaatcccttcaattatgtcttcgtcacatgttgagaagttggctgattcgtatgttgatatctttaggc

ccacaaccaatttggacaaagtaagtaaagggaactcacctcgtccatacgacaaatcaactaacttttcattctccact

caggaaaacatccgatcgctagctgaatcattagctcagtaagcttctgctctgtttggcttcttgaatcagtcatcttg

tctaatccaatactgcctgttttctcaagaattattgaggagagacgatcgagtggaaacttcca (**downstream homologous arm fragment**)

actaattgtcatcacgtaggagcttttgatgatcacttcttcctttatcaagcgctttgaagctcatgtattctcttccacttcaatgttcgttt

ctagtcacgatgaggagttccttcgggaaattgcatttgtaggtaacatttcacatcattgataacgctaaagagaaata

cctgctaatctgcatatcgttttaggcggagtaagcaccattggcaccttttgcccctttatatagaagcttgttcctga

tccaacgattcaatcgcccttccttccttcataaagcggtatcgattattatatgtatgtgttgaattagtttctgacaa

**>Sae2_LN483142.1**

ccatttaccgcattcctctccgatttctgtcttgtcctttcctgacatgtcaatcaatcgacccccaccttctttatctc

tctcactaaatggttctagtaaccaactactaaaagaactactggcttgtttgtcagatctctcatgaact

atggaagggtctctgatagaaaatctattaaaggctagacaacaagctgctgccgatcatgtcggtgtattggaaggggt

gatagagacgtgggaatcggagtatgccggtatgtctcaacttccttacaacctgatcacgaagaatcaaataggttcac

ttgttgttccgtgatccaattctgatttcattcgaaaaatgatggtgactggcttgatgtggcgtcttgtgatagtcata

aagcaagaggtaagttcgagtcctgtcaagcgcatatagtcctttccgtcaattaatgcacatatgtcccttttgacgtc

tcggtcggtcatacacagctaaaactgaaagaagacaagcttcagagatgcattgctcagaaggacaatcttcaaaatca

actagagactgtgcagaatcatgtattgtccctcgagaatagtgtaaaccgtccagtcgactctacagcttttacctcca

cttcatcgtcccctagtatcgatatgttgaatcagcttcagactcaacttgaaaccgaaatcgcacttcggagaaaatac

cagaagaggttgatggactccacccgtcatgtagacgccttattggaattgatcagtgaaaaagatcaatggcgagagag

ggaagcatcgatggtcaagagggttgataccttggagagtgagattagaactatagaagaccacggcagaacctttgaag

cagagaggagtaaatcagatcaacagtcaggtgggctagaatatcaggcgtgagtctctccgctcctttttgttgatgtg

tgtcggaaaatttatacttcataccagatgattagcttgcgtgcatgcatgctcatacctttctctcggttctttttcgg

atcccccccccaaaaaaaagaaccagccctggccggtctgatcttccgaaacctgtctccactgccgaggttcctctatt

gtcctcttcatcaaacaatacatcacctgttactctctcttcagagtatatctcgcttcaagcttcttatcgctctcttc

aggcacaacatgaacaacttcaggaaaagtatagatcaagaaagcaagaatggacgagttttgcgacgaggtacattcaa

gatctcacaaaatttcgggctgcggagaaatctagaggaaggaaccgtttggcaatgaaagagggcgctgacgtgaaagg

taaaagaagagaagtagtgaaggtgcaagcagacaacggcgcatcggcgggttttttaaggaaggccaagggcacgagtc

aagattcct (**upstream homologous arm fragment**)

tttcaagcaaatttggcgttgcaacaaagacgtcgagcgctcctaaggtcaatacaccaatgtccttcaca

cacaaccactctcccccatctttaatatcatcggcaaccgtcacaccaaatcttttaccaacatcagcagtgaaaaagga

gctcctagatactccttcactcacgttcaggaaatcaccacagcagaacaaaccttcagaatcatcggcccagcactcag

gtctttcagaaattcatgtatttcactcgaaacaaatccatcctgggacatttgtgtcggcagccattcatgatcggaaa

aggattactccgtatctcacccctgacaccgttcgtcacaaagtgaggcagagcgctggagatcatatgtctgatgtgca

tccggagaaaggtattcactcgaggcctaatatcacttcgatcggaacgaaagaatccaaacgcacatcaatgccactcg

ttgacacgtcaaaagatacaggtgttccttcttcagcatgcactacaaaatggctggggaagcctacacttcagcctgtt

cttcatgaccagaatgttcagcccgtaca

tctgactggtctagacagaggtgccaaccaggtcaagagatcattatcatc

aactgatcagatgatcgtgacctctactccttcaaaccgggctgataattgggaccatcagagccggaaaaggaaatcac

cgaatcaactagagcatgaacaacacagagtcacggtcaaaaatgaactagatgtagatgagttgcccgagatctgttat

cagccattagaggccgatgtacccaatcctgatatatctcgaacagtggactattcatcagactcagagggcaaagcttc

tacgaaggcgggacctgaaattactgaagcagtctcccgctcgggttccgtctttcgacaatctgatgattcgggtagcg

actggatcgatacgacatataagagtcgtccggtagatcctacggctggaatgacacccgacgaaaagaggatattcttg

aaatcattgagaaacaagaacccgaaggaagtgactgctctcttcgccggcttcaaaggtaacgggaggtatgccttggc

cgattcctctacgtaagtcgcacttctaatcttctgccttctcttccctcgagccacaagaaatgactgacctttaccat

tctgactcgacatccagtacgccatcgaaaactatcaatgaggagtttgagataaacaaagataacaatcaaggcgtcaa

ttatggtttgtctctttcttaagttttcttcaagattattacgataacaataaacctgcacgcttgctcatgacatccat

atgcatatagctttcaacagggttgttcgtaagaaggacgaacgtaaacaacttcatgcgacggattgcgaatgttgttc

cggcgtatgttattcccatttaattgtatctgttcctatccatggaaactaagctaagacctcatgctttatctgatctc

tactatatgatttagtactatgaagcggtaggcgcgctaccacctggtcctcaaggccctcgatggaaaagtccggctcc

gacatccagctctaaagagacaagggaatcttcagatacatacaaatttggccagagaggagaatccttagaagagagaa

ctgagaggctaaagcgggaatctgtacagatgagaaagcaacaaaattcgagacatcgagctgattgggaagaaccgcct

accccaccgggttattggtacattccgactccgattctgatcatcaacatctaattctgtaaatacatctgctgatttct

ttttgatcaaaactaccccttcgtaggaacatcggatttccaagctccccagagatcgccgctcaa (**downstream homologous arm fragment**)

aaccaagaggccgaacggttagctcaggcaaaacgtgcgcggattgaagcagaagcctcgtga

**>Rev1_LN483167**

agactacatgaagaacaaacggatgaagctacagttacagaacagagagatcgctcagactaggcaggatggagtgcctc

agatctttaaaggtctggctatatatattaatggctatgtccagccttctcttcaagagctcagagatatgtttctctct

catgggggagagtatcacgcctatttagataaaaagagcttagtgacccatatcgttgcgaccaatctgacgcctgccaa

gatcaaagagtttcaacaccggaaagtcgtgacacccagttggatcgttgaatcgactcaggctggaaaactacttcctt

ggtcgaattatcgctggaggcctgtacccctagcaacatctacgtctaactcccgacctactcaaccaagccaactgact

actcaacggagtattcgatccttttctggtcaatactacgagccagtcaaacaacccctcaaaccgctgataactgagcc

atctcttatgtctccatcgcctgattctatcactgcaaacttatccgctaaacacaccctcacagttctcaaaaaacaat

cttcgtcctcgttaaatcgcagctcatcctctttgatattggcgaagacggatatcagtcctcctcggacgactccctcc

tctttcgtaagaaaagagaagacaagggtgtctcaggaaatcatacgacccgtaacggacctggaggttgaggaggagtt

gcttgtcatggattctttcgagcattctagagactcacaagaggagcacgccaaggaaggagactcacttaagggaaagg

ggaagaatctacgacgcaagaaggagaaagggaagaaaaagaaacgcgctgatcctttctcttcctcttcttcatcctca

tcctcagcttcggatgaagaagctgacaagagaacgaagagaagcatgagtcaccgtctttcgtcccagcgggttcgtct

ggagatgactgctagaagtcttactcgctctcagagagccgccgtccaaccaaacccatacgatgatgactttctagatc

agcccacatctaaattaaaaccttcatctcgcccgacagtggctgaccggtctttacgtctttatcaatcattatcaacc

gatcgcctccctctggatatgccccaaacctccctggtgaagacgcgtgaccagcccacctctactcgtgatcgagctag

agaccagaatcgttcccaactttcaccacgcacaactctcgaattaagcaatgatgatcctcttcttctggtgccttcca

ataagacatcgatatcaactcagccaaagacctatcgctctctcaagactgcgcgtataccgtctactctcataattccc

tcagatcctcgtcctacgagtctcaaagagaactctaaaacaaccagtagagagaggtcagagcgggaggagggaaaggc

aaagagaaaaaagaggacgaggagagtcagatttgaagatgaagtggatataggtatgaagcggacagatctcacggggg

cccttaagtcaccaccaccactcttgcaatcaacacctttaggggcgacaaagtcccggtgcttagaggatcttgagatt

gagaaggaggaaaggagggctctcatgatgttagacaagatgagaaggatcgagtcagaccttatctctgaagagaaaca

gaacgatgggcgggcagtctcttctttcaaagcttcatcgacttctctccctactttagtttctccctcccctctctccc

atacttctcatgctgagcagtccacggttccttcccggacttcgatagaatccactgaatctccagcccaacttccaact

actcaccctcagcctgttctggtcggaatcgctcccgccgagaaggtagtgtcgataagtcacgttgataccctgaaatc

tcccaagagtgcgcttgtagtca (**upstream homologous arm fragment**)

attccatgactagatatgccttgcatgaggagcatcctcaggcaagagagttaatgc

aagattcagcgtggagaagagggcatacagctcagagcgaaacatttttggaaggatactatgggaaatccaggctccat

catctttctaattggaaagctgaattgaaagttcttgtcgctcaggcccggtgtgactctcgctcgctccccgtttcgcg

agttttagctcctcattctttggccgattcttttcttcctcgacctgcttctctcactacctctgtcaagcgatccggaa

gaggagagaggatcatcatgcattgcgactttgacgctttctttgtttctgtcggtttggcgggagaggagaggaaggcg

ttgagaggtagaccagtggtggtctgtcattcgctgacaggtggaagacagagtacaagtgagattgcgagtcctagcta

cgaggctagggcgtttggaataaaagctgggatgagtcttggacaggccaaaaggttgtgtcctgaagtgaagagtattc

cttacgagtttgagaaatacaaatctacctctctaaaattctacaccatacttctatcttacgccgacagtttcttccaa

gcggtatcgatcgacgaagccctgatggaagtgacctctcgggtcaatgatcttgctaagaatggcactgagagggctgg

aagagaattggctgaaagaattagagaagatattaggcaagctactggttgtgaagtctcgatcggggtagcttctaaca

cgctattggcccggctagctactcgaagagctaagcctgcaggatcctttcatctgatccaggacagagtatcagagttt

atggagagtcttgatgttgccgatctatgggggattggtagagaaaccaagctcaagatcgaagctgctttcaaaacgac

ccgagttggagaactcttgaagaagcgcctcggagaatttcagcatattctgggtcccaaaacaggagagaccgtgtaca

acgcctgtcgaggaattgac

tcgaagcctttgaaggaagaatcagaaagaagaagtgtcagcgccgaagttaattacggc

gttcgatttgagaagaattcccaagcagaagactttttgcagaacttatcggtagaagtctcaaacaggatgaagaaagt

tgatcgggctggacggatgataacactgaagatcatgaagagagctgaaggagccccggttgaggcgcccaagtttatgg

gtcatggaaaatgcgacgtcgtgaacaagagcaaagaactagtgggttctagaggaggcgcgacggacgaccctgtcgcc

attggaaaagccgctgtcactttgctaaagctgatggacataccagcatctgaacttcggggcattggaattatgattac

gaagcttgaggacccagtagtcatttcaaatctcaagaaggctggtcaaaccactttaactttccaaaagccagacgttg

cctcagtaccaaagtcggcgcaaacttctatcctaccaccaccatcaccaccaacttctgcggctcccgccgtagttgct

cctcccgttgacctaccgacctgtctggatcagtcgatcctggatgttgcccttgtagactcttctacatcagctcctac

caatgcagatcgtcaatcgatccatgcgatactcacaaatactccacgaaccacgaagaaaccctcccctccacctggaa

cccctctgaatttgctcccagcgtactctcaagtcgataaagaaacccttgcacaactcccttcatctatgatccgagag

ctttatcctcagtggcagtctacgcaatcgaagccgtcatactctcagcctaatgatgtgcctagtccatctattcagct

tgtagattctccaccacccctcaggaatccatctcccgtcaaaaaatcgcgtgtagacgtctctcatattgccaagcagc

ttcgaccttcaaagaaatctcctatgcggaaccctcctgcttgcttcgcgccgcctgtagcaggacgatctttacctgct

atagaatcgctccaagcgaatgcgtctagaaagcttgcgcccaaagggtctctagaggagattactcaactggccgaact

actaggttgggatctgaatttcatcaagtccttgcctccggctgacctgaaagaaattttggatgaagggaggaggcaac

gcgatgttcgagatcgattggtcagaaaaggatcaggcgacaaatccaacccgacttcagccttcccaccccgcgtcact

tctcttacgccagaaaaagccttgcacgatagaccgggtaaccaattgatcacaatgatacctacgagatcgatggcgac (**downstream homologous arm fragment**)

gatgtcgagggacaaccatacgactgatcgatggaaaggcgatccaccagcaccgattcttgggaagaagaataaactag

acccggaggttctagcccaaggccaatcgattgattgtctgcgcgagatcatctcgtcttgggtagatactgagggagcg

actgggccttcgcttccaggcgttcagatggtaggtgattttgtggtggcttgtgtgaagacgaagggaaaaggtcataa

**>Rev3_LN483116**

catccgtattgacccgtgcctcaccatctctcatgcagtcacctgagatgtcttcaggatcgcaaatgaccaacccaatt

cgtagtccagagatatcgagagatgccgatgaaaaagaaagcgatcctaaatcgacgcctactatgtggtctcttcccag

aacgcgacctacaataactcctctcgaaattcaatcaacgacatctcggcagatcgtaaatgttccaagttctatttata

aaccagcatccgggaatacaacagcagagaaaatcgcacggtcttcagcggcggacgactctttttttgatgcatcactc

gatgcaggggatgactggattttctctcagaccgggccggattctgtattgtcttctctgcgggccgatcggtcggtcca

gcaggtatccaacgatagttcttctccagttaaacaattgcttatgagtcgtgtcgaga

ctcagtcagatgcagcgacaaataatccgcttggtgggccctatggcttatcatctagttcctattcaatacggagagctcaacataagagtaatcgagagaatgccaaagacgcgcccttaagcctacccactgatacttctgaagcgacgatagaccagccagtccttaacccttcagt

cattcgtcagtcggatcagttgtcttttttgcctttttgtctcggtttgagttaacacgtttggctatctaatgattgca

tttgaatagcatcggtcgactcgatcatctcctcctcagctcagctcaagacatcggcgtttatcccttcaccttcgcaa

caatctaaacaatctctggcatctcccacagaatatggctgtgatattgattatcaagatccctacttctccaatccagt

cgacgttccaaagtttcctagagagtatgccggaagactctggacgtttgataaaaacggtaattctatccgtcctgaag

aggtctggtcgccgaacgtgttcgatggcattcgagatcaacccgaaatcggaggtagcagagtctttaaatggggcggt

cttaagggaggtctttcaagattcagacgaatggtcgtgggagaagtagcatctttggaactaggtcatcgcgcaccatc

gaggagtgaggctaaggagtggctaaagaaggatattcaagcaaaagagaatcgtgggtttatctacttcttggtatcct

gttctctttttctactcttttatcttttatcgagatcaatattattactttgtgtctatgttatcttgttgtatccaaaa

agtttctcacaaactacttttttaatcaatttttaatagggagaaaaaagattttcgcatctcaggtatgtctccatgtt

taatactcattccaacaattcttggggggtttcgaccagcgattgtttgctaaccgtgtatttgttttttttttgctggg

gaaatgttctgatcataagatagaaggacctactcagttgaatactcaaggatttaagttcagtcagattaatgaagtct

ctagctctcgggagaagaggttcatgagtgttatgtctctagaagtccttcgttagtcttaatgtgttacctcttttctt

tgaccatgaagctaacttatgcttaaacaaactggctctgctttacatcctctcttctcacaatcggtttatagctgcaa

caaggaatggccatctaccagatgccagagaagatcctattctcgctgtttttttctgctttcaaagcgatgggcaagct

ttcgacaactgttatatgccaggatgccatgtcgggctcatcatcattatgacagacgaccattctagcaatcacagtc

(**upstream homologous arm fragment**)

aaccaccttcgacagatcccaggcgcactataggatcgcgtccgatagagcctgagtttgtgtctagcgaattggaccttt

taaatttattgactgagaaggtctgggcttgggacccagacatcttgagtggctgggaagtacagggagcatcttgggga

tatgctacgaaacgaatgtcgctttatggaggtgatatctctccgatatttcttattataacacttctggagtgatgaac

tgacagcacttactccatggtacatgctctttgggttctctttgaacaggcgaacagttctttatggcggagctttctcc

tgtactggatacagacggccaaaacacatcgtctgaatatgactccgatcatagctccgccttccagatccaagggaggc

atgtctttaatctatggaggatgcttcgagcagaactgaacctcaatcaatacacattcgaaaacgtggcttttcatctt

cttcatcgtcggtaagcagcttcctttcgaccatctgtatcttcagggtgtggtcttttgtctgaagctcttctcgtcat

gtggcatgcagagttcccaaattctctccacaaagtctccaagattggttctgctcggacataccgtccaaagtgattcg

gtgtatcaatcactttcttcttcgagctgtgatgattatcgagatgctagatttatcggaacttgtttctcgaaatgcgt

gagtggtttttcaaaatcatccggattgacattccatatgatgggagattttacagattgacatggacccaatttctgca

aggacagagagttcgcacgggtatatggcgttgacttctcggacgtttatcaaagagggtctcaattcaaagtagagtct

ttcatgttcagaatagctaagcctgagagtctgttgttccgtacacctactcgagaacaggtcagtgttttgtcaacgtg

gagtcgatatggtatgtacgtcttcggacaggcactgatgcccttagtacttctttctttacaggtcggccaacaagccg

gggcggaatgtattcctatggtgatggagcctcaatcagccttctacaaggacccagtagtagtccttgactttcagtcc

ctctatccatcaattatgcttgcgtataatatctgctattcgacttgtctcggacgcgccgaccgactgaaggataaaga

tgggaatccgactgacaggtttggctttaccaacttagagctctcacctgggacgcttgaggtactcgaggatcacatca

ctagtaagtttggctgtgtagacctccttcttaaagatattgatagattgatttcaaactgaaatatgctattatacaat

gctctgtgttggtagtttctcctaacgggatgatgtttgtgaaaccaactgtgagaaagagtttacttgcgaagatgctt

ggcgagatattagatacccgggtgatggtcaaggacgctatgaagggatataagagcaacaaggtgatcattgttacgta

tccatctacgctgcgacagtagtgatataggctgatcagtgtgagtttactgccatacagagcttgacgagaacattaaa

ctctagacaacttgcgttaaagcttttagccgtaagactgttttatcgagattgtctactgcctcgattttgacttcaag

tttgacatgcattgttttgaatctgccgcttttttcaaaccttcagaatgtcacttatggctatacttccgctagttttt

cgggtcgaatgccccatattcaggtggcggactcaattgtgcagtacggtcgcgagtccctcgaaaaagtttgtctttta

catgtctatacttgaataaccaatgctggaggtctcatatctctcttactgcag

gcgatttctctcattcactcgacccctgaatggggcgcacatgtcgtgtatggagataccgatagtttgtttgtccatcttccgggaaagagcaaagagatggctttccggatcggatatgagatggctgatgcgatcacggctaagaacccaaaaccaatgaaattgaaatttgaaaaggtttgc

gttctatcaactttttctctggtatcaataagtatagtgctgacgaacttctgtgttcataatttcgaccatgacaggtt

tatcttccctgcgtgttacagtccaagaagagatacgttgggttcaaatacgagcatcccgacgaacttgtcccgacatt

tgatgacaaaggtatcgagaccgtaagacgagatgggatacctgctcagcaaaaggaactcaaagaatgtataaagtgag

ttttgtttgatttctaacatactgcacattaggttattcagaagcgtgtcttgctaacatacttgagactaacgaaactt

tgtcggtctattttgaacaggttgttgttcagagatccagacctctcaagagtcaaagaatattgttatcgtcagtggac

tagaatacttcaaggccgagtttccgttcaggacttcactttcgccaaagaagtccgtttgggatcttataggtaagttc

agaatacggttgtttgtttcatgtgtttgctgttcagttttctcatcgcttccttgctcgttcttctttctttcttttat

aatcaacatcaatctaaacagcgaaaaaagtcatccgcctcatgtagtggttgctgcgattcgagcgaatcaagatcctg

gtgacgagcctcagtacggagagcgtgtaccctacgtgattgtccaaggggaacctggagagcctcagtatagacgagcg

cgtcgaccggaagatgcgttggaggacgagtgagtcgtttgtctcctgtgacccgaccgtactggcgccttaatagactt

gatttttttgatggctgattttacttcgtttttttctttcaccgatccacagaggactgcgactagatggtgaatattac

atacaaaatatgctaatcccgcctcttggccgagttttcaacctattaggagcggacgtagcgggttggtaccgaacgat

gcctcgtccgaagtggattgagcctgctccttttggtataccttcttcttcggctgttactactgctgcttctgctgtt

(**downstream homologous arm fragment**)

ggtccggacccattaggtcctgcgcagatgattgagagcgctagagggaaagttgttgctcgaaagattaagttgacagac

cacttcaagagtgatctttgttttttgtgcaaagctgaaacccatctaggtaataatttcctatctttcaggcaaagagc

ttttcgtttagtgagcaaccctgaatagttgctgaaacgtttgccttctctctctgcattgttcgtgatcgcggcttata

gtccccgcgatatgtaaatcgtgtcgatccgatccttacaccacgatctacgctcttcagaaccatcgacaacaagccga

**>Pol4_LN483249**

tctcaaggccatcactcgccattgtttggatgtgtacaaagcacgaccagaacagaacaggaaaggagagaaggagataa

actcaaggtaaatgacgaacaggggctccagaatgatagatacatgacccaggctgattgttctctgggtcgaacgaaag

accacatcccggccggttgcagctcagtgttaaggacatctgtttttcaaaagctagaggtgcccggagtgcctgtcagc

gtccttgtacttgtattaacacttgactttcgactgactagatggatagaaggatgaattgaattgcaagtagtgggtgg

atagaaagaagaagaagaaggaaagaaacaaattcaggaaactacaaaaatctctaatactttttttggatctttgtaat

gaatcttgatcgaattttctatgatgttcgtttgttccttctacgtccgacgctcatcctaagaaagaccagacacagca

cacaaactcacacaaacgcacactgaaaagactcaaaaactgacgagactgattagtctgtcgattgagatgcctaatat

tctaggttatttgaagatctatataatccctacaaaatat

tctccatctgatatccggcaactctcggccgaactcaccattcatggcggtcagctcatagaccaccccgaacgagcagatatcattctgactaggctgatgggcagaaagcggctggcgttagcactggatccgacgttgattgtgaatacttcttgatatccagatcaggcagtgtctcacgagtgtagtttgttcagatactagctaaacctttgaacgttttcgtctctccacattctcttctgacgcgcaggactccaagatcgttgttcttgat

cgatggctcgccgattgcatagcccacgaagccctgctggaccatcgaccttacctggtcccccccgatcattcctcgta

agttacagagtgacttgttggatcagtcgcttcctcttccgtatctgtgacgttgcgctcacgttgtttggtcgtttggt

ttctctctccctactcccatctgtttttattttattttattttagagaatttttgattcctcccttgatgtcttctttct

ctgatcagatagaacccttttcttttgatccttgctttcttcctccaaccccagatagccagcctactcacaagtggcct

cggaaacaaacagcctctccttctactcgacctcccaatcaacataataacacacgttctccatctgatcataatccctt

caagacaacctctcctgcactcattctcgcttcattcccagcgacggaaaaatctatcgaaccatccactctcccgagac

tggcatgctgtcgaaaatccccgtcaatatgtatcaatcaacttttgattgaggaactcgacgtcatccgtgtttggaaa

gagaacggggaagacgacacgctagaaagtgatcttaatcaagcaggaaagaaagatgacgagaaacatgcattggcgta

cagtagagcgtgctccgctctaaaggcttacccaacaaaaatttgctcagtcagcgaggcccaggtgattc

(**upstream homologous arm fragment**)

ctttcattggtcccaaaatagctttacaggttcgtccgtcgatgtgtctgagcaaagaatgatattaaaaaacagagtcaggctgacac

gctgccttcagatcggagagtatctttcaaccggacagatcaaactctcacgtgtgttctcgaacattttcatcgatcat

ttacgcgtacaaaacacaattgaccacttgatgttcatatatgtacatgtccgaaatggaccgtcgcatttgtcttcaat

cagaaacactccggacatctgttcgacttcggtcaatccttctcttccaaactcttcacggtgtaggcagtcggctctct

cgagagttctatgacatccatgattgtcggacgctggaagatgtggcaaacgcgagaccatccttgaaggtccaagtgac

ctactggtcagacttacagacgccgtgagtttcatgggtgctgattgatgactgttt

gtacgtatatgcgaggcatactcgatcgaagatctgatctaggataattgaatgttgattttagaatccctcgatatgaggttccattgatcgcaaagtttctctcggatgaactcgaaatgatttctcctggttgtatacacaccatctgtggaagctaccgtagaggccgagaatattcaa

atgatgtcgacatcgtgtttacctatcctgcgatgaacaggtcctccaaggtatcaagccaggtgctcgatcagttggtt

gatcgattgagaacgcttagtgcgtggcctcttgtcgtttcgtcttagtctcatcgttcttctcattgaatcactaacat

atggcctatcttaggcgcattgacggacattttgtcggggagttcaattgggtcgaatgcgctacgagggcttcatcaac

gcttcatcatgatgagcctacccagacaaaccaggcaaattcgtgtggacatcatctttgcgcgtacgtcagtttcatta

cttatttcgccatgaacaaaaatgagagacagctcatccgagtcatcccataccatgcgttgacttgaactttatgtgac

tgactgtgttccgtgtcgccttgggtcgagcagcatacgagatatattgggtctgtgtggttgggtggactggatcgaca

atgtttgaacgggatattcggtgagtcttgcctctgcttgcttgcttgcttgcttgctcgtatcacatgtagaactgatc

tgaacatgtatctcttttctttcaacaattgttcaagaaaacacgccaacgggctcaatctcaaatttgcttctcatgga

atatctcgactatcagacggttcggagatcgccgtttctcaagaaaagggagaacctgctgtattcgaggcgttgggatt

agagtggattgagccaactgaacggaacgcagacgtgtag (**downstream homologous arm fragment**)

atatgagaaacgatatgtaaaaacaagtatcatggttgtagatgtactgtagatgttctatagatctgatgatacctgaagaatgatgctcatgtatagatctaacggcggttgattcagccaagaccgagataatttgatataaggcggtagcgtgctgctgattcttcagccaccaccaccaaaccctttcgtcgttcccactaacggtcatcatctttgtctcttcctttctctctgcccattcctccgttctcgttcttgaatcattgtcgagggt

taaatagtagaaaaatggacgtacaagtgtccggctatgtgccatacgtcgagccatcccttgtctcgattctctctttg

ggatccttcttaatcctcgccaacgtctttgggtatgtcatcgttctttatcatacgtttacgtcgtttatcagatgaac

tagcgatgaacctcttggagataattggtgctcatttctaggtcgcattgatattgttctgcaacaccggaatcatggta

**>RAD30_LN483249**

tggattaacgttctgtaaaggacctaaaggataaatcaacatggaggagagcaaggaccccaaaatgttgtgacgtgaca

aaccattaaacgactcccggatatgtgattcaaacaatgggagatcagcccaatcgaacttaacgccccatattgttgcc

tcagaagttattttttgctcagtaagagctaagagaagctcgaccgctcgtggctcgtgaccgattgcatttctcaacag

agaattccacaatgaatcgtgagaggaccacagaaatgattcgacagagaagaaagtagaggatgacgaagaaagaagag

tgttgaaatgagcggatatcttgttggctgtatgtggtgctgtatcagtaccttcaggacagaaataatctacaagaatc

gcgttgatgatgtgcatcccaactgttggtgatcgatgcctgtatatattatacctagcaggccttaggcttggtgagag

cggcctgaactgatggtggtcgatacatgcgccttttggactggggtaaatgctcatcaccaaaaccaatctgttcattt

aggagaatgacactggctgagtcatcaaacgcgtcgcaccacacgcgcaacaaacctcatttatcacgtcctcatacgcg

tcttcgaaccgtccgcttcctgttcgacaatacttccacgatggaaaatg

aggcgtatccttcagatcagccagtgacctataggcacctcctctccccccaaacacttggaccatcaaatcccctacga

gtcatcgctcattgcgacgtcgatgctgcttatgcccagttcgaagctaaacgcttagacctcgatcggagtgtacccat

tgccgtccagcaatggtctgcactgatagctgtaaactatgccgccagggcgtttggaataactcgtcatgaaaatgtgt

ttgaagctaagaagaaatgtcctgagttagtatgtgttcatgtggctacgctcaaggacggagacgaagagccgggctac

catgagaagcccaatatgaataccgacaaggtgagtctagaaccatatcggcgcgagagccagaagatcatcaagttatt

caatagggaggcccccagcggcgaagttgaaaaagcctccatcgacgaagcgttcctggatctgactgtccctatccgta

ccgtccttcttgagaaatatcccttcctttgtgctgtaccccccgattcacctcttggtttggatacgccccttccaccg

gcaccgactctgaaatggaaggagagaccatgggggatcgtacctgtgcagacagagctaggtaaggatgcttcagatgc

tgagacgcactcagaagcggaagccgacaaaagttgggcggacgttgccctctgggaaggaggagagatgcttgacagaa

tgagggaaacaattgagctcgaattgggctatacgacatcagcgggagtcgctcataataaagtcttagcaaaattgtgt

tctgctttcaaaaagccaagggcacaaaccattcttcgtacttctgccgtttctcaatttctaagacctatgaactttca

aaagatccggaacttgggagggaaactaggtgaagtaatcg (**upstream homologous arm fragment**)

ctgaaacatatggagcggctaccgtcgaggatcttctgaagatagaccttggcgatatgagaaagactttaggggaggac

tcgggttggatatacaatatcttgaggggaatcgactacagcgaagtcaaagctaaatcactcccaaagtccatgatggc

atccaagaacttgcgtccgtatctaacaaatatggagcatggatcaagctggttgcgcgtactctccgtagagctgatag

gcagattga

aagaagcgagagaactagctgggctcgatggcggggatggagccggaggagtatggccaaggacacttgttttttcttgg

agagaggctggaagcgcctccaaatccaaacagatcccctttccttatcacagtccgctctcagcctccggtcccattct

aacgttatcaatcaaacttttgtccgaagttcttaggcctaatctccgtctcacctctgatcgcaaacaagtccaaatca

acaatcttgccttaggatttactcaattagagagacttgagactggtcagagaggcatccagggtttttttgctatgcct

atccaggagactggaagcagaactggagagaagcgacgaatgtcaaatgagaacgaagaggatcggagtaaacgacctgg

atctggctggttccaaaaggccccgatgggctacaaagctcatttgggagctccaccatcgacctcatcatccttagcat

cctcctcttctctttcaacagcaacagacccctatctatcaattacaactccgctcgatagcgctgctgtcgacaaaatt

ataagaccaggaccgtctgttcttgcagatcttagtcaccgctcgaccctttcctcatcttcttcttcttattctttccc

atccgtgccgcgaccttcgacaaaacctatatctgcgtctgcctctgaccctggctcttcatgctcgctatcttgttcct

tctgggtgtgccctgaatgcaaaggacggatatcgctcgctcgagagatcatgattgatttcgatccagaggatgaagca

agggagagtgagcggatcaagtcggagcatgccgattggcatgttgctttgaaactccaggccgagttcgaaggcaatct

gccccatcagaacggtacagatgttggcaaagatgaaaaatcaatcatagaaagagaaggaaccgggttcggacggatgg

gaccaaagggactgccgaaggtagtaacagcatcatcagacccggatatgtcgaagaggaagtctggagaagttagtaag

ccgagtcggaagggaagcggggatcacgatcggagctcaacaggatcagagatcagaggcaa

(**downstream homologous arm fragment**)

aggtggattagggaaatggctgattaagaagtgatcttttgaacgaacgggatgacaactttggtcccgggactttggacttttcgaaagatggcctatgctgagtctgtgtagatggaatttcgaatttactcgaattatataaataataggcattcatctacatggtttttgaattgattgtatacagccagttctatctctttgtcaacgacggtaaatgggctcggtcccattctccctctaaacccatgggtccttgtctggcgcacca

cggaggtctgtaggacgagtcaccctgccaatattgtcggcatctgagagtgtaaccagtggggactgatgaagaggtga

atcgtattttatttcagagaaagaccaaatggattcaaggtgtgatcaatgatggtcaaaagtcaaatatcaggatagaa

**> tRNA-specific adenosine deaminase_ADA1_LN483124.1**

agttaaactactcaatccgttccgacctttcatagcaattctgcaagagtgcgcatctggtgggccactaacatccagga

caatattatcttgagcttgcgtttatcagatgccgaacaagcagtggtctagcttctacctggcttagatccaccaaggc

aatcggctggccacggcctgataacgcggccgatacgatgcgtaacgcactgcgcgtaaccgcgtaaatttgttgaagct

gtctaatctaatctgtgttcgcaacatttttgctgatgttccatgcttcttgaatcatatcctc

atgaccgacttgcccactagctatatcgacccggatatcgtagcatcgctcgtcgttgagacctatctatccttccctgc

gtcctgtgtaccatcaaagagatccaacggcgtccaagagtggacgcc

tcttgctggagtcgttctctctcgaacgaccgatagttctcccatcctcaagctcatatccgttggagcaggttccaaagttgtgcccacaattagaaagtctgctagaggagacattgtcaatgatctacatgcagagattctgtctatcagaggagcgaggcgttggattctggaggaagtcggtagatgtttgaatggagccgatgatgattggctcgaacagatcaccaaggaaggtagggtgaaatggaaattaaaaactggagttc

aggtccatatggttcgtcctgggtaccacatcacgttcccgtatccaggattcaggctgattgtccgtttgttcatcaca

gtacatctcgacccttccttgtaagtcgcacagctcgactgttcttctttcagtattccttgcgctcagttcatcgaatc

gttgtgttgtcctccctcaaaggtggggatgcctcgactgtacacctagccaatcaacaacaaatactcgacccagatat

ggctgctctcaaatcttctcaccatatgccctcagaaacatcttctgactctgctcctagcggcgtgaatgcaccgcgaa

agtcatttgtcctcgcacgaggacgagacgggtactccagccttggtgttattcggacaaaacctggtcgggcagactct

tctccaacaacgacactctcttgctcagacaagctagcggcatatactctactaggtatacaagggtccgtcttatcaac

agtcgtcgagcctatctacctaagc (**upstream homologous arm fragment**)

actatggtaattggtgatatcaaaggactggtggacaatgggttcgaaggagagaaaatcatacacgaagagttagacagggctcttgtgcagagggctagaaatacatgg

agtagccttcaatgtatgtcaaacatattcctctcagcgttctttgaataagggtgtatgtatcttattcgttcttgcgcttgtggcaacagccgaattaccaaatctatcatatcgtcatcaccctcctcgtctgaatctcacgaaaattacattttcctactcaaaatcccttctcctccga

gatacgccagatttggctatagctacaccttcaatatgtaagctttcatcatcgctttttccacatctgtcaatgtcatc

ttttaacaagttcttcatgtctacttgtgtataggtctcgtatacacagcggatgacaaaccagagcgtctggttgctct

gggacatctccatggatctttcaaaaagagcatacaagccgctgggggcaggctacctgacaaggctgtatccggcgttt

gcagaaaagtcttgtttgcgttgacgctggatattttgaacaaaacgcgtgatgagggctttgacattcggtgggtcatt

ccacaaccgaagcacacttctttaccatctatcacttgctttcaattgattacgttctcatcgatgtacaacctcaggca

atcataccgatccctcaagaagtctttcctggcattcgagtatcagcagaccaaacagatcctccggacgcatccgatgt

ctttcttgtgcggctgggaagaaggagacagagagcagtgggaatcctttgattcagctggagacatcgaatccctagat (**downstream homologous arm fragment**)

gttgagaaatcgacatcgagttctgtatcatccgaaacgcctgtagagaagcggcagcgactgaggtgatggagtttgacttttgatatctccgtccaacgttgtctcccgtttgtctttttagctatccgaactatttattttatatttttgaattattatcgtcgagcatgtggactaattttgtttttattttgttaagagcgccattctatatattatcctacaacctgctctcttgtacatcagctctattcattgattgcagatgtatcggtcctgaacagatggctcgatgtaacccaatgattgtggtttccactgtgattgtactcacgtgactacttctggtcacattgtcaacgacacgcgagttccaaccctgcactttctttc

**> Cytidine deaminase_CDA1_LN483157.1**

tacgaagaggccaatacgtaaggatcgagtgcagtataaatcagagtcaagatgagtgttcgttaggagatggttcggag

gagtcgggcaatgaggtattctctggcggatgctcttctccgatataaagctactttattt

gatctttcgtacgccgaccatccttatcttcatctcaacagccatgtcgcacacagttaccgtagaacaatatgaatccctcattcaagctgctcttcaaggttggtgatttgatggttctggcaccaagatgcctgctgagaacgacttgtcgcgggaatatttagacaacatgtattctcttcttttctatatcgatcactttacagccagagatggatcttactcaccttactcaaagtttagagtaggtgctgccttgctcaccgctgaaggagaaacc

atcaagggtgctaatgtcgaaaacgcatcgtatggtccgtcctatatctccttcatcatcttcttggctgtggccatgtt

atagaaccgacatcatatgtactgatgataatattgaactaatattgcccaacttaggggggacaatttgtgccgaaaga

tcggcgctttgcaaagctgttgtgcgtactcatgctgtctgttgttgtcactggc (**upstream homologous arm fragment**)

ggctccgaccatctcttgttgtttccaaaatgcc

ggagtttgatggagtagagactttgagctcacagatctgactattcaaccatcatctcaccctcttcgtcc

atcgtcaaagactgatggaaagaagaacttcttggcgatagcagttgctgcgtgagtctcttctatttcgttcactctcg

gtttccgaaagcttcatccaaatgtaataactatccggaatcttaatccatacagcgatgttccctcagcttctgtctca

ccttgtggaatctgccgtcaatttatcagagagttttgccccttgactgtttgttctaatcttgaataggttcttctttc

tattccgctgtggaacgagacctaagctgaatcgacgtcgacactctacccatcttcctgatacagactccgattgtgat

ggtttcctcgacgtatcaaactggagattacaccaacgatcaacaggctgtaaatgaaaagaataaaacaacggtgacga

tgacattagaacagttgctgccaatg (**downstream homologous arm fragment**)

agtttcgggcctgatgagctcaaaatgggtcagagatcctgaagctgtgtctgagatgggttgcaaggcagttccagttgagagagagagaagaaaagaggaaactgattggagagaccgcagcagcgaggaagagaaggaataagatagagacagagacagatgatggggttgtagagaccgttaatgcccagaaaaatgttttatacatctatactaaatacagcattcaccagaatctaaaatacatggtggaaaatagcaaagtat

**> Cytidine deaminase-like_CDA2L_LN483166.1**

caccgatcatcacattaggattatgaacgatatctattacaaaattttgtagtcgatgtttacacaacttgtagtctttt

ttccgctctctcacaacccgatacaaccatcaaccatcaaccatcaaccatcaaccatcaaccatcaaccatcaaccatc

aaccatcaaccatcaaccatcaaccatcaaccatcaacacccaaacgcatcgttaggag

atgtcgtcgaacagtagacatgtcgcgttctttaaagcgtgtgctcaacaggc

agctttatctcagatgacgttccgtttgggagcggtggtggtgaagaacaacaggatcatcggcagagggaagaacatgtccatcaggtaagctcttattcgttctttctcggcattcgtgcgatgaggagaacaaagagaaggaaaaggagaaggagaagagcaggaagctgaaggtaacaccaacgtcgttcaccaacttagatccactcatgaaggcccaatcagtcccacatctatcccaaacttgtcgctgcatgccgaagc

ttccgccctccgatcttgccttcacttcaacgcacgaactccattgttcgcctttccttcttacactcttcccgtcgtgt

cccactcaggccaaggtgatgatagcaagggctctgtatcgtctccctgcttctcaacggcagcgaacttgaagcatctg

cctcaagtcaagcaccttcatcatcaccgaaaacgatgctacactgatgcactcacaaaactctcgaacacgacggtgta

cgtcggtcggtggaacaacttgggcgaatgtttgagcgcaaaaccgtgctggcggtgtattcatctgatgctccagtatg

(**upstream homologous arm fragment**)

gtgtcaaacgagtattttggacggttccatcgaccactcaagatgcctctctcttgggtacagtttcgacatcgcttcac

aaggtcgcccgatcaaattccacatcttcaccgtcgtcgcc

atcatcttcatcgccgacacggaagcttccggccggaaa

ttgtcagcttcattggcaagaggggaaagtggaagatctgtggagagagattgtcgagcacgaaaacttgatcacaagcg

ggtacctgacgattgtagaagagcaagcttggagaaagggtcaagagagacataatcaaagagtcaagtctcaatgaagg

aatataacaaggaaggagattgattcttagccggtatgttcggtttggtttcattctttgtacaagacgttttcatatgt

ccaagtttaaactcaaattcgaccttgtatctgattcaaagtagccgtcgttgccgaccggtctcgcacaagccccctgg

accaatctaacttagcagcatctttactgaagttgactgtcaagccagatgaaggccatgtccttctttggtttactgca

accgtcgatagaggcacgaccgacgacgaaatcaaaatcgcgttccaagctcttcttcgccatgccctcgaaattgtcga

gagaaccagctggatctcgaaacgtcaactgtcagaaggaactcgacagtatacgaagcagtcagatcatgaatcgtcgg (**downstream homologous arm fragment**)

cttatttggtgaagaagatgaagaaggaaagcacaaacgctagatatcaggcatagatgcggtgaaaaagcacactctggattggcacgatcgcagaaggaatgggcgctcgaatatacgcgatttgtactctgagatgaacttgtctccataacatgtctccctgatacgtccatctgggtttccatggcaagaaaggaagcggcaagacggaaagcaaaggacacaaagagctgactggccggaggacaaatctgagtgacaacaggaggcga

**>Atrazine chlorohydrolase/guanine deaminase_GUD2_** **LN483142.1**

agaaaaggaactgagaacaagatggttgaagtcgcacaggtcgtatttcgcttaacaccgtaatcggatttgaaagtaag

ccgccaaaacaccaacttcattctcaccagcttcatcatcatcatcatcatcatctgacgactc

gtacatcgctagttcaatggaatctgaaatcaacaaagaatcctgctttctcggcacgttcatccacactccaactctcggctcgctcgagatact

agaagatcatcttctcctggccgactctcaaggctatatttcc

ttcttcggtcctcagtcgtccaaagccgctatcgaca

ttctcaaccgtcattttccagccggaaacaaacatgtgcttcctccgagttctttcttccttccttcattctctgatctg

catatacatgcccctcagtacctctacgcgggcacagggcttgatcttcctctcatggaatggctcgaacggtatgcgtt

cagagccgaacagcgcattgatggagacaaggaattggccgagagagtgtacgaacgattaggaaggcggatgattgagg

tcggaacaggagcagcgctggtatttggaacaatcaacaccgaatccaagtatgaaaccccaaccaattcttttgttctt

tcttagtagaaagtcttgaatactcaaactccttggcctttttctcttactctgattatgtctagcttaattttggcaag

gacgttcctcaagcttgggataagaggctatattggaaagcttagcatggatcaaaatagtccagcaacgtacacagaga

ccacagcatccaacgccctctctgctgcgtctaccttcattgattctatccgatcactaaatgagcccaacaatgcctta

tcagttcatccagtgatcacacct (**upstream homologous arm fragment**)

cgatttgtcccaacttgctcggatgaattactacaagggcttggtaagcttagaga

ggaaaaaggtgtgagggtcatgagtcatatgtgtgaggccagagatcaagtcgattgggtcaaagcaactcgggacgatg

tagaagacgttgagattttcaagcacgtacgttcttcgtcattcatccggttcttcataaaatacataaaaagtcttcca

ttatcccatttttttcttcttgcacttctgtagcagaatactgatttttaagctttgctttttgcttctaaatagactgg

gttgttggaatcatccgtacaagcgcactgcacctttctgacaccagagaacatcgagcttatttcccacacaaacacag

cggtcgctcactgccc

gctatcgaacgcgtacttctcttccaaaacatttcctctccgagagttggttcttgtgtttttt

tgtctctctatccgcgattcttctgagctaacaaacgttctcattctcgttgataacagatgcctcgataaaaatatcca

tgttggtctgggttcggacatcgccggtggttatcagcttgatctacaaacccagatgcgtcaagccgttgtcgcttccc

gtcttcgagactctcaactcaaagagaatgcacttggtcatccccgtggacaagaaaatgaaccgaaattaaaggatctg

agatcaagctggaaggaaagtctctatatggccactaggggtggagcgttggcgctcgcggacccaaacgtcggaggcag

gttcaaggtcggagaggcttttgacgctcagtcaagtacgacccatccaaatatctatctcacacccatggtgtcaacat

actgatccgcgtgtgacgtccatcagttgagcttctggatccacagacaaacgaagggacaggcgccctagacttcttcg

atgtggacgctgtcaagggcagcgtcagcctaggttgggaggagatgatcgagagatggtggtgtgtaggagatggtgcg

ctccctttctcttgtctcatcctgtact (**downstream homologous arm fragment**)

tttctctgaatttgaccacaaacgcaagctgatatccgttgttataaactatgtcaacatcaccagtcaggaacagaaaaggactttgggtacaaggtagacagcttttgtagatattcaatcgttcgatttgtttcatgtagtcacagttagacatcgtctttcgttgtataggaaaatatttttcaaagaa

aggcatctaattaaacaacagacccgacaaggatatatctaatttcgatgattcacaattgggcagatcgagatttcatctactgctctcttgatcaactctaatcaaagcgagacatgtttcctcctccccctccttc

**>Cytosine deaminase FCY1 and related enzymes_FCY1-2_LN483116.1**

tgcatcgttcaagtatactaacttcagtgacagatgccgatcaagaagttc

gctctttccgaatctgctgaaactgactatgtgttttactaaaacattcttcttcttcttgtctggtaacatatttactatgtctattttcgttcatttcttgaaacaaaggcctttgaggttataaccaaccgagcgatttcttccaggtattcatctgacagccaagagtctcatctgggaattttg

aacgaccgcgaggatcggatgtccgttcggctcgtcatttctcctccaaagttatcggacgtggacatcacacaggcttcaggtcatgtagagtccttgacgtcgatatcaacagcagagagaaggaaaaggccgtcaagaggtaaagtcaggaaagagggaagaagaccaaaggatg

tagaactcgaggtagaggtcaaacaggatatgaaacgattgaagggccatgctggcgggacgggttgtgttgtctggaga

acgaggtgagcatttgcgattttctctttgtcgtggtagccgactgaatgcacttaggaatcaaaccttctgttttgtca

gtctttatttcttggaatatctgcttcctcagttatacttctc (**upstream homologous arm fragment**)

ttctatgccttcactataccctccgctattta

acgtagaagcacttcaacaagcgaacgtgctcgagttagggtcgggtacgggtctcgttgctactttcttggcgaaattcgtcag

acgatgggtctgttcagaccagtaagtctctgtctttgacggttacttgtctcatgacaagtgagcacctgattaaggtt

attcactagtctgatatctcttcttcgtccgcgctagacttgactgcctcaaattgatctctcagaacgtatcttcgctt

gacccatcaaacaagacaaatatcgagattgccgaagtcgattggcttcactctacgcctcctcctaccgaagcatacga

tttgatcctttgcttggattgcgtttataaccctaatgtgataaaaggactcgtccagacattcgacagccaggcagagg

aaggcagaacggtcatttgtatagtgatggaggttcgagctgaggatgtcgtcggagagtttctcgagacttggttgcaa

aatgggaagtcggacggaaacaaagttcactggatcgtcaacagatttgattggggcgacggtgacggcgctggtggaaa

gtttgtcggctggct

tggctggaaggggaagcattgatgtacaacatgttaagagaaggatactgccatgcgttaattgcggttggcgcgatagacggtgattcagagcacacgtattagacaagatcaaccattcatacattgaaccctcaaactttcttttcatcatctcatactttgttacctacattttttga

tttctgttgcactatgctggatatattcttcccaaaaactgtttgagtctttccctgtctcgtaccaaccccgagctttgtgacg
